# Complete Reconstitution and Deorphanization of the 3 MDa NOCAP (NOCardiosis-Associated Polyketide) Synthase

**DOI:** 10.1101/2020.01.28.923383

**Authors:** Kai P. Yuet, Corey W. Liu, Stephen R. Lynch, James Kuo, Wesley Michaels, Robert B. Lee, Abigail E. McShane, Brian L. Zhong, Curt R. Fischer, Chaitan Khosla

**Affiliations:** Department of Chemistry, Stanford University, Stanford, CA 94305; Department of Structural Biology, Stanford University, Stanford, CA 94305; Department of Chemical Engineering, Stanford University, Stanford, CA 94305; Department of Bioengineering, Stanford University, Stanford, CA 94305; Stanford ChEM-H, Stanford University, Stanford, CA 94305

**Author notes:** Department of Systems Biology, Blavatnik Institute at Harvard Medical School, Boston, MA 02115 and Wyss Institute for Biologically Inspired Engineering, Harvard University, Boston, MA 02115. C.W.L. and S.R.L. contributed equally.

## Abstract

Several *Nocardia* strains associated with nocardiosis, a potentially life-threatening disease, house a nonamodular assembly-line polyketide synthase (PKS) that presumably synthesizes an unknown natural product. Here, we report the discovery and structure elucidation of the NOCAP (NOCardiosis-Associated Polyketide) aglycone by first fully reconstituting the NOCAP synthase *in vitro* from purified protein components followed by heterologous expression in *E. coli* and spectroscopic analysis of the purified products. The NOCAP aglycone has an unprecedented structure comprised of a substituted resorcylaldehyde headgroup linked to a 15-carbon tail that harbors two conjugated all-*trans* trienes separated by a stereogenic hydroxyl group. This report is the first example of reconstituting a *trans*-acyltransferase assembly-line PKS either *in vitro* or in *E. coli*, and of using these approaches to “deorphanize” a complete assembly-line PKS identified via genomic sequencing. With the NOCAP aglycone in hand, the stage is set for understanding how this PKS and associated tailoring enzymes confer an advantage to their native hosts during human *Nocardia* infections.

---

Within the past decade, genomic sequencing has exposed many “orphan” biosynthetic gene clusters encoding assembly-line PKSs whose products have yet to be identified^1^. Analysis of orphan PKSs has the potential to reveal new biosynthetic strategies as well as products with unprecedented structures and biological activities. Of particular interest to our laboratory is an intriguing family of orphan assembly-line PKSs termed NOCAP synthases. NOCAP synthases harbor *cis*- and *trans*-acyltransferases^2^, and are invariably found in strains of the actinomycete *Nocardia* isolated from patients affected with nocardiosis, a serious pulmonary or systemic disease^3–5^ (**Table S1**). The NOCAP synthase is composed of four separate proteins (NOCAP_PKS1-4) containing nine PKS modules (eight of which are collinear) (Figure 1). Modules 1 and 3 possess their own acyltransferase domains, whereas the remaining modules require a *trans*-acyltransferase (*t*AT) to supply malonyl extender units. Notably, this PKS has three other unusual features: a) a “split and stuttering” module capable of catalyzing three elongation and reductive cycles (module 5)^5,8,9^, b) a terminal thioester reductase (TR)^10^, and c) a thioesterase (TE) domain fused to the *t*AT. In a preliminary study, several unprecedented, albeit partially characterized, octaketide and heptaketide products were generated by incubating purified modules 4-8 with the surrogate primer unit octanoyl-CoA^5^. Building on our laboratory’s prior experience in functionally reconstituting the complete 6-deoxyerythronolide B synthase (DEBS) in *E. coli*^6^ and from purified protein components^7^, we sought to deorphanize a prototypical NOCAP synthase outside of its genetically difficult and potentially hazardous natural host using both of these approaches. Here we report on the successful reconstitution of the entire assembly-line NOCAP synthase *in vitro* as well as in *E. coli*.

**Figure 1.**
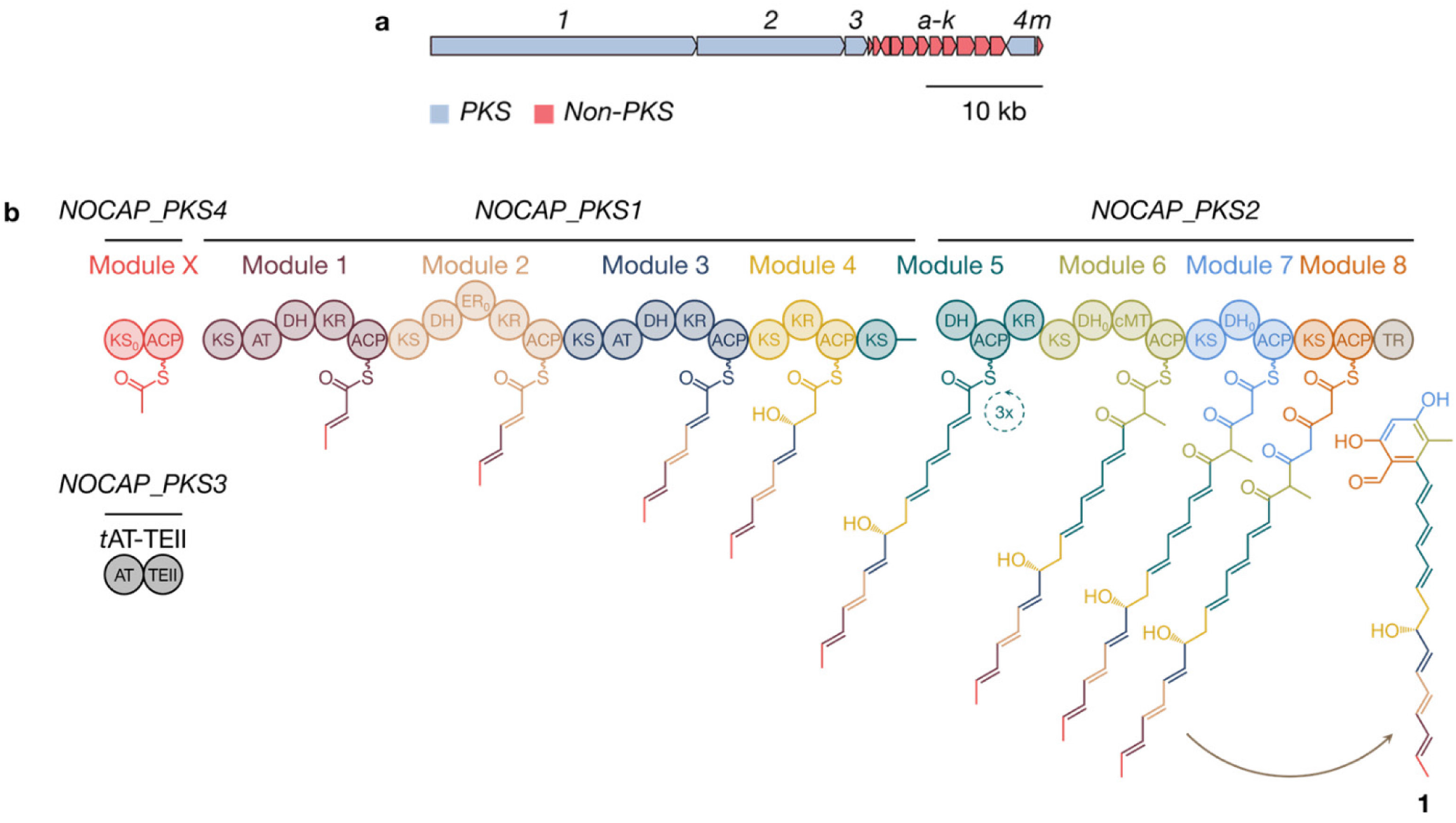
(a) Prototypical NOCAP synthase biosynthetic gene cluster from *N. puris*. (b) Biosynthesis of **1** by the NOCAP synthase. Key: KS, ketosynthase; AT, acyltransferase; DH, dehydratase; KR, ketoreductase; ER, enoylreductase; cMT, *C*-methyltransferase; ACP, acyl carrier protein; TR, thioester reductase; and TEII, thioesterase II.

We hypothesized that the uncharacterized module “X” synthesizes a primer unit for the collinear assembly line comprised of modules 1-8. Accordingly, we expressed a soluble maltose-binding protein (MBP)-module X fusion protein in *E. coli*, and purified it to homogeneity (**Figure S1**). To identify the substrates and products bound to its acyl carrier protein (ACP) domain by intact protein LCMS, we further expressed and purified two derivatives of this protein: MBP-module X without the ACP domain (KS_X_), and the stand-alone ACP_X_ (**Figure S2**). *Apo*-ACP_X_ was incubated with the Sfp phosphopantetheinyl transferase11 and malonyl-CoA to obtain malonyl-*S*-ACP_X_, which was then incubated with KS_X_. LC-MS analysis revealed that malonyl-*S*-ACP_X_ was predominantly decarboxylated to acetyl-*S*-ACP_X_ in a KS_X_-dependent manner (Figure 2a), suggesting that KS_X_ is condensation-incompetent (hence the designation KS_0_ from here onwards) but is able to decarboxylate malonyl-*S*-ACP_X_ to generate an acetyl unit for translocation to module 1. Interestingly, while KS_X_ appears functionally analogous to specialized KS_Q_ domains^12^, its active site Cys residue is not replaced by a highly conserved Gln. KS_0_ domains are prevalent in *trans*-AT PKSs, and lack the active site His residue located within the HGTGT motif^2^; however, KS_X_ also retains this conserved His residue.

**Figure 2.**
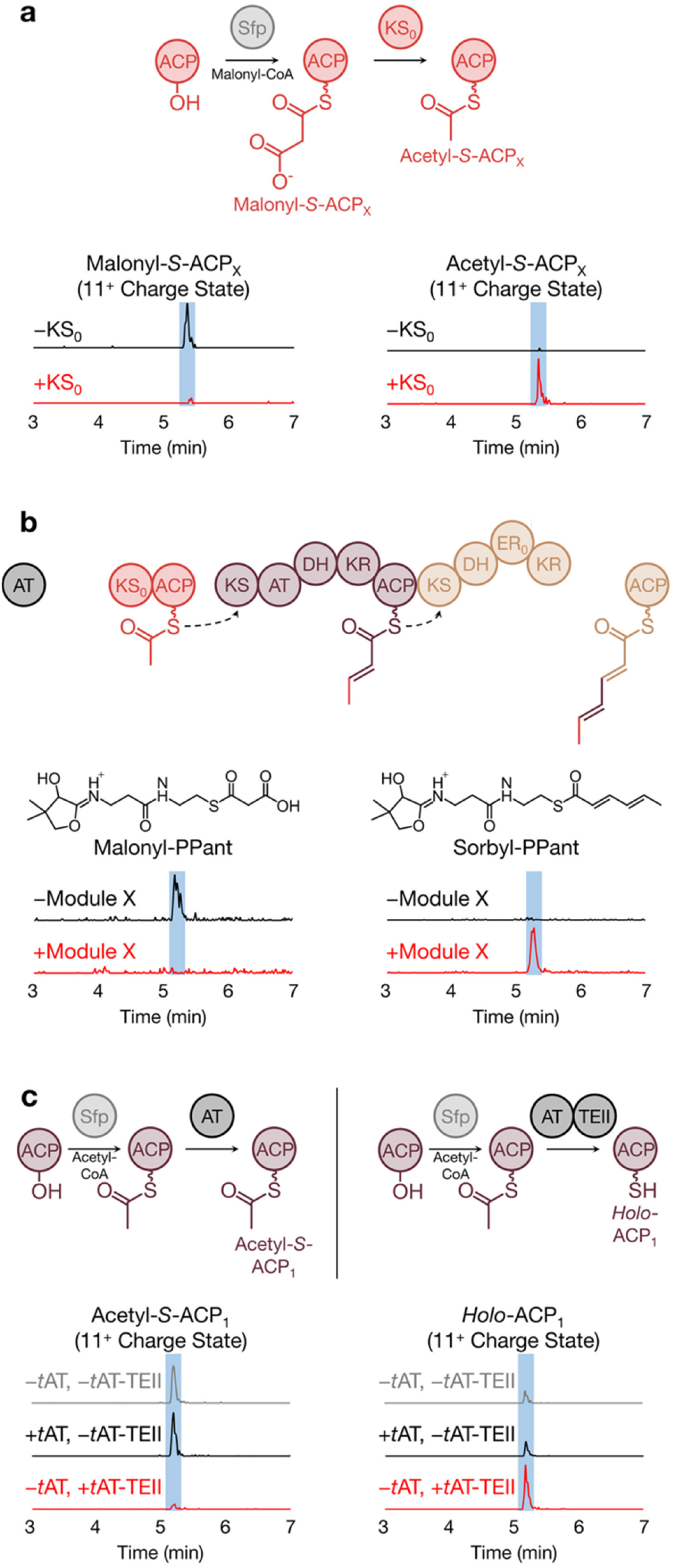
*In vitro* characterization of modules X, 1-2 and *t*ATTEII. (a) Extracted ion chromatograms (EICs) of the 11+ charge states for malonyl-*S*-ACP_X_ and acetyl-*S*-ACP_X_ in the presence or absence of KS_0_. (b) EICs of malonyl-PPant and sorbyl-PPant ejected from malonyl-*S*-ACP_2_ and sorbyl-*S*-ACP_2_, respectively, in the presence or absence of module X. (c) EICs of the 11+ charge states for acetyl-*S*-ACP_1_ and *holo*-ACP_1_ in the presence or absence of either *t*AT or *t*AT-TEII.

To assay the overall activity of modules X, 1 and 2, a bimodular protein (KS_1_-AT_1_-DH_1_-KR_1_-ACP_1_-KS_2_-DH_2_-ER_2_-KR_2_) lacking ACP_2_ was expressed and purified; separately, stand-alone *holo*-ACP_2_ was also expressed and purified (**Figure S3**). The two proteins were assayed via a phosphopantetheine (PPant) ejection assay^13^ in the presence of MBP-module X, truncated *t*AT (i.e., lacking its TE domain) and appropriate substrates. In this and all subsequent assays utilizing malonyl-CoA, this labile substrate was generated *in situ* by adding malonic acid, CoASH, ATP and the *Streptomyces coelicolor* malonyl-CoA synthetase MatB^14^ to the reaction mixture. Instead of detecting the anticipated hex-4-enoyl-PPant species, sorbyl-PPant was the major observed product (Figure 2b), implying that the enoylreductase (ER) domain of module 2 is inactive (designated ER_0_ from here onwards). Together, these results confirm our hypothesis that module Xmodule 1-module 2 comprise the first three modules for initiation of NOCAP biosynthesis.

We hypothesized that the TE domain of the *t*AT-TE protein hydrolyzes acyl-ACPs under conditions of “stalled” polyketide biosynthesis^15^. This suggestion was consistent with our earlier observation that use of the truncated *t*AT in place of the full-length *t*AT-TE resulted in a two-to-ten-fold decrease in product formation^5^. To test this hypothesis, we incubated Sfp-derived acetyl-*S*-ACP_1_ – a stalled acyl-ACP surrogate – with either *t*AT-TE or *t*AT. LC-MS analysis uncovered that acetyl-*S*-ACP1 was hydrolyzed to *holo*-ACP_1_ in the presence of *t*AT-TE but not *t*AT (Figures 2c, **S4**). Analogous radiolabeling experiments further verified the above findings (**Figure S5**). Together, these results provide strong evidence that this TE is a member of the “TEII” sub-family of thioesterases (designated TEII hereafter) that acts as a proofreading enzyme by hydrolyzing unproductive intermediates.

Buoyed by the reconstitution of modules X, 1 and 2, we endeavored to reconstitute *in vitro* the complete NOCAP synthase. To overcome its exceptionally large size (the synthase’s homodimeric mass approaches 3 MDa), NOCAP_PKS1 was expressed and purified as three standalone proteins: modules 1 and 2 as one bimodular protein, module 3 as a unimodular protein, and module 4 along with the KS domain of module 5 (module 4-KS_5_) as the third protein. Separately, NOCAP_PKS2 was dissociated into two proteins: the DH-ACP-KR tridomain of module 5 fused to the complete module 6 (DH_5_-ACP_5_-KR_5_-module 6), and a bimodular protein comprised of modules 7 and 8 along with the terminal TR domain (modules 7-8-TR) (Figures 3a, **S1**). To facilitate intermodular chain translocation between separated modules, each protein was fused to complementary N-terminal and/or C-terminal docking domains from DEBS that have previously been shown to facilitate non-covalent interactions between successive modules on a PKS assembly line^16,17^. These five NOCAP synthase-derived proteins were mixed with MBP-module X, *t*AT-TEII, malonyl-CoA, NADPH and *S*-adenosyl methionine. To confirm that products originated from the assembly-line PKS, [2-^13^C]-, [1,3-^13^C_2_]- or [^13^C_3_]- malonyl-CoA was used in place of malonyl-CoA in parallel reactions. By high-resolution MS, we identified a polyketide **1** with a molecular formula of C_23_H_26_O_4_ (observed [M-H]^−^ *m*/*z* 365.1762, theoretical [M-H]^−^ *m*/*z* 365.1753, 2.5 ppm) (Figure 3c).

**Figure 3.**
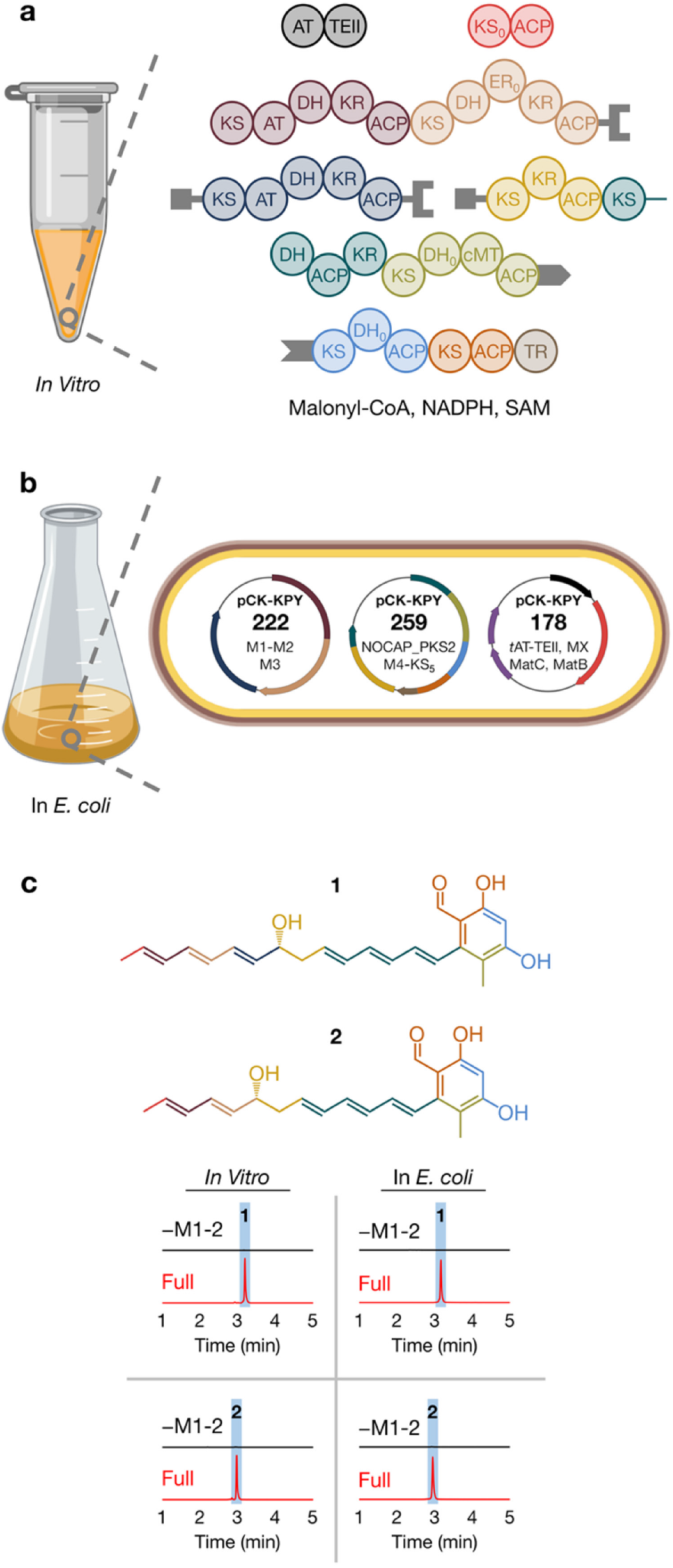
Reconstitution of the NOCAP synthase (a) *in vitro* and (b) in *E. coli*. (c) EICs of **1** and **2** from either *in vitro* reactions or *E. coli* pellet extracts. As a negative control, modules 1 and 2 (as one bimodular protein) were omitted.

The observation of +11, +11 and +22 mass shifts for **1** in mixtures containing [2-^13^C], [1,3-^13^C_2_] and [^13^C_3_] malonyl-CoA, respectively, indicated that **1** traversed the entire polyketide synthase and underwent three rounds of chain elongation, ketoreduction and dehydration at module 5 (Figures 1, **S6-7**). Because biosynthesis of **1** requires the entire assembly-line PKS, we propose that **1** is the aglycone product of the NOCAP synthase. A closely related polyketide **2** was detected that had presumably undergone one fewer round of chain elongation, ketoreduction and dehydration than **1** (molecular formula C_21_H_24_O_4_, observed [M-H]^−^ *m*/*z* 339.1604, theoretical [M-H]^−^ *m*/*z* 339.1596, 2.4 ppm) (**Figures S8-10**). Its MS/MS fragmentation pattern matched well with that of **1**, leading us to hypothesize that an upstream module was “skipped” during the biosynthesis of **2**. Two more minor polyketides **3** and **4** were identified with MS/MS fragmentation patterns noticeably different than **1** and **2** (**Figures S11-16**). **3** and **4** are most likely premature polyketides with a pyrone moiety that originated from spontaneous release after module 7 extension and C-1–C-5 oxygen lactonization.

For definitive structural analysis, we sought to produce the NOCAP synthase products by using *E. coli* as a heterologous host for scalable polyketide biosynthesis^6^. Informed by the *in vitro* reconstitution experiments summarized above, we engineered three plasmids with compatible antibiotic resistance markers and origins of replication that collectively encode the pathway. Plasmid pCKKPY_222_ encoded modules 1-2 as one bimodular protein and module 3. Plasmid pCK-KPY_259_ encoded module 4-KS_5_ and the intact NOCAP_PKS2, and pCK-KPY178 encoded the *t*AT-TEII and MBP-module X proteins (Figures 3b, **S17**). To enhance the malonyl-CoA pool in *E. coli*, pCK-KPY178 also encodes MatB and the *Rhizobium leguminosarum* malonate carrier protein MatC^18,19^. Gratifyingly, *E. coli* BAP1[pCK-KPY222/pCK-KPY259/pCK-KPY178] produced **1** and **2** (Figures 3c, **S18-23**). *E. coli*-derived **1** and **2** had the same MS/MS fragments as **1** and **2** produced *in vitro*. Derivatization of **1** and **2** with Girard’s reagent T^20^ confirmed the presence of an aldehyde (**Figures S24-27**). **3** and **4** were much lower in abundance from extracts of this strain, suggesting that these metabolites do not arise under physiological conditions, but are only synthesized under conditions with excess substrates. Because of their scarcity in *E. coli*, **3** and **4** were not further characterized.

We isolated **1** and **2** as faint yellow solids from 4 L of *E. coli* BAP1[pCK-KPY222/pCK-KPY259/pCK-KPY178] using lipid extraction with methyl *tert*-butyl ether/methanol^21^, C_18_ solid-phase extraction and UV-absorbance-guided semi-preparative HPLC (**Figures S28-30**). A number of 1D- and 2D-NMR experiments (^1^H, COSY, TOCSY, HSQC, HMBC, NOESY and ROESY) allowed us to fully elucidate their chemical structures (Figures 4, S**31-62**; **Table S2**). For **1**, COSY, TOCSY and HMBC experiments established carbon-carbon connectivity from C-1 (195.1 ppm) to C-22 (18.3 ppm) as well as the phenolic moiety resulting from C-2–C-7 aldol condensation. These spectra also revealed a pair of conjugated trienes, one synthesized by modules 1-3 and the other by module 5. The aldehyde substituent at C-1 shows that **1** and **2** were released from the assembly line by the terminal TR domain. We observed the expected hydroxyl substituents at C-3 (163.6 ppm, module 8), C-5 (160.8 ppm, module 7) and C-15 (71.9 ppm, module 4), a singlet methyl substituent at C-6 (114.7 ppm, module 6) and a terminal doublet methyl (C-22 for 1, C-20 for 2, module X). ROESY analysis of **1** and NOESY analysis of **2** verified that all of their double bonds have *trans* stereoconfigurations as predicted by bioinformatic analysis^22^. To determine the absolute configuration of the stereocenter set by the KR domain of module 4, we converted the C-15 hydroxyl substituent of **2** to a Mosher ester^23^. Mosher ester analysis with COSY confirmed that the absolute configuration at C-15 is *R*, also as predicted by bioinformatic analysis^22^ (**Figures S63-66**). Unlike **1**, compound **2** featured a conjugated diene – not triene – in its “tail”. We therefore hypothesized that a combination of the dissociated-by-design nature of module 3 and the broad substrate tolerance of the KS domain of module 4^5^ permitted facile chain translocation of the growing polyketide chain from module 2 to module 4. Indeed, *E. coli* that does not express module 3 only produced **2**, substantiating our hypothesis that the biosynthesis of **2** involves bypassing module 3 (**Figure S67**). Collectively, these spectroscopic efforts validated **1** as the aglycone product of the NOCAP synthase.

**Figure 4.**
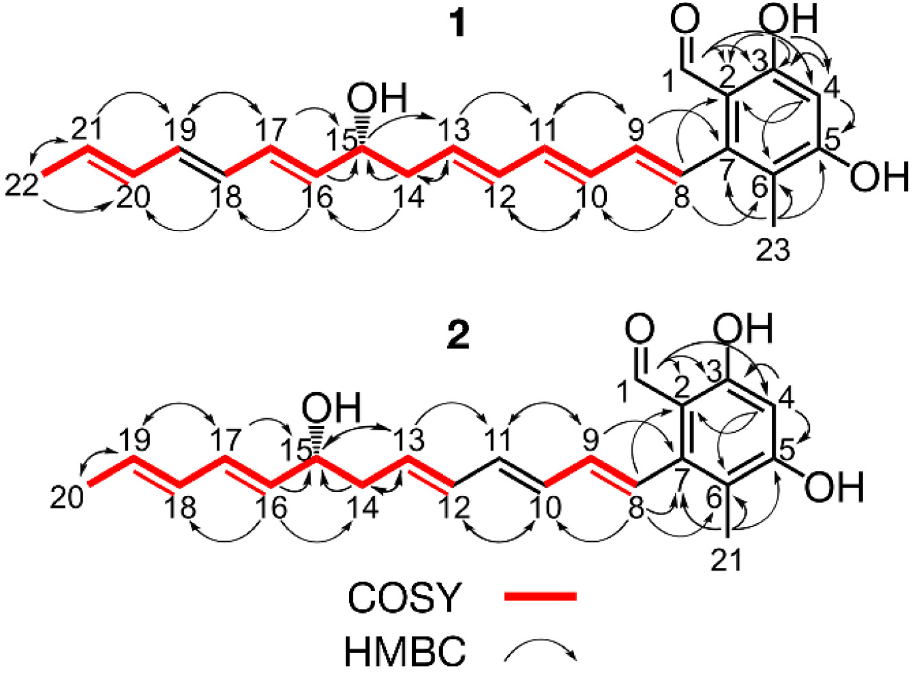
Structures of **1** and **2** assembled from 2D NMR data.

This report represents two milestones. First, we describe for the first time the full reconstitution (both *in vitro* as well as in *E. coli*) of an assembly-line PKS that is predominantly comprised of *trans*-AT modules. *trans*-AT PKSs represent over 23% of all sequenced assembly-line PKSs according to a recent survey^24^ and display remarkable architectural diversity; however, the understanding of *trans*-AT PKSs has significantly lagged that of *cis*-AT PKSs^2^. Based on this report, the NOCAP synthase is a strong contender as a model *trans*-AT PKS in the same vein as DEBS has been for *cis*-AT PKSs. Second, this work concludes the first example of natural product discovery by reconstituting orphan assembly-line PKSs *in vitro* or in *E. coli*. In principle, the methodology described here could be applied to other orphan polyketide natural products, especially those synthesized in low abundance or from unculturable organisms^25^. The discovery and structure elucidation of **1** will also allow us to turn our attention to the tailoring enzymes clustered with the NOCAP synthase genes and the ultimate characterization of the biological role of its fully decorated natural product. Such efforts are compellingly motivated by the statistically significant but nonetheless correlative occurrence of this PKS only in strains associated with clinical cases of nocardiosis.

## AUTHOR INFORMATION

### Author Contributions

^⊥^C.W.L. and S.R.L. contributed equally.

### Notes

The authors declare no competing financial interests.

## ACKNOWLEDGMENT

We thank past and present members of the C.K. laboratory for fruitful discussions and suggestions. We also thank Theresa McLaughlin (Stanford University) and Jeffrey G. Pelton (University of California, Berkeley) for technical assistance, and Prof. Joseph D. Puglisi for access to his 500 MHz NMR spectrometer. This work was supported by National Institutes of Health (NIH) Grant R01 GM087934 (to C.K.) and NIH Grant F32 GM123637 (to K.P.Y.). This work utilized the Stanford Cancer Institute Proteomics/Mass Spectrometry Shared Resource, which is supported by NIH Grant P30 CA124435, and the 900 MHz NMR spectrometer (funded by NIH Grant P41 GM068933) at the Central California 900 MHz NMR Facility.

