## Supporting Information for "Complete Reconstitution and Deorphanization of the 3 MDa NOCAP (NOCardiosis-Associated Polyketide) Synthase"

Figure S28. Isolation of **1** and **2** from *E. coli* BAP1[pCK-KPY222/pCK-KPY259/pCK-KPY178] ... 41
Figure S29. TIC, Mass Spectrum, Total UV Chromatogram and UV Spectrum of Purified **1** ... 42
Figure S30. TIC, Mass Spectrum, Total UV Chromatogram and UV Spectrum of Purified **2** .. 43

### Materials and Methods

##### Plasmids

Plasmids: *In Vitro* Reconstitution of the NOCAP Synthase.

| Plasmid Name | Vector Backbone | Protein(s) Over-Expressed by T7 Promoter |
|---|---|---|
| pCK-KPY028 | pET-28 (KanR) | MBP-*Na* $KS_X$-Strep-tag II |
| pCK-KPY054 | pET-21 (CbR) | *Na* $ACP_X$-6×His |
| pCK-KPY058 | pET-21 (CbR) | *Np* $KS_1$-$AT_1$-$DH_1$-$KR_1$-$ACP_1$-$KS_2$-$DH_2$-$ER_{02}$-$KR_2$-6×His |
| pCK-KPY059 | pET-28 (KanR) | MBP-*Na* Module X-Twin-Strep-tag |
| pCK-KPY099 | pET-28 (KanR) | DEBS(5)-*Na* Module 4-$KS_5$-Strep-tag II* |
| pCK-KPY102 | pCOLADuet-1 (KanR) | *Np* Modules 1-2-DEBS(4)-Strep-tag II* |
| pCK-KPY130 | pET-21 (CbR) | DEBS(5)-*Np* Module 3-DEBS(4)-Strep-tag II |
| pCK-KPY137 | pET-21 (CbR) | *Np* *t*AT-TEII-Strep-tag II |
| pCK-KPY142 | pET-21 (CbR) | DEBS(3)-*Na* Modules 7-8-TR-Strep-tag II |
| pCK-KPY144 | pCDFDuet-1 (StrR) | *Np* $DH_5$-$ACP_5$-$KR_5$-Module 6-DEBS(2)-Strep-tag II |
| pJK32[1] | pET-21 (CbR) | *Np* $ACP_1$-6×His* |
| pJK50[1] | pET-21 (CbR) | *Np* *t*AT-6×His |
| pJK72[1] | pET-21 (CbR) | *Np* $ACP_2$-6×His* |

Key: *Na*, *N. araoensis*; *Np*, *N. pneumoniae*; DEBS(2), DEBS Module 2 Linker; DEBS(3), DEBS Module 3 Linker; DEBS(4), DEBS Module 4 Linker; DEBS(5), DEBS Module 5 Linker; *, Codon Optimized for *E. coli*.

The plasmid encoding *S. coelicolor* MatB (pET-28a-$His_6$-MatB.SCo[2]) was a gift from Prof. Michelle Chang (University of California, Berkeley). The plasmid encoding *B. subtilis* Sfp (pSfp[3]) was a gift from Prof. Kira Weissman (University of Lorraine).

Plasmids: Heterologous Expression of the NOCAP Synthase in *E. coli*.

| **Plasmid Name** | **Vector Backbone** | **Protein(s) Over-Expressed By T7 Promoter** |
|---|---|---|
| pCK-KPY102 | pCOLADuet-1 (KanR) | *Np* Modules 1-2-DEBS(4)-Strep-tag II* |
| pCK-KPY178 | pCDFDuet-1 (StrR) | 1: 6×His-*Np* *t*AT-TEII, 2: MBP-*Na* Module X<br>3: *S. coelicolor* MatB, 4: *R. leguminosarum* MatC* |
| pCK-KPY222 | pCOLADuet-1 (KanR) | 1: *Np* Modules 1-2-DEBS(4)*<br>2: DEBS(5)-*Na* Module 3-DEBS(2)* |
| pCK-KPY259 | pETDuet-1 (CbR) | 1: DEBS(3)-*Na* Module 4-$KS_5$*<br>2: *Na* $DH_5$-$ACP_5$-$KR_5$-Module 6-7-8-TR* |
| pCK-KPY292 | pCOLADuet-1 (KanR) | DEBS(5)-*Na* Module 3-DEBS(2)* |

Key: *Na*, *N. araoensis*; *Np*, *N. pneumoniae*; DEBS(2), DEBS Module 2 Linker; DEBS(3), DEBS Module 3 Linker; DEBS(4), DEBS Module 4 Linker; DEBS(5), DEBS Module 5 Linker; *, Codon Optimized for *E. coli*.

Genes were amplified from *Nocardia (N. araoensis* or *N. pneumoniae*) genomic DNA (Leibniz Institute, DSMZ-German Collection of Microorganisms and Cell Cultures GmbH) or DNA codon optimized for *E. coli* (Gen9, GENEWIZ, Integrated DNA Technologies and Twist Bioscience) using Q5 High-Fidelity DNA Polymerase (New England Biolabs) and cloned into pET and Duet vectors (Novagen) using restriction enzyme-based methods (New England Biolabs), Gibson assembly[4] or commercial assembly master mixes (NEBuilder HiFi DNA Assembly Master Mix, New England Biolabs and In-Fusion HD Cloning Plus, Takara Bio). Plasmids were cloned and maintained in *E. coli* DH5α, Stellar (Takara Bio) or *E. coli* TOP10 (Thermo Fisher Scientific).

**Protein Expression and Purification**

*LB Broth*: 25.0 g/L LB Broth, Miller granulated powder (Thermo Fisher Scientific) (supplemented with 100 mg/L carbenicillin disodium (Gold Biotechnology), 50 mg/L kanamycin monosulfate (Gold Biotechnology) or 50 mg/L streptomycin sulfate (Gold Biotechnology))

*Ni-NTA Lysis Solution*: 50 mM sodium phosphate (Thermo Fisher Scientific), 500 mM sodium chloride (Thermo Fisher Scientific), 10% (v/v) glycerol (Thermo Fisher Scientific), 10 mM imidazole (MilliporeSigma), pH 7.4, 1-2.5 mg/mL chicken egg white lysozyme (Alfa Aesar)

*Ni-NTA Wash Solution*: 50 mM sodium phosphate (Thermo Fisher Scientific), 300 mM sodium chloride (Thermo Fisher Scientific), 10% (v/v) glycerol (Thermo Fisher Scientific), 25 mM imidazole (MilliporeSigma), pH 7.4

*Ni-NTA Elution Solution*: 50 mM sodium phosphate (Thermo Fisher Scientific), 20 mM sodium chloride (Thermo Fisher Scientific), 10% (v/v) glycerol (Thermo Fisher Scientific), 500 mM imidazole (MilliporeSigma), pH 7.4

*Strep-Tactin Lysis Solution*: 50 mM sodium phosphate (Thermo Fisher Scientific), 500 mM sodium chloride (Thermo Fisher Scientific), 10% (v/v) glycerol (Thermo Fisher Scientific), pH 7.8, 1-2.5 mg/mL chicken egg white lysozyme (Alfa Aesar), 4 mg/L chicken egg white avidin (MilliporeSigma)

*Strep-Tactin Wash Solution*: 50 mM sodium phosphate (Thermo Fisher Scientific), 500 mM sodium chloride (Thermo Fisher Scientific), 10% (v/v) glycerol (Thermo Fisher Scientific), pH 7.8

*Strep-Tactin Elution Solution*: 50 mM sodium phosphate (Thermo Fisher Scientific), 500 mM sodium chloride (Thermo Fisher Scientific), 10% (v/v) glycerol (Thermo Fisher Scientific), pH 7.8, 5 mM d-desthiobiotin (MilliporeSigma)

*FPLC A*: 50 mM sodium phosphate (Thermo Fisher Scientific), 10% (v/v) glycerol (Thermo Fisher Scientific), pH 7.4

*FPLC B*: 50 mM sodium phosphate (Thermo Fisher Scientific), 1 M sodium chloride (Thermo Fisher Scientific), 10% (v/v) glycerol (Thermo Fisher Scientific), pH 7.4

A single colony of *E. coli* BL21(DE3) (for *apo*-protein) or *E. coli* BAP1 (for *holo*-protein[5]) housing the appropriate plasmid was used to inoculate an overnight seed culture of LB Broth (40 mL) and grown at 30ºC. The overnight seed culture was pelleted by centrifugation at 4000 × *g* for 10 min at room temperature, re-suspended in fresh LB Broth (40 mL), diluted into 8 × 1 L of LB Broth in 2.5 L Tunair shake flasks (IBI Scientific) and agitated at 37ºC until the $OD_{600}$ reached 0.2-0.3. At this point, the incubator temperature was set to 16°C. After the addition of 0.1 mL/L culture of 1 M isopropyl β-D-1-thiogalactopyranoside (Gold Biotechnology) at an $OD_{600}$ of 0.4-0.6, the cultures were agitated at 16ºC for an additional 16 h. Cells were harvested by centrifugation at 5000 × *g* for 15 minutes at 4ºC, frozen in liquid nitrogen and stored at -80ºC.

Thawed cells were re-suspended either in Ni-NTA Lysis Solution or Strep-Tactin Lysis Solution, incubated at 4ºC for 1 h and sonicated. Lysates were clarified twice by centrifugation at 25,000 × *g* for 1 h at 4ºC and incubated overnight while rotating end-over-end with either 2 mL/L culture HisPur Ni-NTA resin (Thermo Fisher Scientific) or 0.5 mL/L culture Strep-Tactin Sepharose resin (IBA Lifesciences). The protein-bound resin was washed with either Ni-NTA Wash Solution or Strep-Tactin Wash Solution. Protein was eluted off the resin with either Ni-NTA Elution Solution or Strep-Tactin Elution Solution and further purified with a gradient elution method (FPLC A to FPLC B over 20 column volumes at 4 mL/min) on an ÄKTA pure chromatography system (GE Healthcare Life Sciences) equipped with a HiTrap Q HP anion exchange chromatography column (5 mL). Fractions identified by SDS-PAGE to contain proteins of interest were concentrated using Amicon Ultra Centrifugal Filters (MilliporeSigma), frozen in liquid nitrogen and stored at -80ºC.

For the purification of MBP-module X, a Pierce Protease Inhibitor Tablet (Thermo Fisher Scientific) was added for every 50 mL of Strep-Tactin Lysis Solution.

Approximate protein yields were 9 mg/L for MBP-module X (CK-KPY059), 4 mg/L for module 1-2-DEBS module 4 Linker (CK-KPY102), 1 mg/L for DEBS module 5 linker-module 3-DEBS module 4 linker (CK-KPY130), 5 mg/L for DEBS module 5 linker-module 4-$KS_5$ (CK-KPY099), 3 mg/L for $DH_5$-$ACP_5$-$KR_5$-module 6-DEBS module 2 linker (CK-KPY144), 5 mg/L for DEBS module 3 linker-module 7-8-TR (CK-KPY142), and 2 mg/L for *t*AT-TEII (CK-KPY137).

***In Vitro* Reconstitution of Modules X, 1-2 and *t*AT-TEII**

*Partial Reconstitution Solution*: 100 mM HEPES (Thermo Fisher Scientific), pH 7.5, 10 mM adenosine 5′-triphosphate (MilliporeSigma), 5 mM NADPH (MilliporeSigma), 10 mM magnesium chloride (Thermo Fisher Scientific), 10 mM tris(2-carboxyethyl)phosphine (Thermo Fisher Scientific)

*HPLC A*: 99.9% (v/v) water, 0.1% (v/v) formic acid (Thermo Fisher Scientific)

*HPLC B*: 99.9% (v/v) acetonitrile, 0.1% (v/v) formic acid (Thermo Fisher Scientific)

Purified proteins (final concentration of 2-5 μM) were mixed into the Partial Reconstitution Solution. For ACPs, final concentrations of 50-100 μM were used.

For *in vitro* characterization of module X, Sfp-catalyzed conversions of *apo*-$ACP_X$ to malonyl-*S*-$ACP_X$ were initiated by the addition of malonyl-CoA (final concentration of 1 mM) (MilliporeSigma); $KS_X$ was added 30 min later and incubated at room temperature for an additional 10 min. Reactions were desalted using Zeba Spin Desalting Columns, 7K MWCO, 0.5 mL (Thermo Fisher Scientific), diluted with water and analyzed by LC-MS (Waters SQ Detector 2 LC-MS system).

For *in vitro* characterization of modules X and 1-2, reactions were initiated by the addition of malonic acid (final concentration of 5 mM) (MilliporeSigma) and coenzyme A (final concentration of 2.5 mM) (MilliporeSigma) and incubated at room temperature for 30 min. In this and all assays utilizing both malonic acid and coenzyme A, the *Streptomyces coelicolor* malonyl-CoA synthetase MatB[6] generated malonyl-CoA *in situ*. Reactions were desalted using Zeba Spin Desalting Columns, 7K MWCO, 0.5 mL (Thermo Fisher Scientific), diluted with water and analyzed by LC-MS (Waters SQ Detector 2 LC-MS system).

For *in vitro* characterization of *t*AT-TEII, Sfp-catalyzed conversions of *apo*-$ACP_1$ to acetyl-*S*-$ACP_1$ were initiated by the addition of acetyl-CoA (final concentration of 1 mM) (MilliporeSigma); *t*AT or *t*AT-TEII was added 30 min later and incubated at room temperature for an additional 10 min (up to 60 min for radiolabeling experiments). Reactions were desalted using Zeba Spin Desalting Columns, 7K MWCO, 0.5 mL (Thermo Fisher Scientific), diluted with and analyzed by LC-MS (Waters SQ Detector 2 LC-MS system). In analogous radiolabeling experiments, [1-$^{14}$C] acetyl-CoA was used in place of acetyl-CoA. Reactions were separated by SDS-PAGE; the gels were washed with water, stained with SimplyBlue SafeStain (Thermo Fisher Scientific) and dried *in vacuo*. The radiolabeling of ACP1 was recorded lane-by-lane with a $^{14}C$ linear analyzer detector (Raytest RITA Star).

For LC-MS analysis, proteins were separated with a gradient elution method (98/2 HPLC A/HPLC B to 5/95 HPLC A/HPLC B over 4 min at 0.3 mL/min) on a Waters SQ Detector 2 LC-MS system equipped with an Agilent ZORBAX StableBond 300 C8 column (3.5 μm, 2.1 x 50 mm) and operating on positive ion mode.

***In Vitro* Reconstitution of the NOCAP Synthase**

*Full Reconstitution Solution*: 100 mM HEPES (Thermo Fisher Scientific), pH 7.5, 10 mM adenosine 5′-triphosphate (MilliporeSigma), 2.5 mM coenzyme A (MilliporeSigma), 5 mM NADPH (MilliporeSigma), 2.5 mM *S*-adenosyl-L-methionine (MilliporeSigma), 10 mM magnesium chloride (Thermo Fisher Scientific), 10 mM tris(2-carboxyethyl)phosphine (Thermo Fisher Scientific)

*HPLC A*: 99.9% (v/v) water, 0.1% (v/v) formic acid (Thermo Fisher Scientific)

*HPLC B*: 99.9% (v/v) acetonitrile, 0.1% (v/v) formic acid (Thermo Fisher Scientific)

Purified proteins (final concentration of 2-5 μM) were mixed into the Full Reconstitution Solution. Reactions were initiated by the addition of malonic acid (final concentration of 5 mM) (MilliporeSigma) and incubated overnight at room temperature. To precipitate the proteins, 0.1 mL of methanol (Thermo Fisher Scientific) was added to 0.1 mL of reaction, vortexed and centrifuged at 15,000 × *g* for 10 min at room temperature. The supernatants were collected, dried *in vacuo*, re-dissolved in methanol and analyzed by LC-MS (Waters SQ Detector 2 LC-MS system and/or Agilent 6545 Q-TOF LC-MS system). Using sodium phosphate in place of HEPES did not improve yields of **1**, **2**, **3** or **4**. Samples were protected from light at all times.

For LC-MS analysis, compounds were separated with a gradient elution method (95/5 HPLC A/HPLC B to 5/95 HPLC A/HPLC B over 4 min at 0.6 mL/min) on an Agilent Infinity 1290 II HPLC/6545 Q-TOF MS system equipped with an Agilent ZORBAX RRHD Extend-C18 column (1.8 μm, 2.1 x 50 mm) and operating on negative ion mode. MS/MS analysis used fragmentation energies of 10, 20 or 40 V; supplementary figures feature spectra with fragmentation energies of 20 V. Alternatively, compounds were separated with a gradient elution method (98/2 HPLC A/HPLC B to 5/95 HPLC A/HPLC B over 4 min at 0.3 mL/min) on a Waters SQ Detector 2 LC-MS system equipped with an Agilent InfinityLab Poroshell 120 SB-C18 column (2.7 µm, 2.1 x 50 mm) and operating on both positive and negative ion mode.

**Biosynthesis of 1 and 2**

*LB Broth*: 25.0 g/L LB Broth, Miller granulated powder (Thermo Fisher Scientific) (supplemented with 100 mg/L carbenicillin disodium (Gold Biotechnology), 50 mg/L kanamycin monosulfate (Gold Biotechnology) and 50 mg/L streptomycin sulfate (Gold Biotechnology))

*Terrific Broth*: 47.6 g/L Terrific Broth (modified) powder (MilliporeSigma), 10.1 g/L glycerol (MilliporeSigma) (supplemented with 100 mg/L carbenicillin disodium (Gold Biotechnology), 50 mg/L kanamycin monosulfate (Gold Biotechnology) and 50 mg/L streptomycin sulfate (Gold Biotechnology))

A single colony of *E. coli* BAP1[pCK-KPY222/pCK-KPY259/pCK-KPY178] was used to inoculate an overnight seed culture of LB Broth (40 mL) grown at 30ºC. The overnight seed culture was pelleted by centrifugation at 4000 × *g* for 10 min at room temperature, re-suspended in Terrific Broth (40 mL), diluted into 4 × 1 L of Terrific Broth in 2.5 L Tunair shake flasks (IBI Scientific) and agitated at 30ºC until the $OD_{600}$ reached 0.2. After the addition of 10 mL/L culture of 500 mM sodium malonate, pH 7.4 (MilliporeSigma), 1 mL/L culture of 50 mM calcium D-pantothenate (MilliporeSigma) and 0.1 mL/L culture of 1 M isopropyl β-D-1-thiogalactopyranoside (Gold Biotechnology), the cultures were agitated at 16ºC for an additional 72 h. Cells were harvested by centrifugation at 5000 × *g* for 15 min at 16ºC, frozen in liquid nitrogen and stored at -80ºC.

To confirm the biosynthesis of **1** and **2**, small samples of 250 mg of frozen cells were re-suspended in 0.3 mL n-hexane (Thermo Fisher Scientific) and 0.2 mL isopropanol (Thermo Fisher Scientific) and vortexed with glass beads for 30 min. After centrifugation at 15,000 × *g* for 10 min, the upper organic phases were collected, dried in *vacuo*, re-dissolved in methanol and analyzed by LC-MS (Waters SQ Detector 2 LC-MS system and/or Agilent 6545 Q-TOF LC-MS system). Samples were protected from light at all times.

To convert the aldehyde-containing **1** to the hydrazone-containing **5** and the aldehyde-containing **2** to the hydrazone-containing **6**, 0.1 mL extracts containing **1** and **2** were incubated for 1 hr at room temperature after the addition of 0.1 mL 2% (w/v) Girard's reagent T (MilliporeSigma) in methanol and 20 µL glacial acetic acid (Thermo Fisher Scientific). Samples were protected from light at all times.

As a negative control in which modules 1-2 is omitted, *E. coli* BAP1 was transformed with pCK-KPY259, pCK-KPY178 and pCK-KPY292 – in place of pCK-KPY222. As a negative control in which module 3 is omitted, *E. coli* BAP1 was transformed with pCK-KPY259, pCK-KPY178 and pCK-KPY102 – in place of pCK-KPY222.

**Isolation of 1 and 2**

*HPLC A*: 99.9% (v/v) water, 0.1% (v/v) formic acid (Thermo Fisher Scientific)

*HPLC B*: 99.9% (v/v) acetonitrile, 0.1% (v/v) formic acid (Thermo Fisher Scientific)

Guided by Matyash's lipid extraction method[7], thawed cells were re-suspended in 65 mL/L culture water, 150 mL/L culture methanol (Thermo Fisher Scientific) and 500 mL/L culture methyl *tert*-butyl ether (Thermo Fisher Scientific) and sonicated in a Branson Ultrasonics M1800 ultrasonic cleaning bath for 1 h. Phase separation was induced by the addition of 65 mL/L culture of brine. The upper organic phases were collected, pooled, dried *in vacuo* and re-dissolved in 20 mL 25% (v/v) methanol in HPLC B. This mixture was diluted to a final volume of 90 mL with HPLC A and equally loaded onto six pre-equilibrated C18 solid-phase extraction cartridges (3M Empore 7 mm/3 mL). The cartridges were each washed with 1 mL HPLC A, 1 mL 25/75 HPLC B/HPLC A, 1 mL 50/50 HPLC B/HPLC A, 1 mL 75/25 HPLC B/HPLC A and 1 mL HPLC B. Fractions identified by LC-MS (Waters SQ Detector 2 LC-MS system and/or Agilent 6545 Q-TOF LC-MS system) to contain **1** and **2** (50/50 HPLC B/HPLC A, 75/25 HPLC B/HPLC A and HPLC B) were pooled, dried *in vacuo*, re-dissolved in 35/65 HPLC B/HPLC A and filtered through a 0.45 μm PFTE membrane (VWR). This mixture was separated with a gradient elution method (35/65 HPLC B/HPLC A to 80/20 HPLC B/HPLC A over 1 h at 2 mL/min) on an Agilent 1260 Infinity LC system equipped with an Agilent Eclipse XDB-C8 column (5 μm, 250 mm x 9.4 mm). Fractions identified by LC-MS to contain **1** and **2** were dried *in vacuo* and re-dissolved in 0.3 mL chloroform-d (ACROS Organics). **1** eluted off this column at 67/33 HPLC B/HPLC A; **2** eluted off this column at 62/38 HPLC B/HPLC A. Typical yields were on the order of 1-10 mg/L culture.

According to Hoye's Mosher ester analysis protocol[8], the *R*-MTPA-**2** and *S*-MTPA-**2** esters were prepared with *S*-(+)-MTPA-Cl and *R*-(−)-MTPA-Cl (MilliporeSigma), respectively. Samples were protected from light at all times.

**Structural Elucidation of 1 and 2**

Samples for NMR were prepared in 0.3 mL chloroform-d (ACROS Organics) in matched symmetrical Shigemi tubes (5 mm).

NMR spectra of **1** were acquired on a 900 MHz Bruker AVANCE II spectrometer (Central California 900 MHz NMR Facility, California Institute for Quantitative Biosciences, University of California, Berkeley) with a 5 mm TCI (H{CN}, Z-gradient) cryoprobe, running TopSpin v3.2. Sample temperature was regulated at 25°C. Experiments acquired with **1** (see table below for experimental parameters) include $^1$H 1-D, COSY, HSQC, HMBC, TOCSY (60 ms mixing) and ROESY (200 ms mixing).

NMR spectra of **2**, *R*-MTPA-**2** and *S*-MTPA-**2** were acquired on a 500 MHz Bruker AVANCE spectrometer (Puglisi Laboratory, Department of Structural Biology, Stanford University School of Medicine) with a 5 mm TCI (H{CN}), Z-gradient) cryoprobe, running TopSpin v1.3. Sample temperature was regulated at 25°C. Experiments acquired with **2** (see table below for experimental parameters) include $^1$H 1-D, COSY, HSQC, HMBC, TOCSY (60 ms mixing) and NOESY (1 s mixing). Experiments acquired with *R*-MTPA-**2** and *S*-MTPA-**2** (see table below for experimental parameters) include $^1$H 1-D, COSY and HSQC.

NMR data were processed on the respective spectrometers and analyzed using SPARKY[9,10].

NMR Analysis of **1**: Experimental Parameters

| **1** | NS | TD | SW (ppm) | SFO1 (MHz) | O1p (ppm) | TD1 | SW1 (ppm) | SFO2 (MHz) | O2p (ppm) |
|---|---|---|---|---|---|---|---|---|---|
| $^1$H 1D | 128 | 131072 | 14.03 | 900.26 | 6.15 | | | | |
| COSY | 2 | 4096 | 14.03 | 900.26 | 6.15 | 1024 | 13.99 | 900.26 | 6.15 |
| TOCSY (60 ms mixing) | 4 | 4096 | 14.03 | 900.26 | 6.15 | 1024 | 13.99 | 900.26 | 6.15 |
| ROESY (200 ms mixing) | 16 | 4096 | 14.03 | 900.26 | 6.15 | 1024 | 13.99 | 900.26 | 6.15 |
| HSQC | 8 | 2048 | 16.02 | 900.26 | 6.15 | 256 | 164.82 | 226.39 | 79.99 |
| HSQC (aromatic region) | 8 | 2048 | 16.02 | 900.26 | 6.15 | 128 | 60.01 | 226.41 | 199.96 |
| HMBC | 32 | 4096 | 14.03 | 900.26 | 6.15 | 256 | 220.86 | 226.39 | 99.82 |

NMR Analysis of **2**: Experimental Parameters

| **2** | NS | TD | SW (ppm) | SFO1 (MHz) | O1p (ppm) | TD1 | SW1 (ppm) | SFO2 (MHz) | O2p (ppm) |
|---|---|---|---|---|---|---|---|---|---|
| $^1$H 1D | 128 | 65536 | 14.00 | 500.25 | 4.50 | | | | |
| COSY | 4 | 1024 | 13.33 | 500.25 | 6.17 | 512 | 13.33 | 500.25 | 6.17 |
| TOCSY (60 ms mixing) | 8 | 2048 | 12.01 | 500.25 | 5.00 | 400 | 12.00 | 500.25 | 5.00 |
| NOESY (1 s mixing) | 16 | 2048 | 12.01 | 500.25 | 5.00 | 400 | 12.00 | 500.25 | 5.00 |
| HSQC | 16 | 1024 | 13.01 | 500.25 | 4.50 | 216 | 222.04 | 125.80 | 98.99 |
| HMBC | 128 | 2048 | 13.01 | 500.25 | 4.50 | 216 | 222.04 | 125.80 | 98.99 |

NMR Analysis of *R*-MTPA-**2** and *S*-MTPA-**2**: Experimental Parameters

| *R*-MTPA-**2** & *S*-MTPA-**2** | NS | TD | SW (ppm) | SFO1 (MHz) | O1p (ppm) | TD1 | SW1 (ppm) | SFO2 (MHz) | O2p (ppm) |
|---|---|---|---|---|---|---|---|---|---|
| $^1$H 1D | 256 | 65536 | 14.00 | 500.25 | 4.50 | | | | |
| COSY | 2 | 1024 | 13.33 | 500.25 | 6.17 | 336 | 13.33 | 500.25 | 6.17 |
| HSQC | 4 | 1024 | 13.01 | 500.25 | 4.50 | 256 | 222.04 | 125.80 | 98.99 |

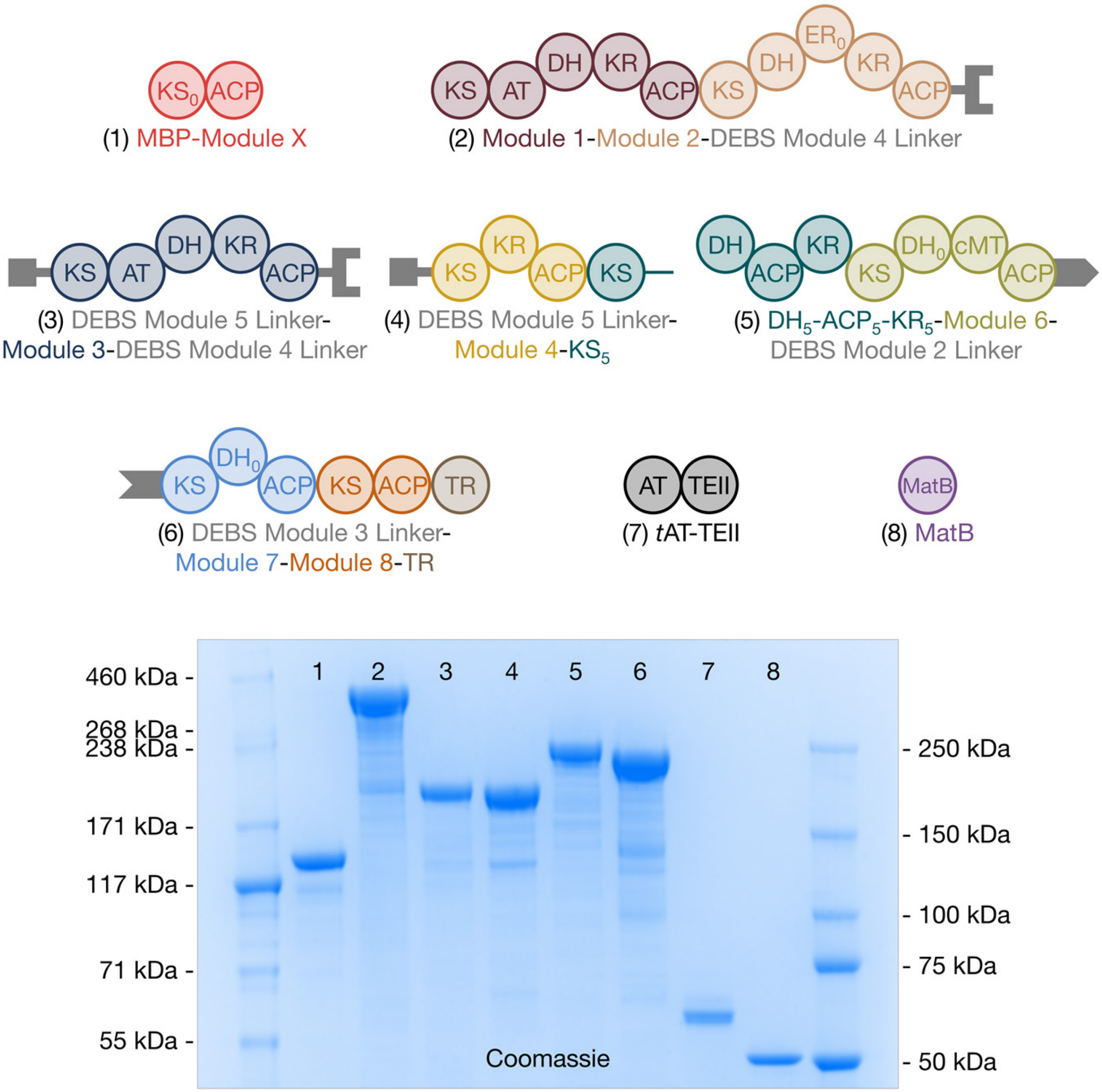

**Figure S1**. SDS-PAGE of purified proteins used for the *in vitro* reconstitution of the NOCAP synthase: **1**, MBP-Module X (CK-KPY059); **2**, Module 1-Module 2-DEBS Module 4 Linker (CK-KPY102); **3**, DEBS Module 5 Linker-Module 3-DEBS Module 4 Linker (CK-KPY130); **4**, DEBS Module 5 Linker-Module 4-$KS_5$ (CK-KPY099); **5**, $DH_5$-$ACP_5$-$KR_5$-Module 6-DEBS Module 2 Linker (CK-KPY144); **6**, DEBS Module 3 Linker-Module 7-Module 8-TR (CK-KPY142); **7**, *t*AT-TEII (CK-KPY137); and **8**, MatB (pET-28a-$His_6$-MatB.SCo).

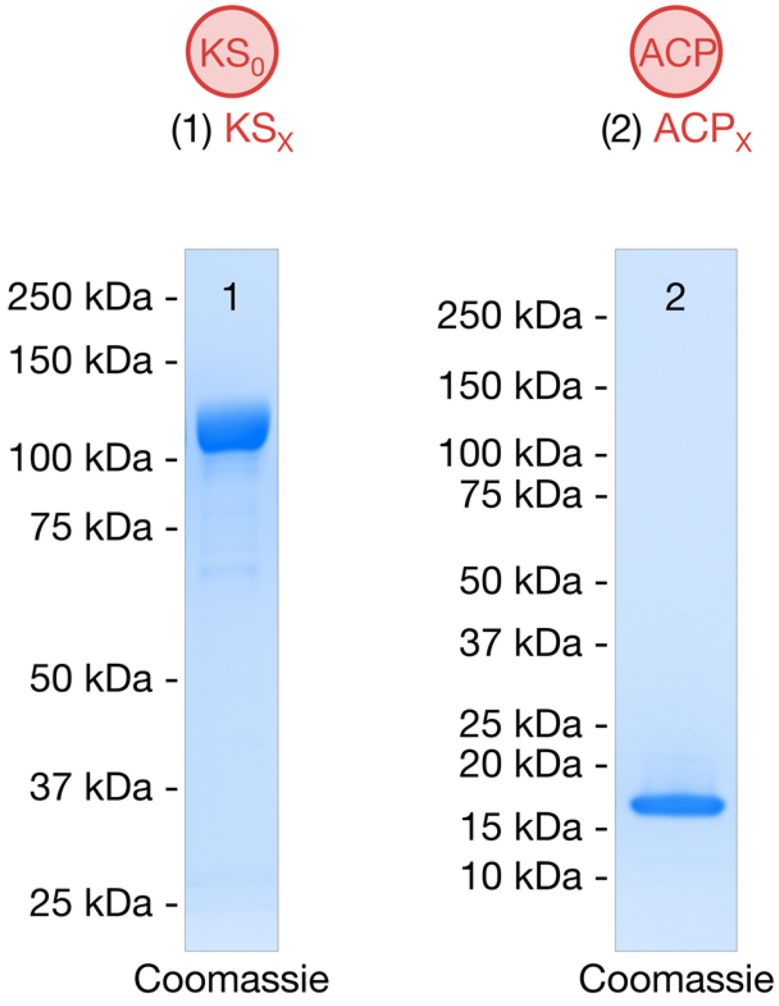

**Figure S2**. SDS-PAGE of purified **1**, MBP-KS$_X$ (CK-KPY028) and **2**, ACP$_X$ (CK-KPY054).

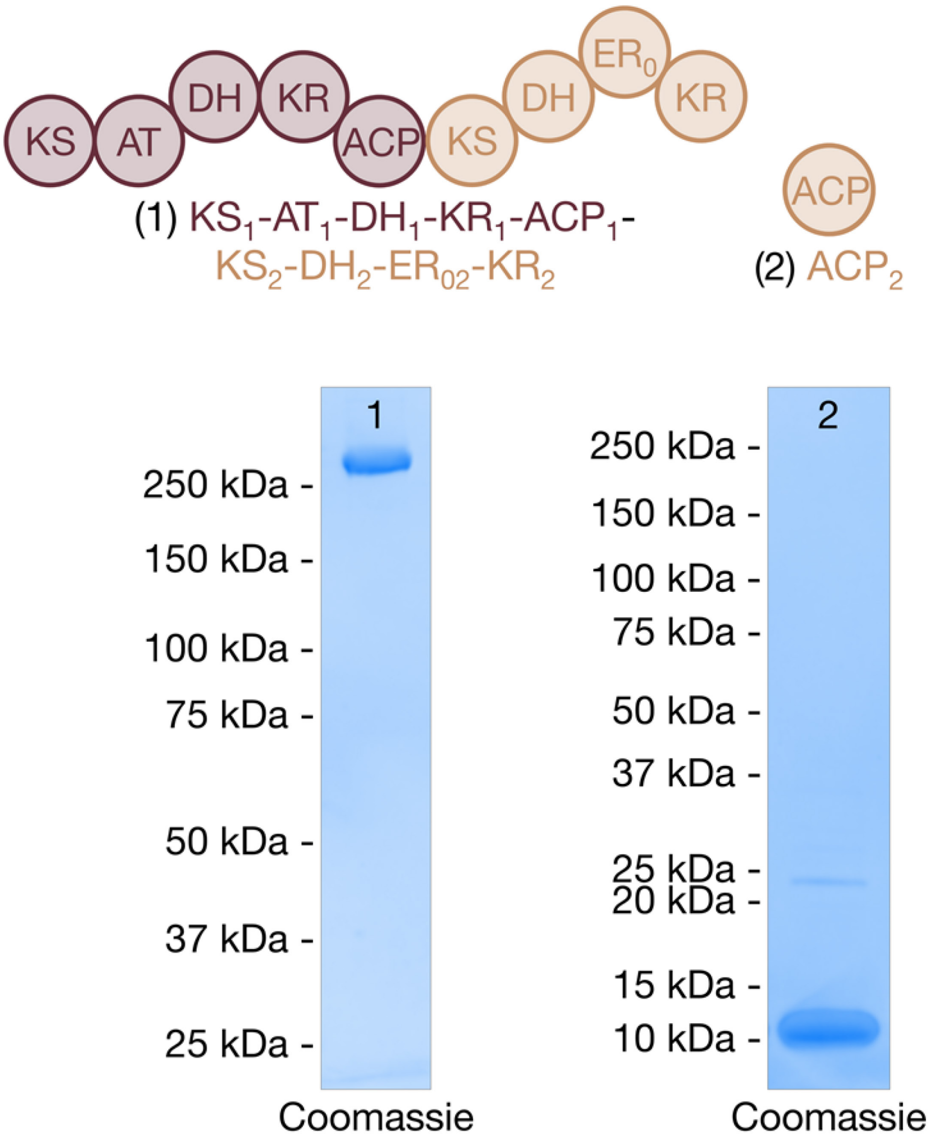

**Figure S3**. SDS-PAGE of purified **1**, $KS_1$-$AT_1$-$DH_1$-$KR_1$-$ACP_1$-$KS_2$-$DH_2$-$ER_{02}$-$KR_2$ (CK-KPY058) and **2**, $ACP_2$ (JK72).

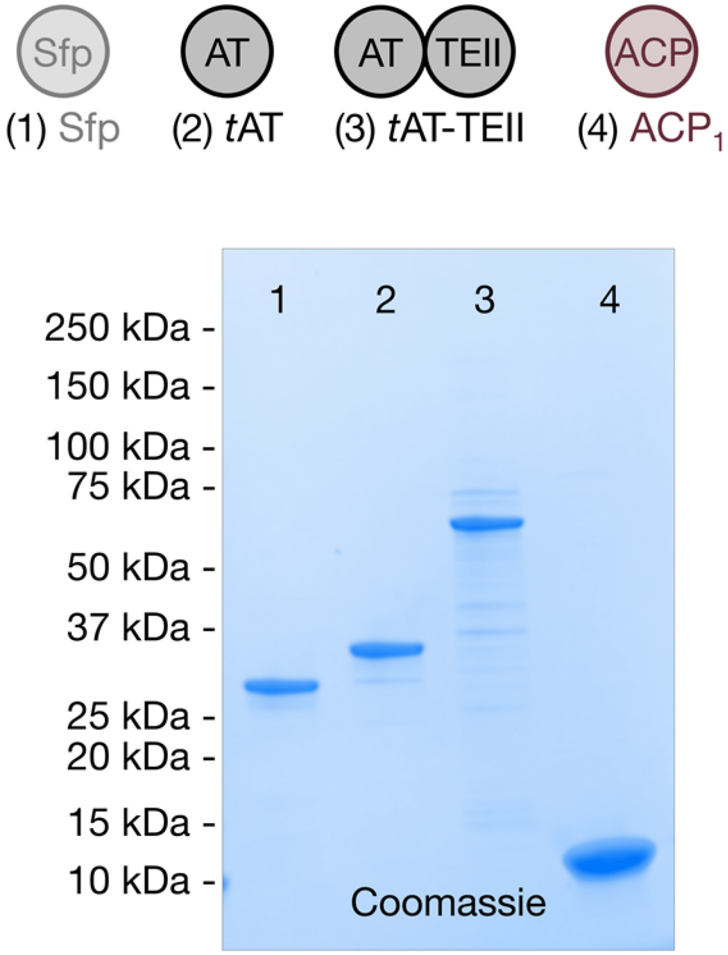

**Figure S4**. SDS-PAGE of purified **1**, Sfp; **2**, *t*AT (JK50); **3**, *t*AT-TEII (CK-KPY137); and **4**, $ACP_1$ (JK32).

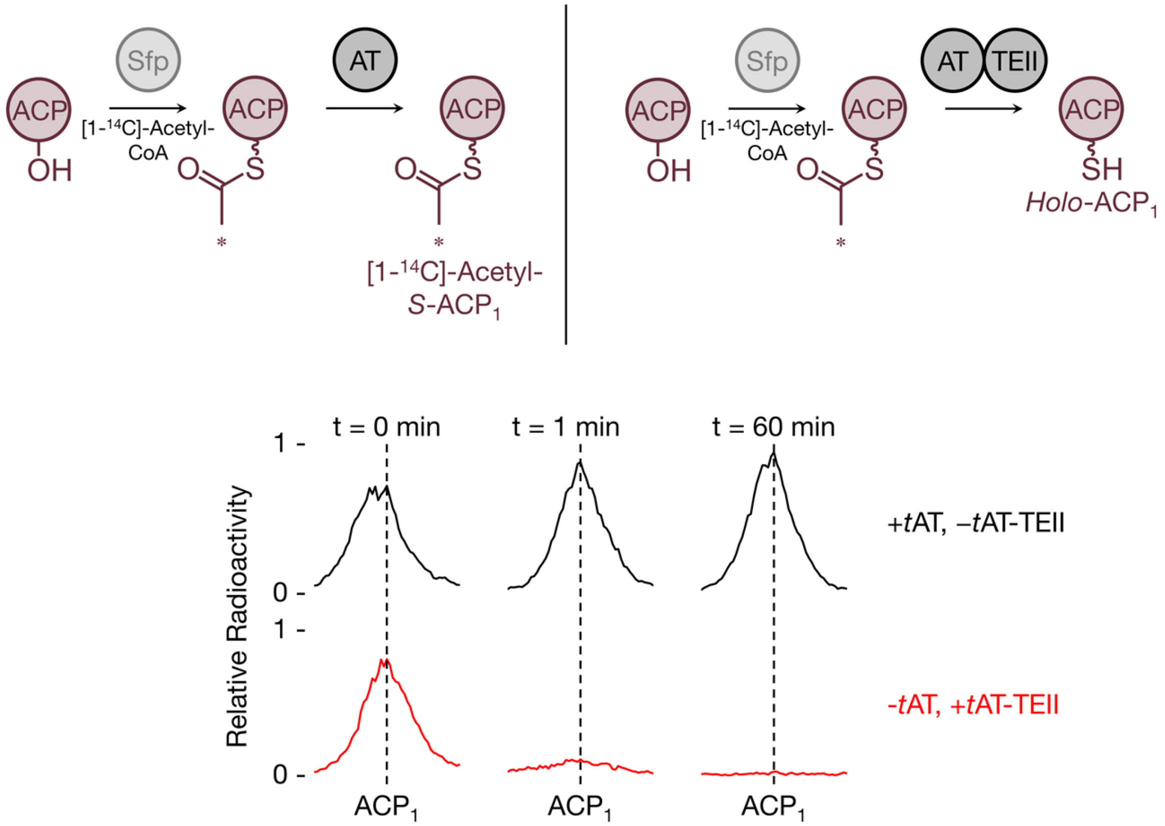

**Figure S5**. Sfp-derived [1-$^{14}$C]-acetyl-*S*-ACP$_1$ – a stalled acyl-ACP surrogate – was incubated with either *t*AT-TEII (bottom row of plots) or *t*AT (top row of plots) and subjected to SDS-PAGE analysis. Radioactivity was monitored with a $^{14}$C linear analyzer detector across ACP$_1$. Upon adding *t*AT-TEII, the radiolabeling of ACP$_1$ fell to less than 20% of its initial value in as little as 1 min. On the other hand, upon adding the truncated *t*AT, radiolabeling remained steady for at least 1 h.

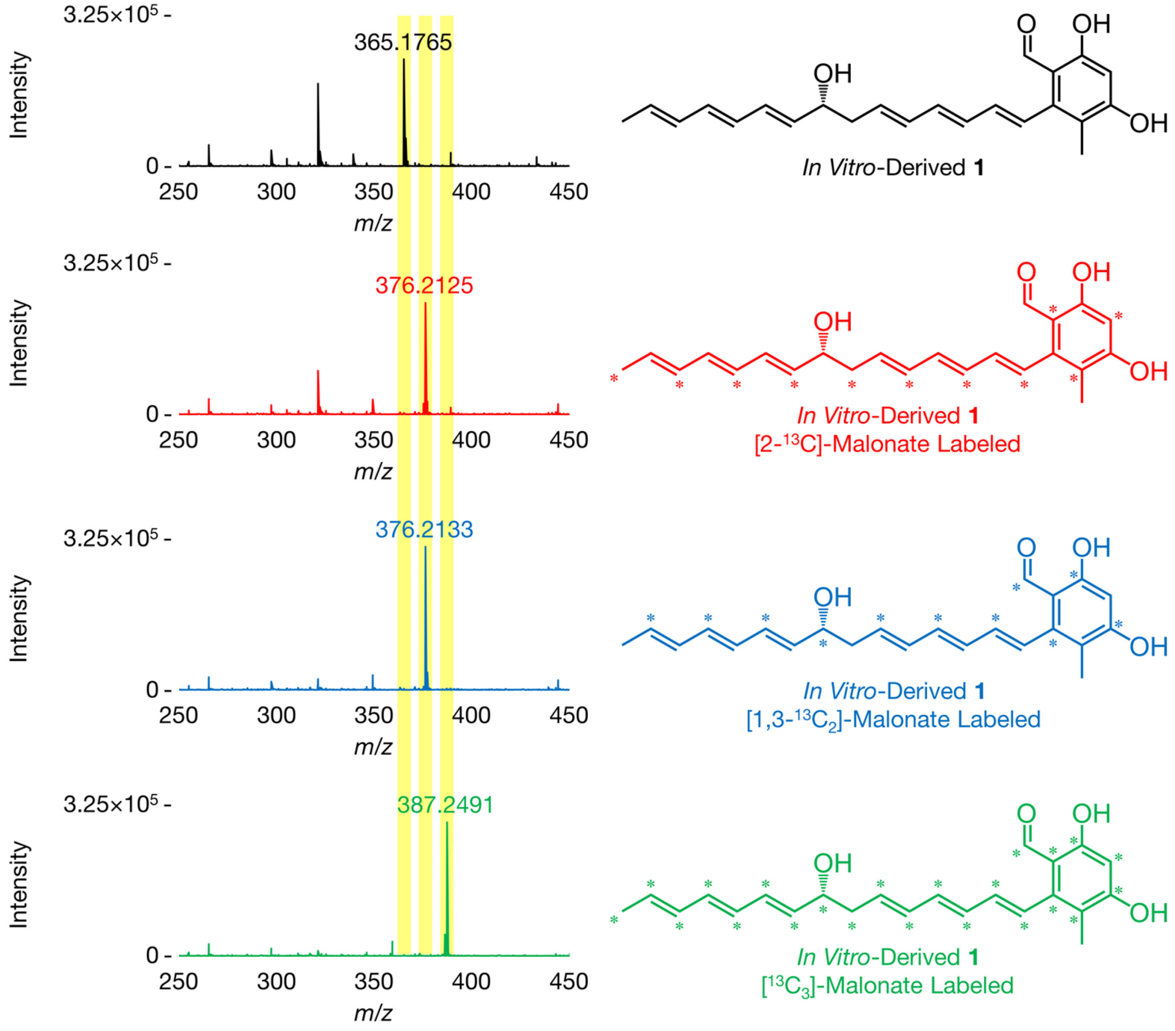

**Figure S6**. Mass spectra (ESI-) of *in vitro*-derived **1** (black); **1**, [2-$^{13}$C]-malonate labeled (red); **1**, [1,3-$^{13}C_2$]-malonate labeled (blue); and **1**, [$^{13}C_3$]-malonate labeled (green). **1** has a molecular formula of $C_{23}H_{26}O_4$ (observed [M-H]$^-$ *m/z* 365.1762, theoretical [M-H]$^-$ *m/z* 365.1753, 2.5 ppm). Spectra acquired on an Agilent 6545 Q-TOF LC-MS system and are representative of at least three independent experimental replicates.

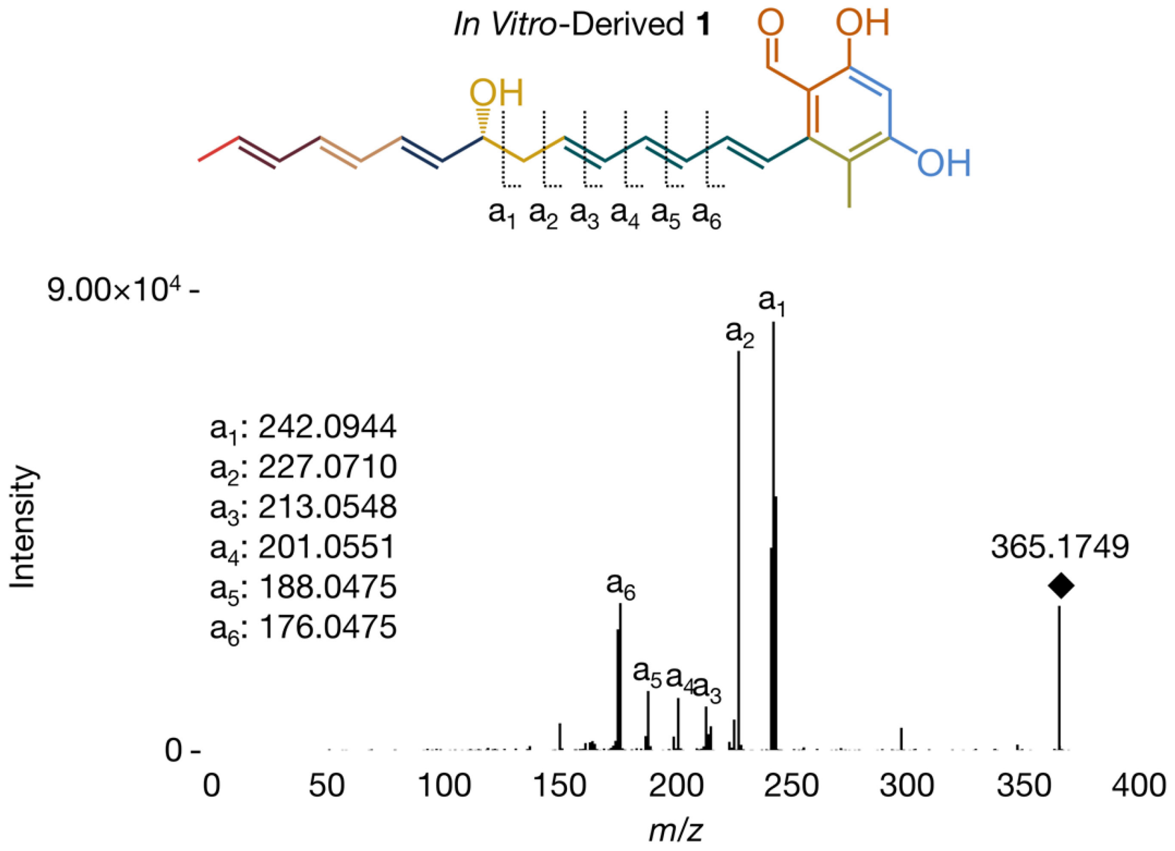

**Figure S7**. MS/MS fragmentation (ESI-) of *in vitro*-derived **1** yields six characteristic fragments. Spectrum acquired on an Agilent 6545 Q-TOF LC-MS system and is representative of at least three independent experimental replicates.

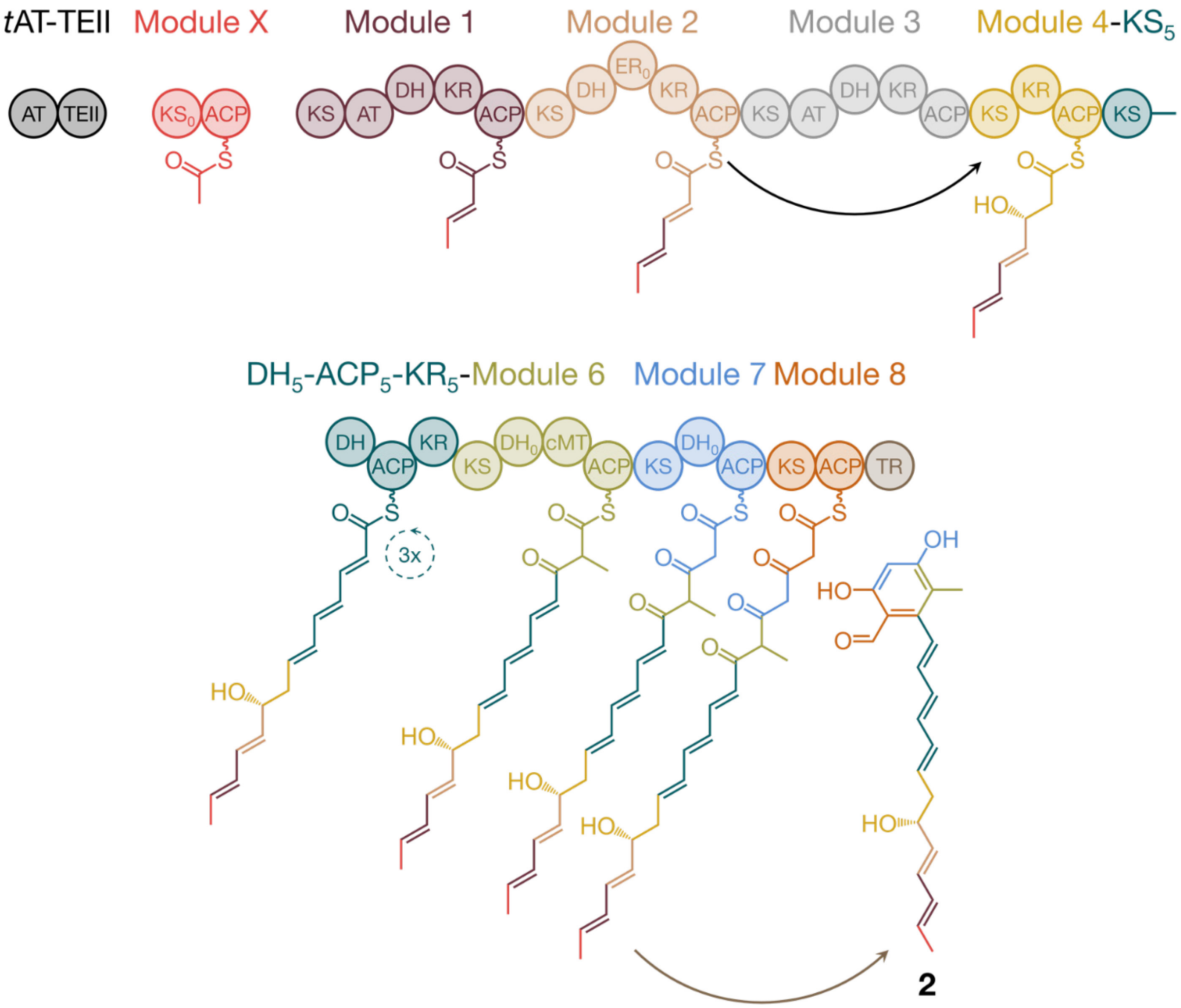

**Figure S8**. Biosynthesis of **2** by the NOCAP synthase.

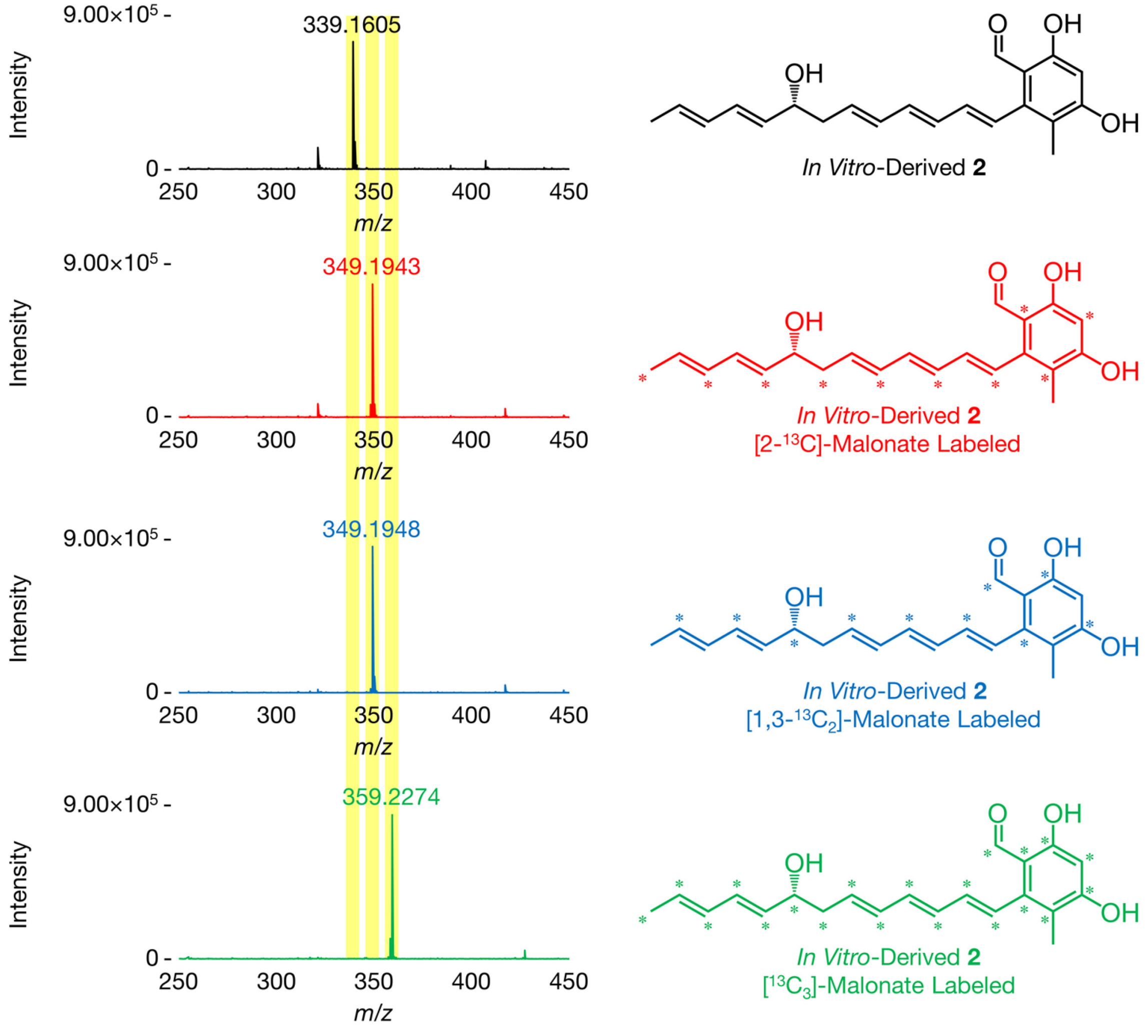

**Figure S9**. Mass spectra (ESI-) of *in vitro*-derived **2** (black); **2**, [2-$^{13}$C]-malonate labeled (red); **2**, [1,3-$^{13}C_2$]-malonate labeled (blue); and **2**, [$^{13}C_3$]-malonate labeled (green). **2** has a molecular formula of $C_{21}H_{24}O_4$ (observed [M-H]$^-$ *m/z* 339.1604, theoretical [M-H]$^-$ *m/z* 339.1596, 2.4 ppm). Spectra acquired on an Agilent 6545 Q-TOF LC-MS system and are representative of at least three independent experimental replicates.

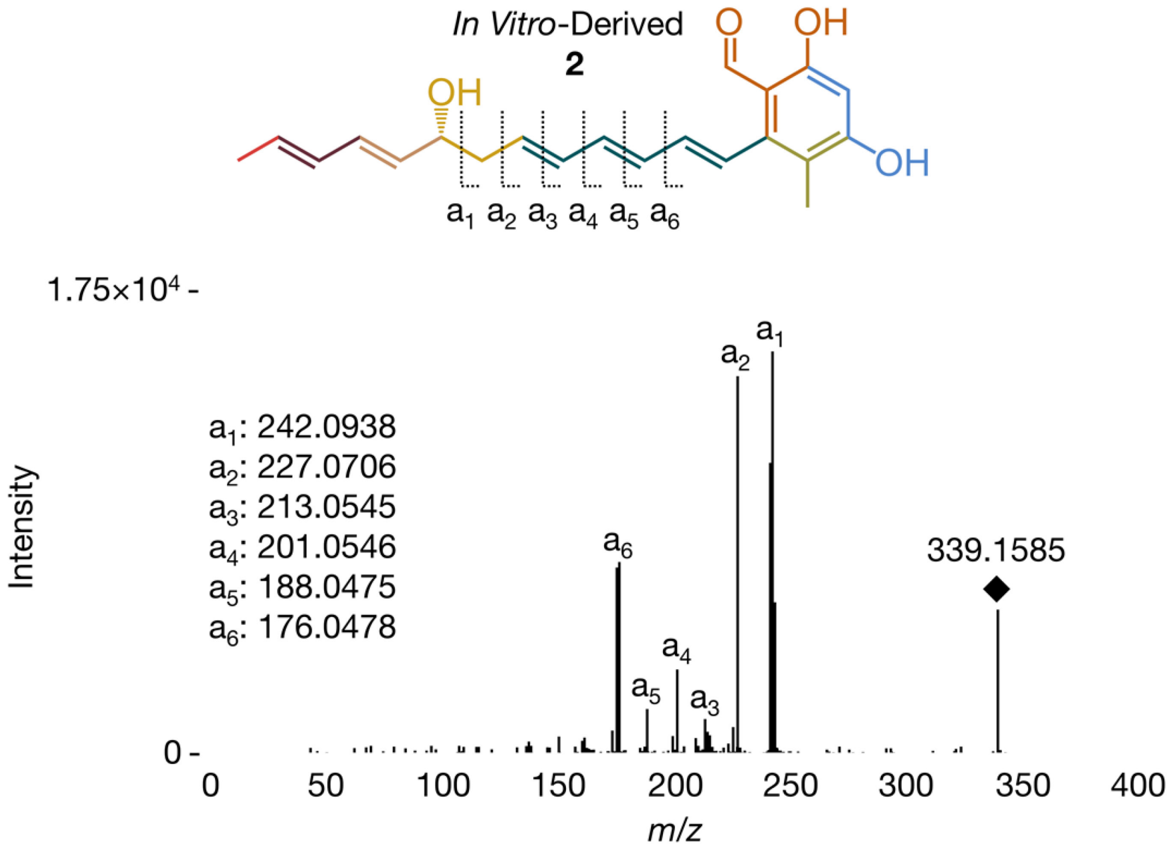

**Figure S10**. MS/MS fragmentation (ESI-) of *in vitro*-derived **2** yields six characteristic fragments. Spectrum acquired on an Agilent 6545 Q-TOF LC-MS system and is representative of at least three independent experimental replicates.

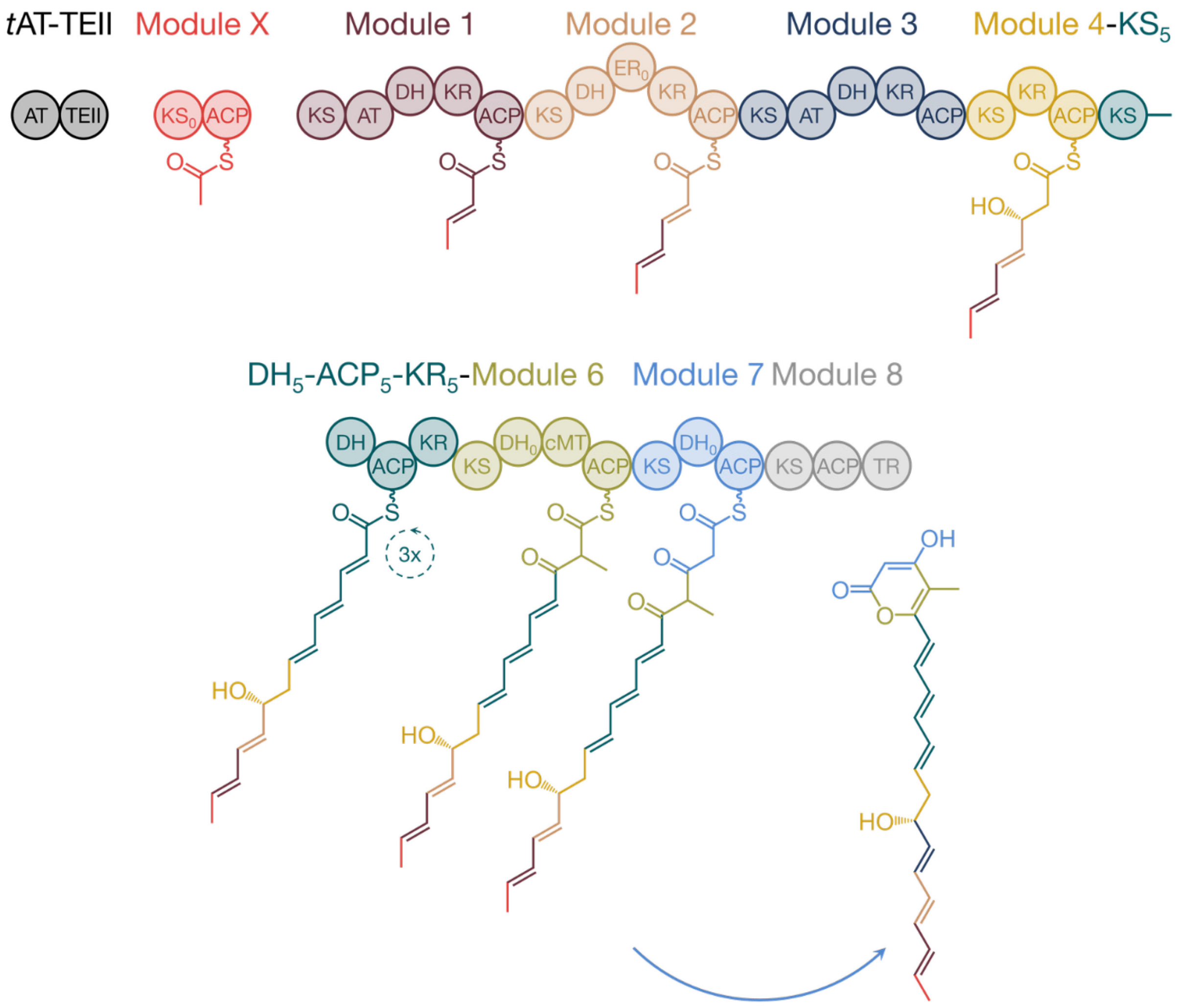

**3**

**Figure S11**. Biosynthesis of **3** by the NOCAP synthase.

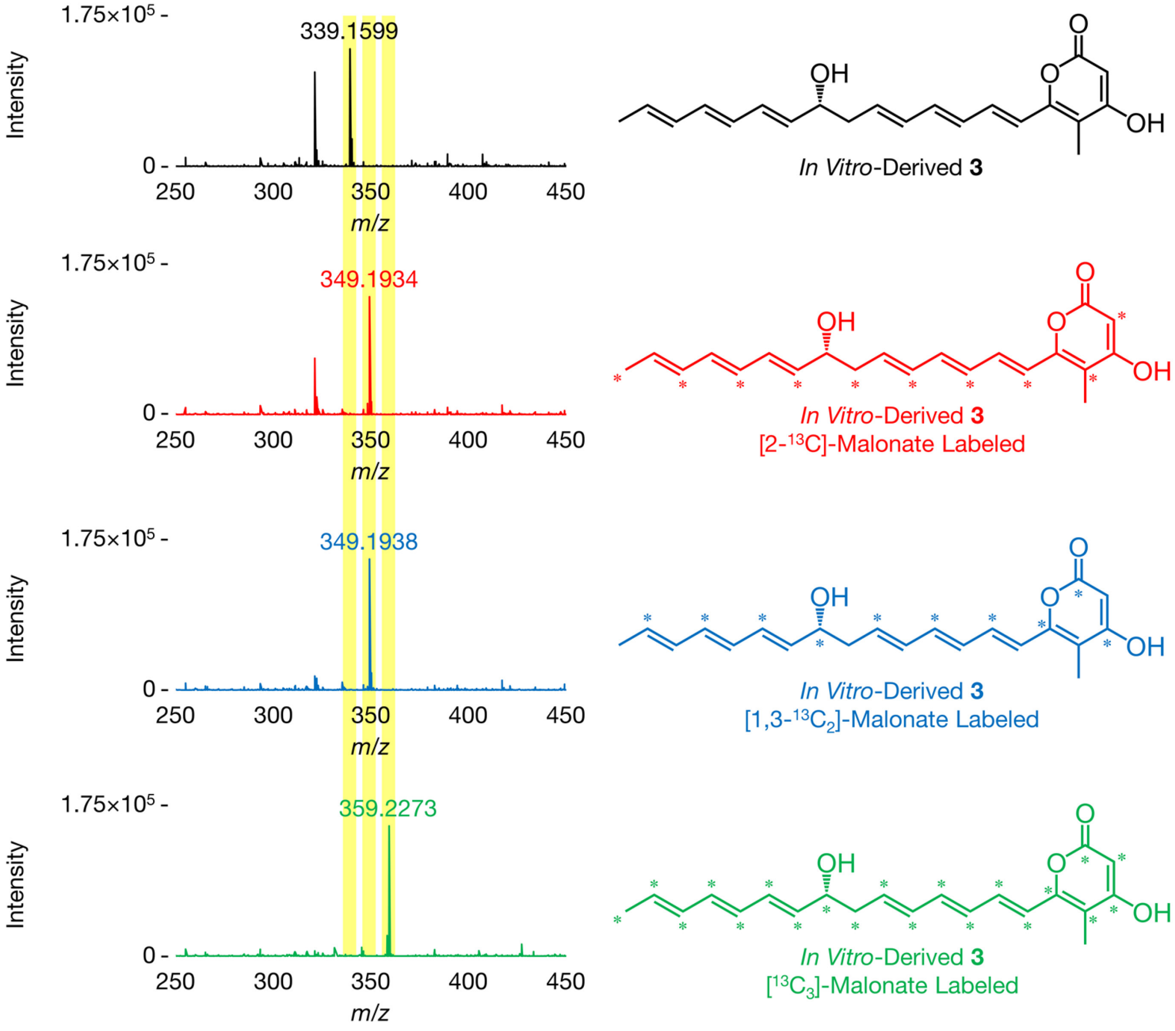

**Figure S12**. Mass spectra (ESI-) of *in vitro*-derived **3** (black); **3**, [2-$^{13}$C]-malonate labeled (red); **3**, [1,3-$^{13}C_2$]-malonate labeled (blue); and **3**, [$^{13}C_3$]-malonate labeled (green). **3** has a molecular formula of $C_{21}H_{24}O_4$ (observed $[M\text{-}H]^-$ *m/z* 339.1602, theoretical $[M\text{-}H]^-$ *m/z* 339.1596, 1.8 ppm). Spectra acquired on an Agilent 6545 Q-TOF LC-MS system and are representative of at least three independent experimental replicates.

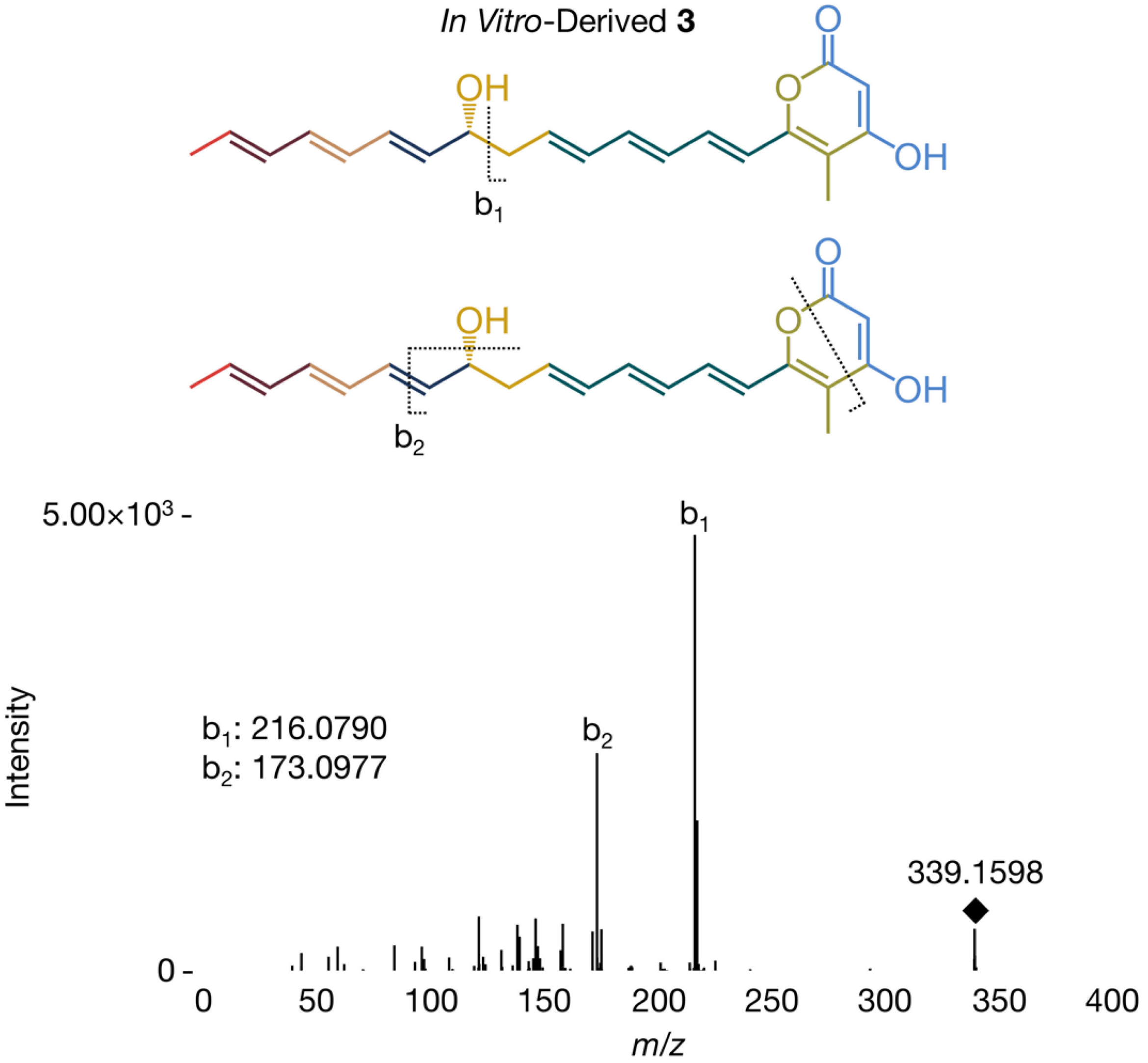

**Figure S13**. MS/MS fragmentation (ESI-) of *in vitro*-derived **3** yields two characteristic fragments. Spectrum acquired on an Agilent 6545 Q-TOF LC-MS system and is representative of at least three independent experimental replicates.

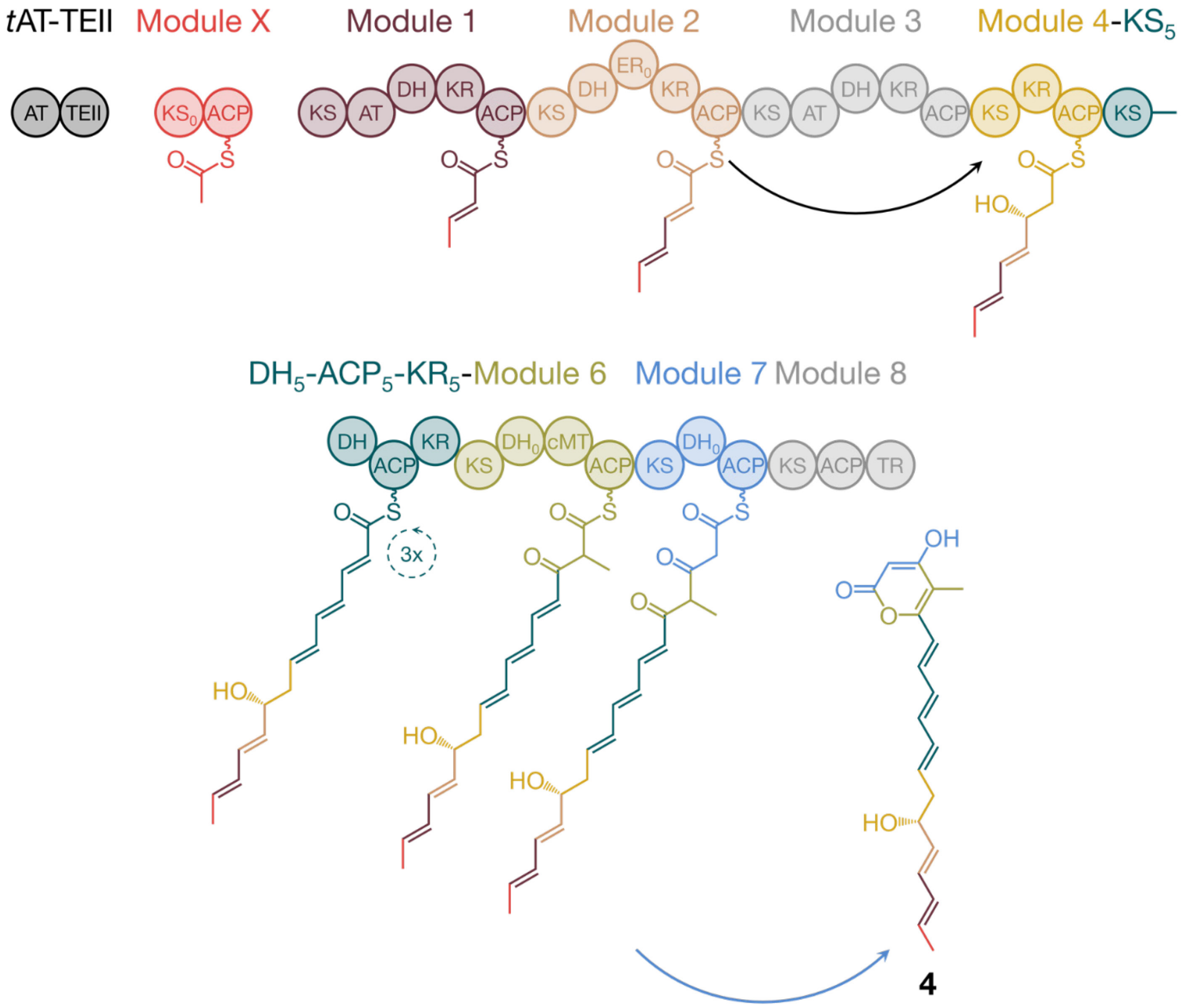

4

**Figure S14**. Biosynthesis of **4** by the NOCAP synthase.

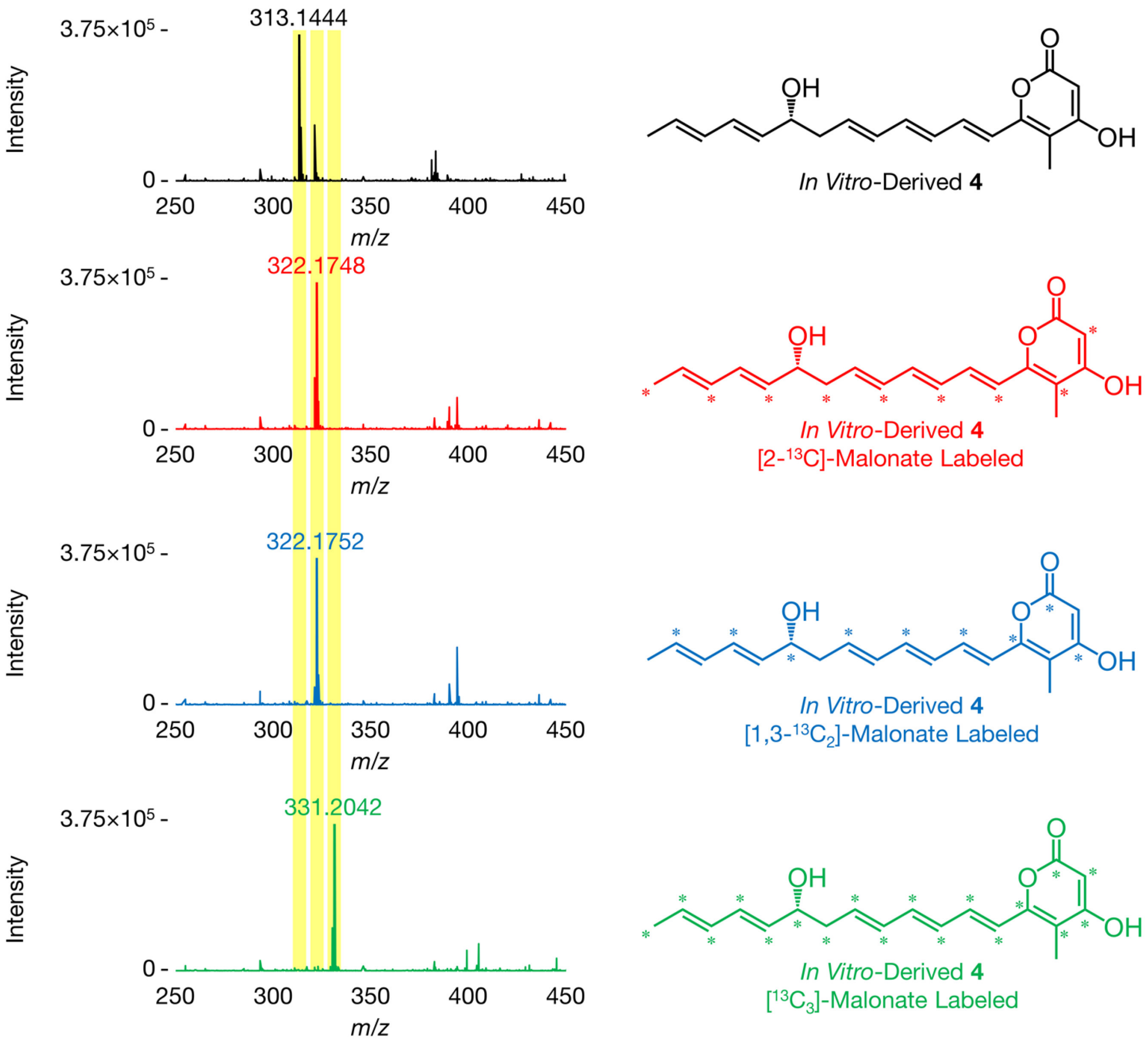

**Figure S15**. Mass spectra (ESI-) of *in vitro*-derived **4** (black); **4**, [2-$^{13}$C]-malonate labeled (red); **4**, [1,3-$^{13}C_2$]-malonate labeled (blue); and **4**, [$^{13}C_3$]-malonate labeled (green). **4** has a molecular formula of $C_{19}H_{22}O_4$ (observed [M-H]$^-$ *m/z* 313.1446, theoretical [M-H]$^-$ *m/z* 313.1440, 1.9 ppm). Spectra acquired on an Agilent 6545 Q-TOF LC-MS system and are representative of at least three independent experimental replicates.

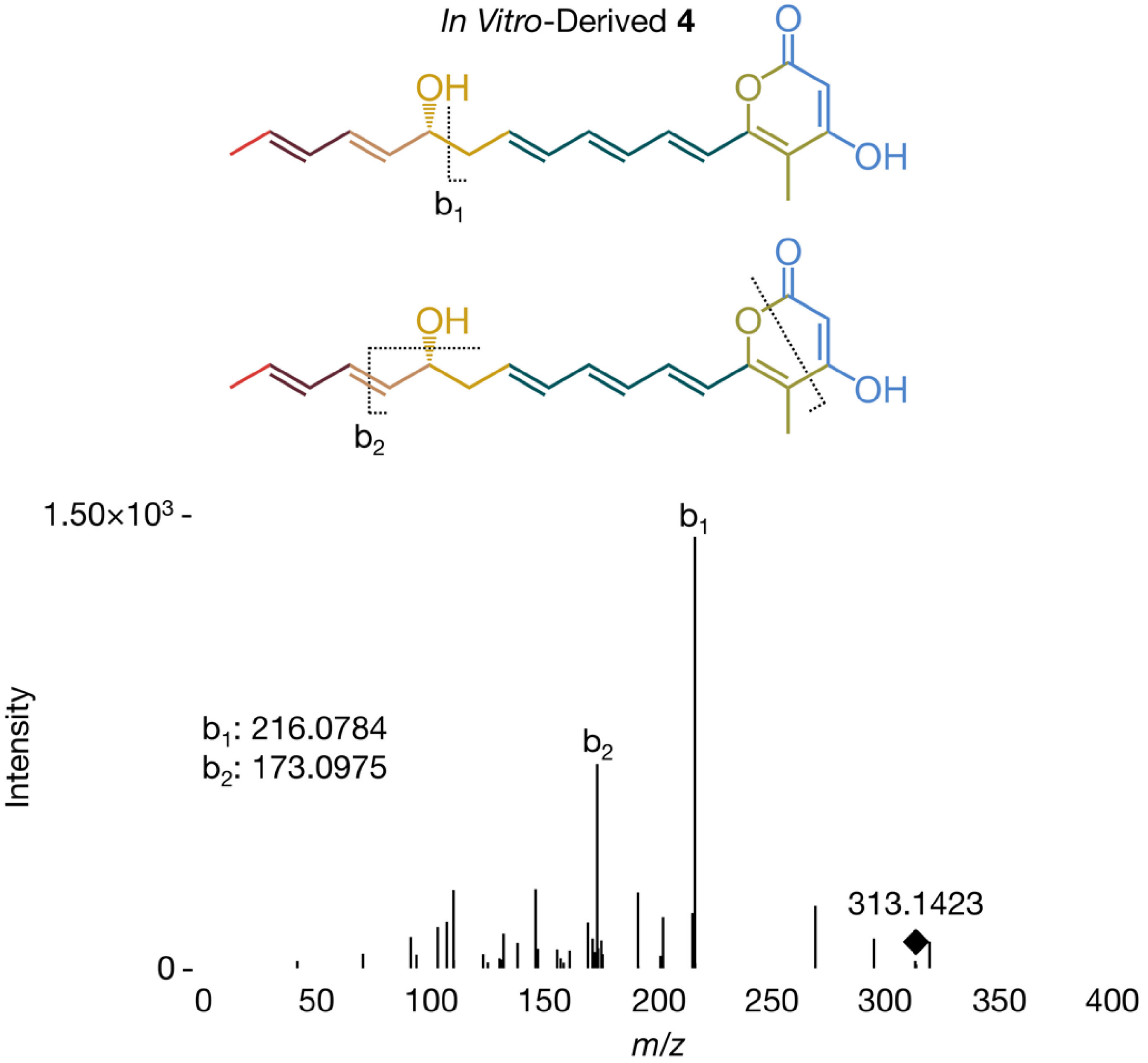

**Figure S16**. MS/MS fragmentation (ESI-) of *in vitro*-derived **4** yields two characteristic fragments. Spectrum acquired on an Agilent 6545 Q-TOF LC-MS system and is representative of at least three independent experimental replicates.

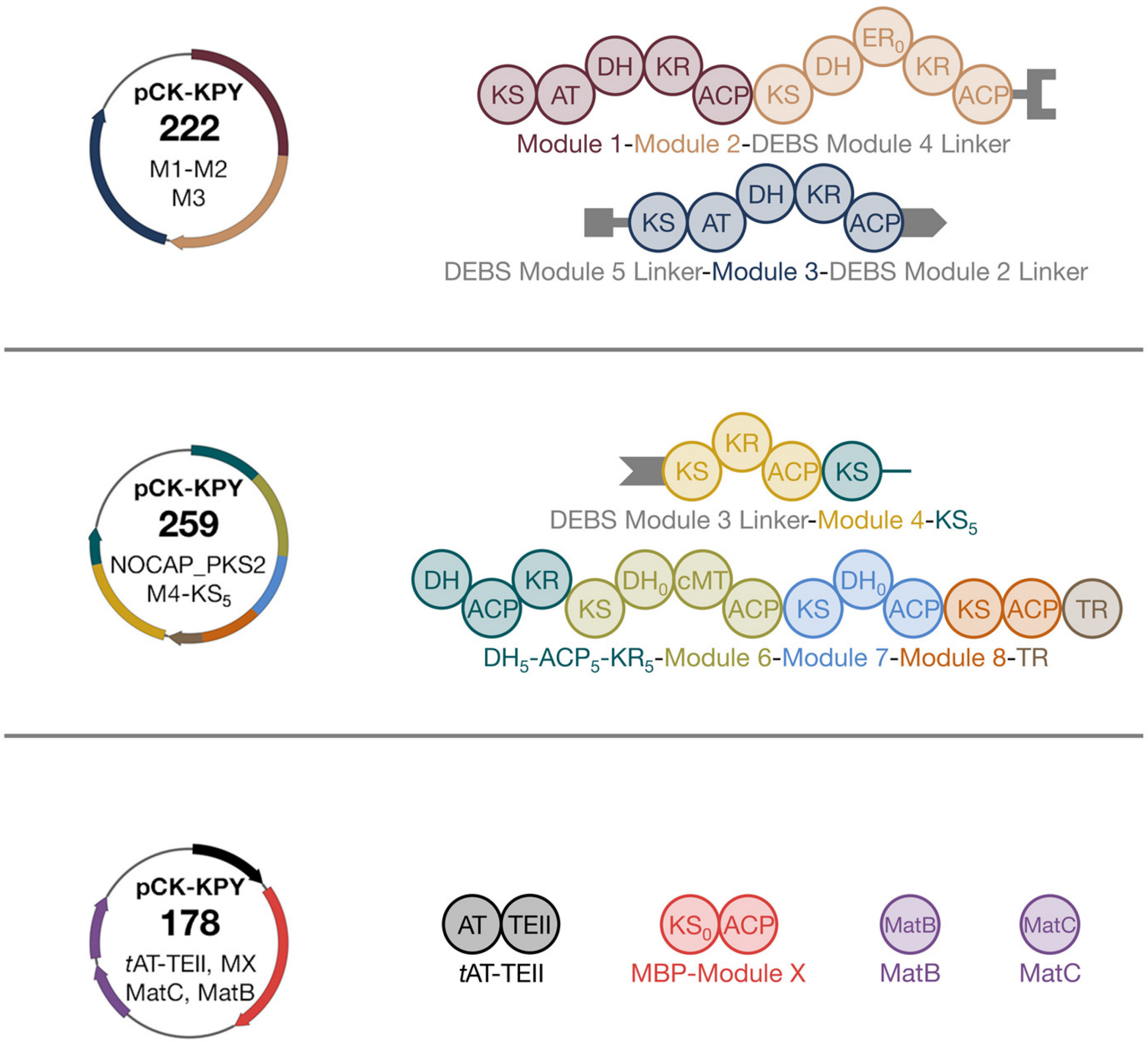

**Figure S17**. Plasmids used for the heterologous expression of the NOCAP synthase in *E. coli*: **pCK-KPY222**, Module 1-Module 2-DEBS Module 4 Linker and DEBS Module 5 Linker-Module 3-DEBS Module 2 Linker; **pCK-KPY259**, DEBS Module 3 Linker-Module 4-$KS_5$ and $DH_5$-$ACP_5$-$KR_5$-Module 6-Module 7-Module 8-TR; and **pCK-KPY178**, *t*AT-TEII, MBP-Module X, MatB and MatC.

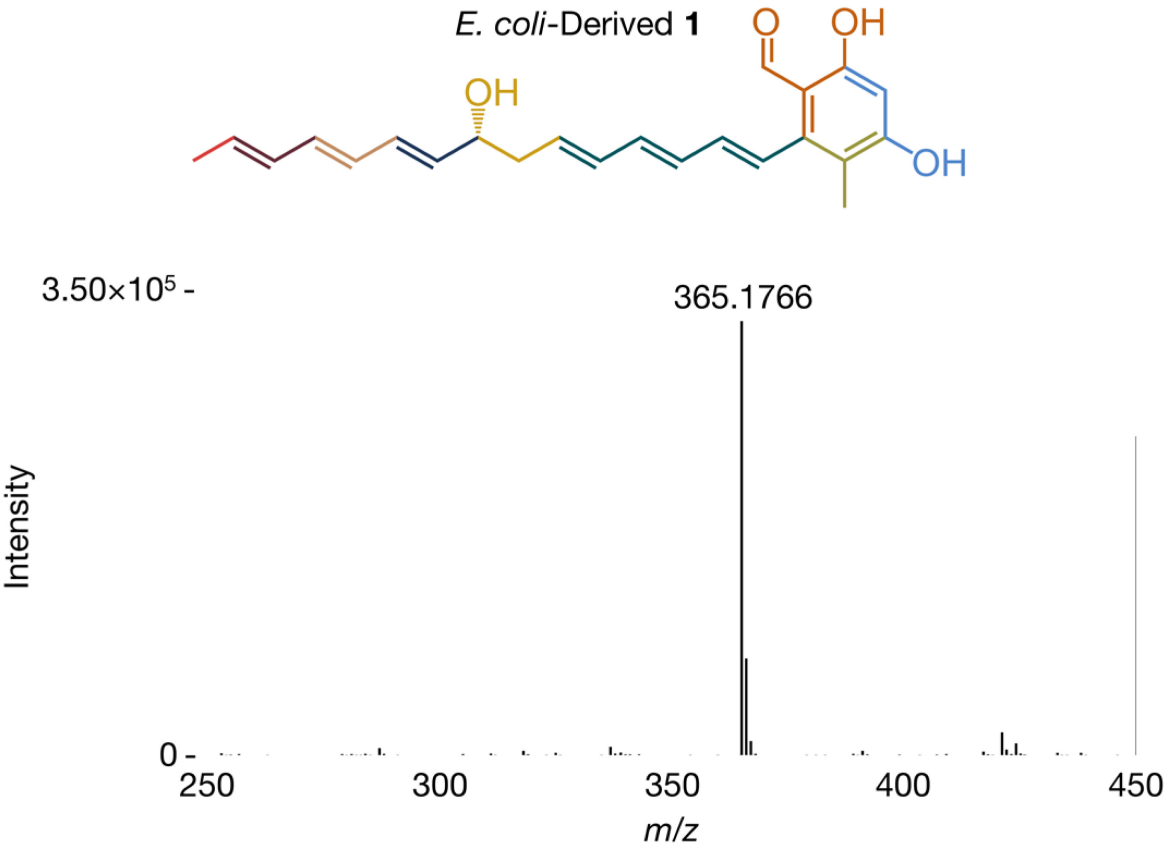

**Figure S18**. Mass spectrum (ESI-) of **1** in *E. coli* pellet extracts. Spectrum acquired on an Agilent 6545 Q-TOF LC-MS system and is representative of at least three independent experimental replicates.

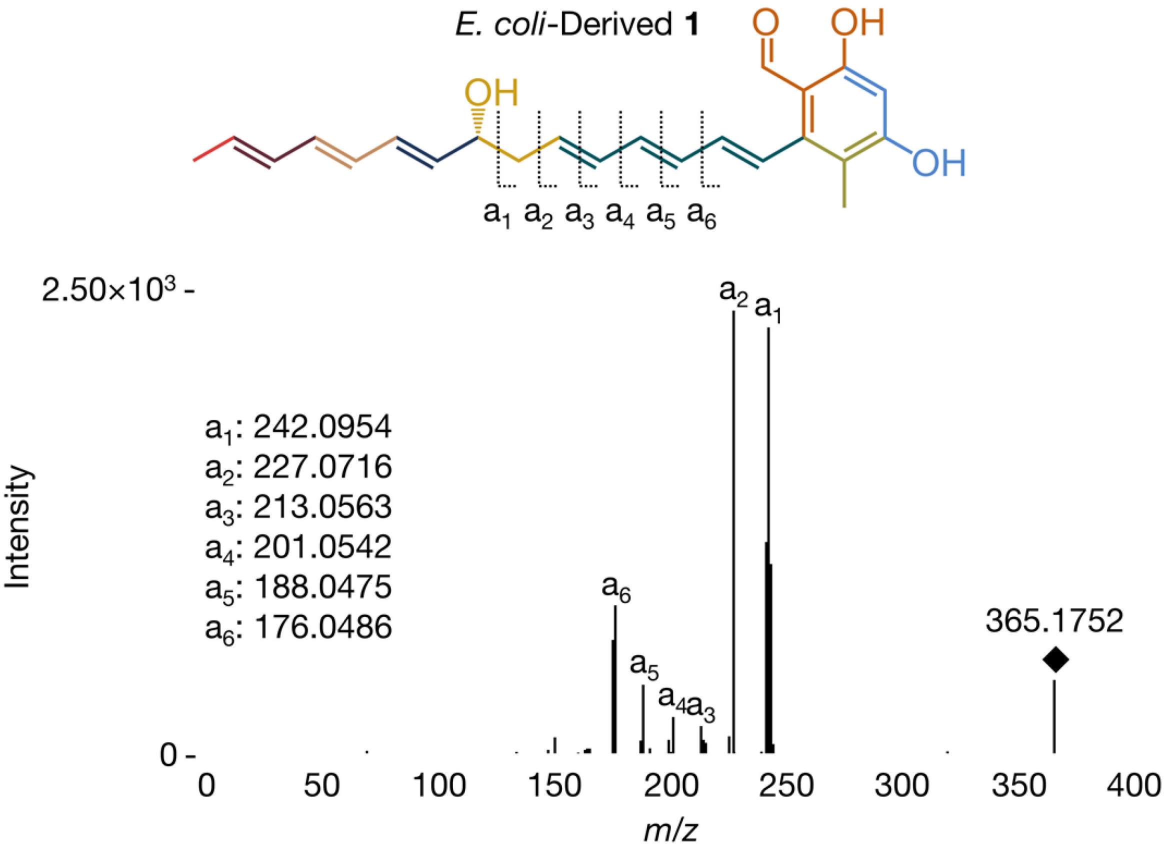

**Figure S19**. MS/MS fragmentation (ESI-) of **1** in *E. coli* pellet extracts yields six characteristic fragments. Spectrum acquired on an Agilent 6545 Q-TOF LC-MS system and is representative of at least three independent experimental replicates.

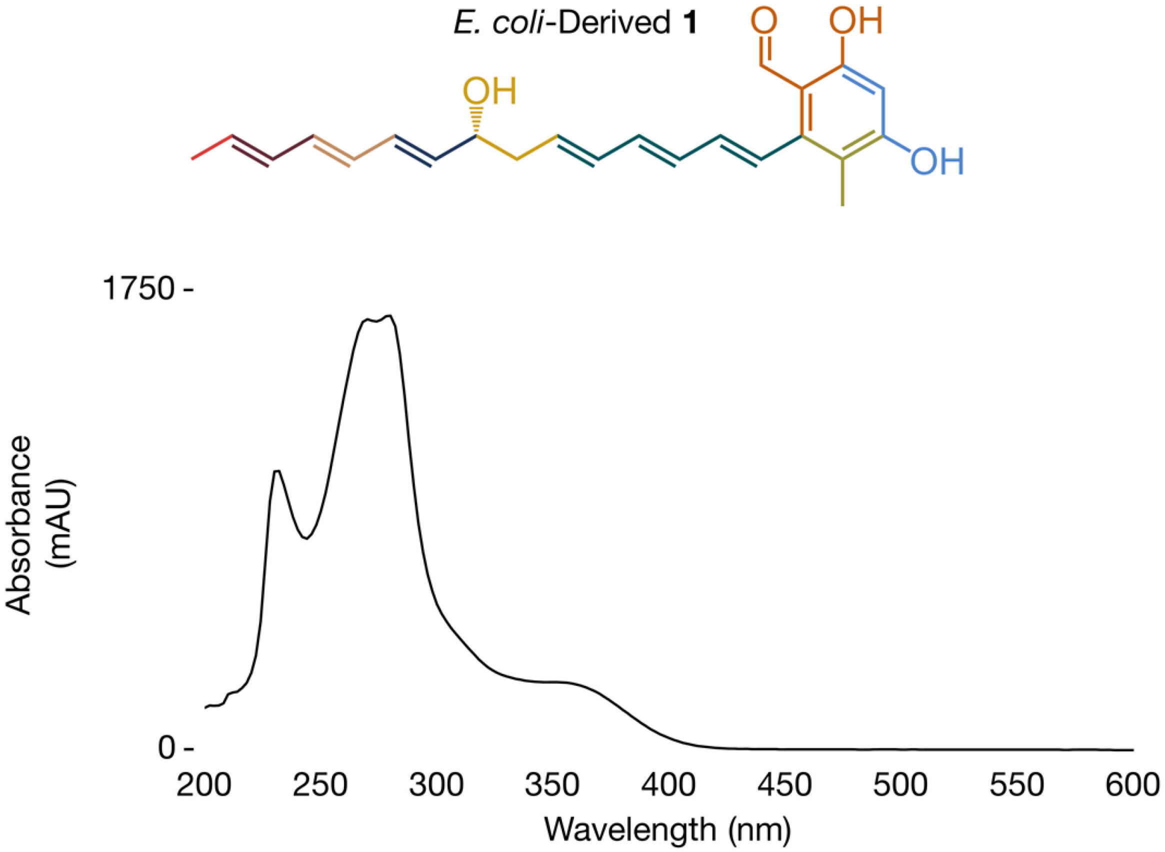

**Figure S20**. UV spectrum of **1** in *E. coli* pellet extracts. Spectrum acquired on an Agilent 6545 Q-TOF LC-MS system and is representative of at least three independent experimental replicates.

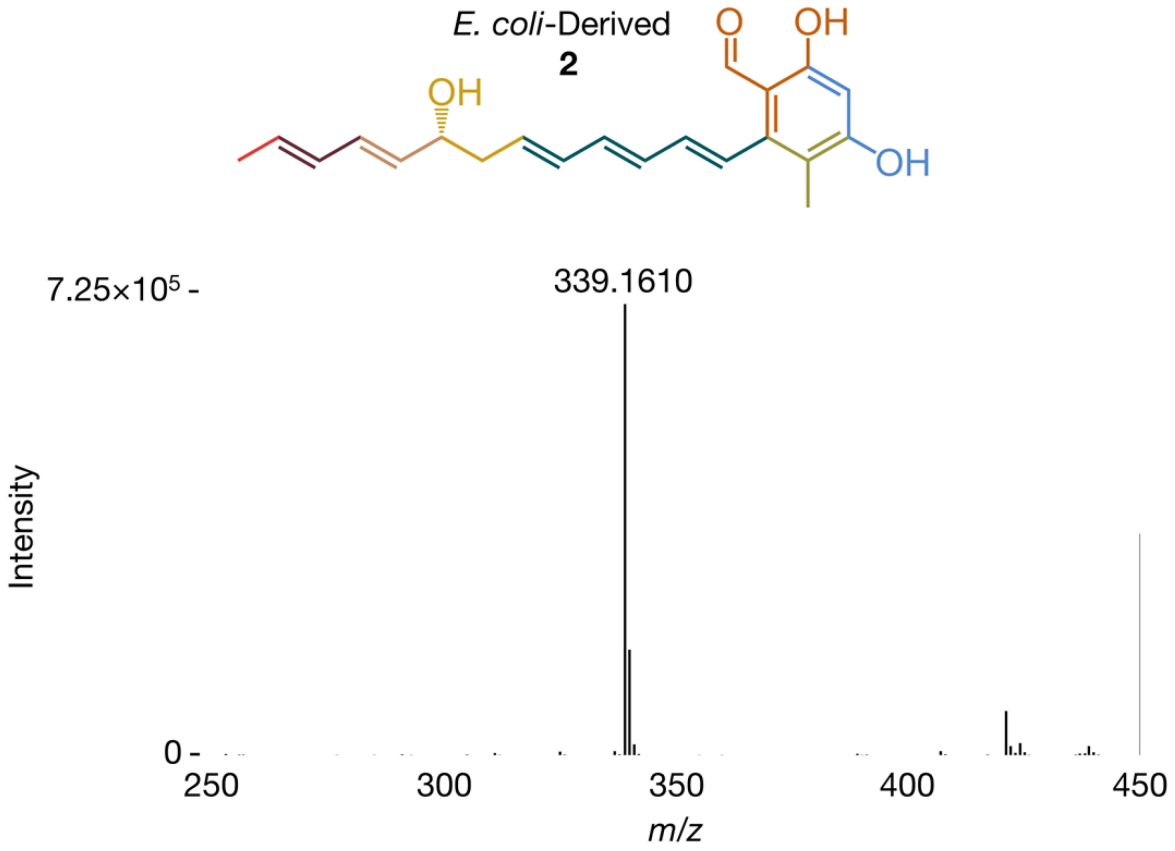

**Figure S21**. Mass spectrum (ESI-) of **2** in *E. coli* pellet extracts. Spectrum acquired on an Agilent 6545 Q-TOF LC-MS system and is representative of at least three independent experimental replicates.

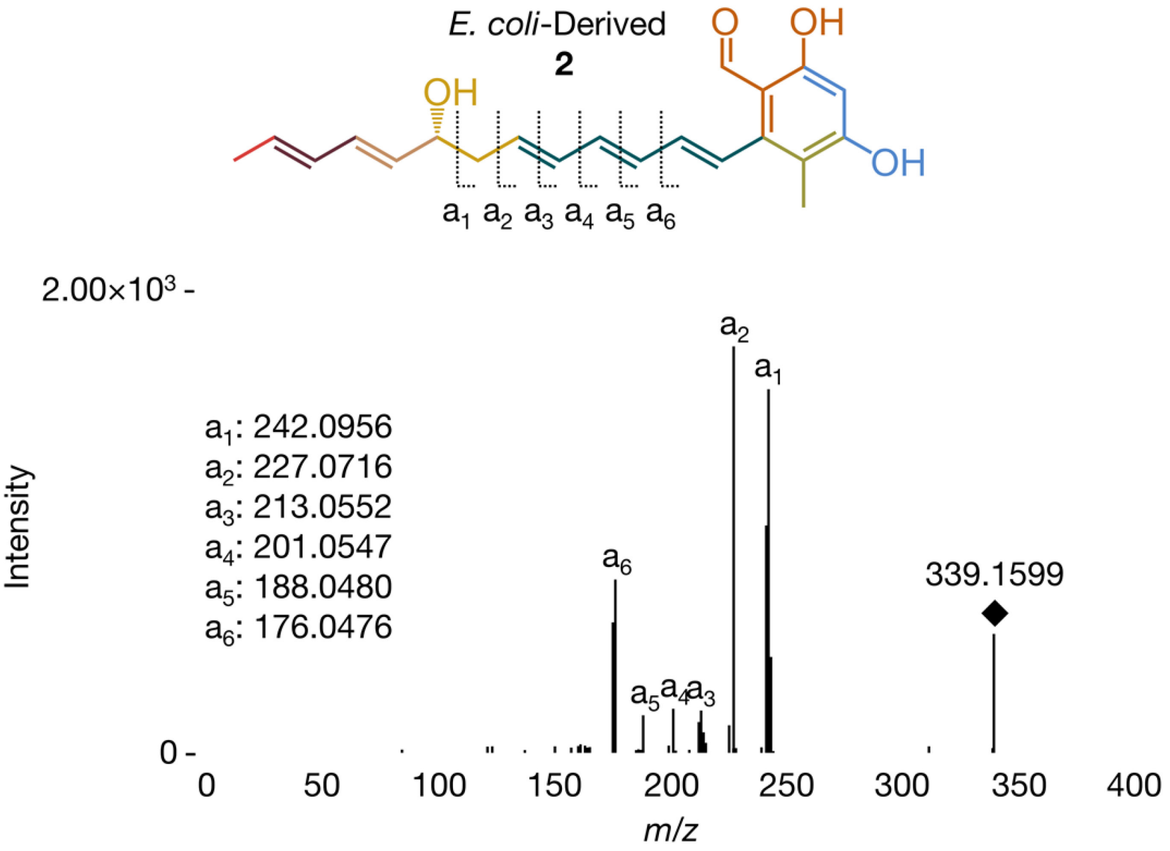

**Figure S22**. MS/MS fragmentation (ESI-) of **2** in *E. coli* pellet extracts yields six characteristic fragments. Spectrum acquired on an Agilent 6545 Q-TOF LC-MS system and is representative of at least three independent experimental replicates.

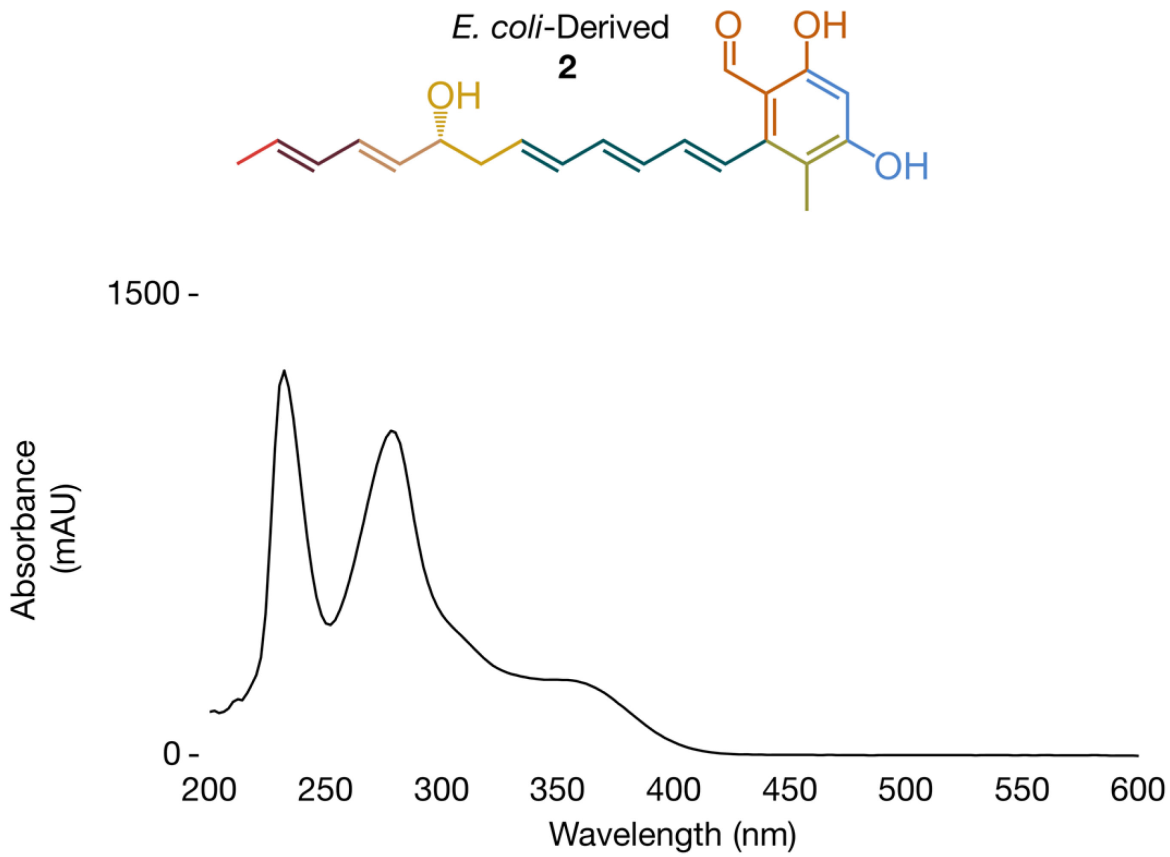

**Figure S23**. UV spectrum of **2** in *E. coli* pellet extracts. Spectrum acquired on an Agilent 6545 Q-TOF LC-MS system and is representative of at least three independent experimental replicates.

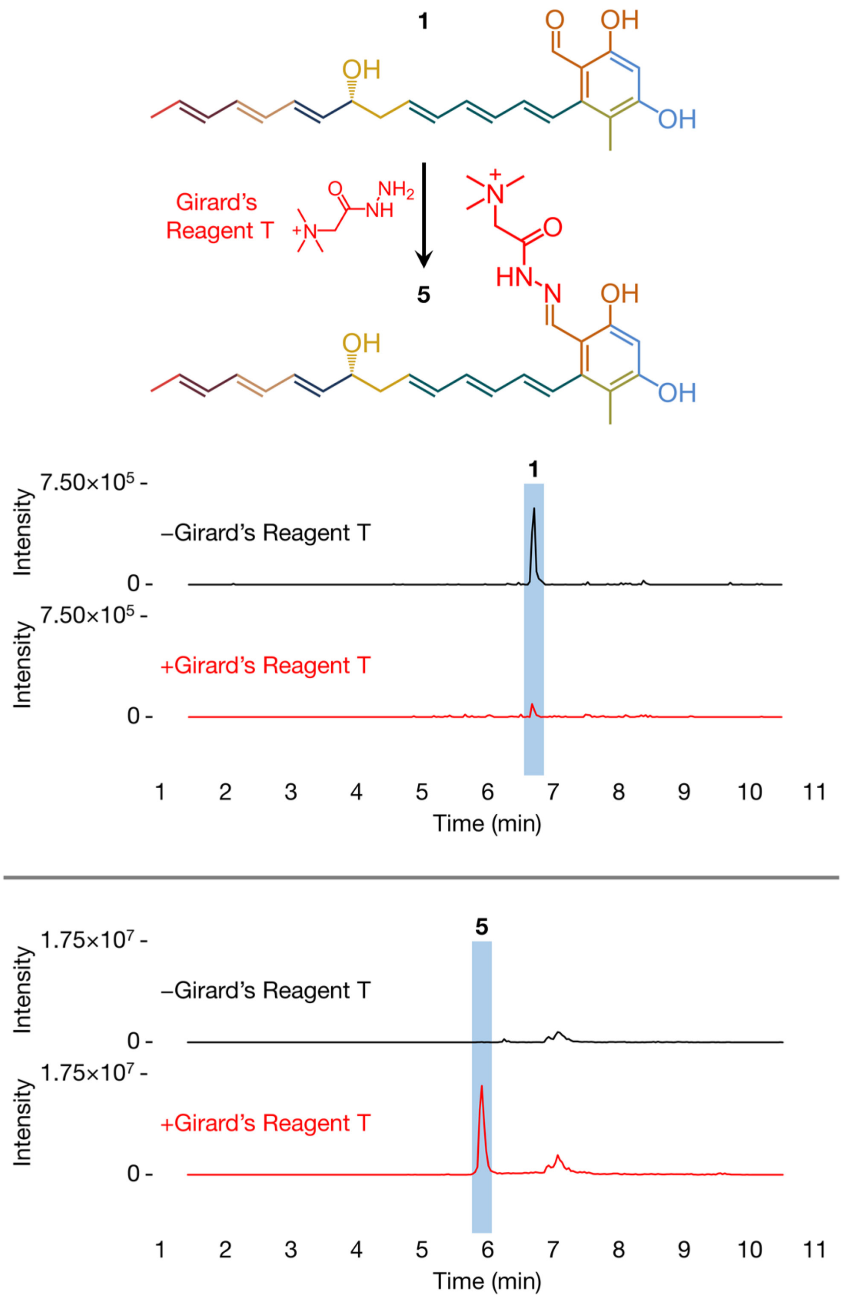

**Figure S24**. EICs (ESI-) of **1** and (ESI+) **5** in *E. coli* pellet extracts. The aldehyde-containing **1** is converted to the hydrazone-containing **5** in the presence of Girard's reagent T. Data acquired on a Waters SQ Detector 2 LC-MS system and are representative of at least three independent experimental replicates.

**5**

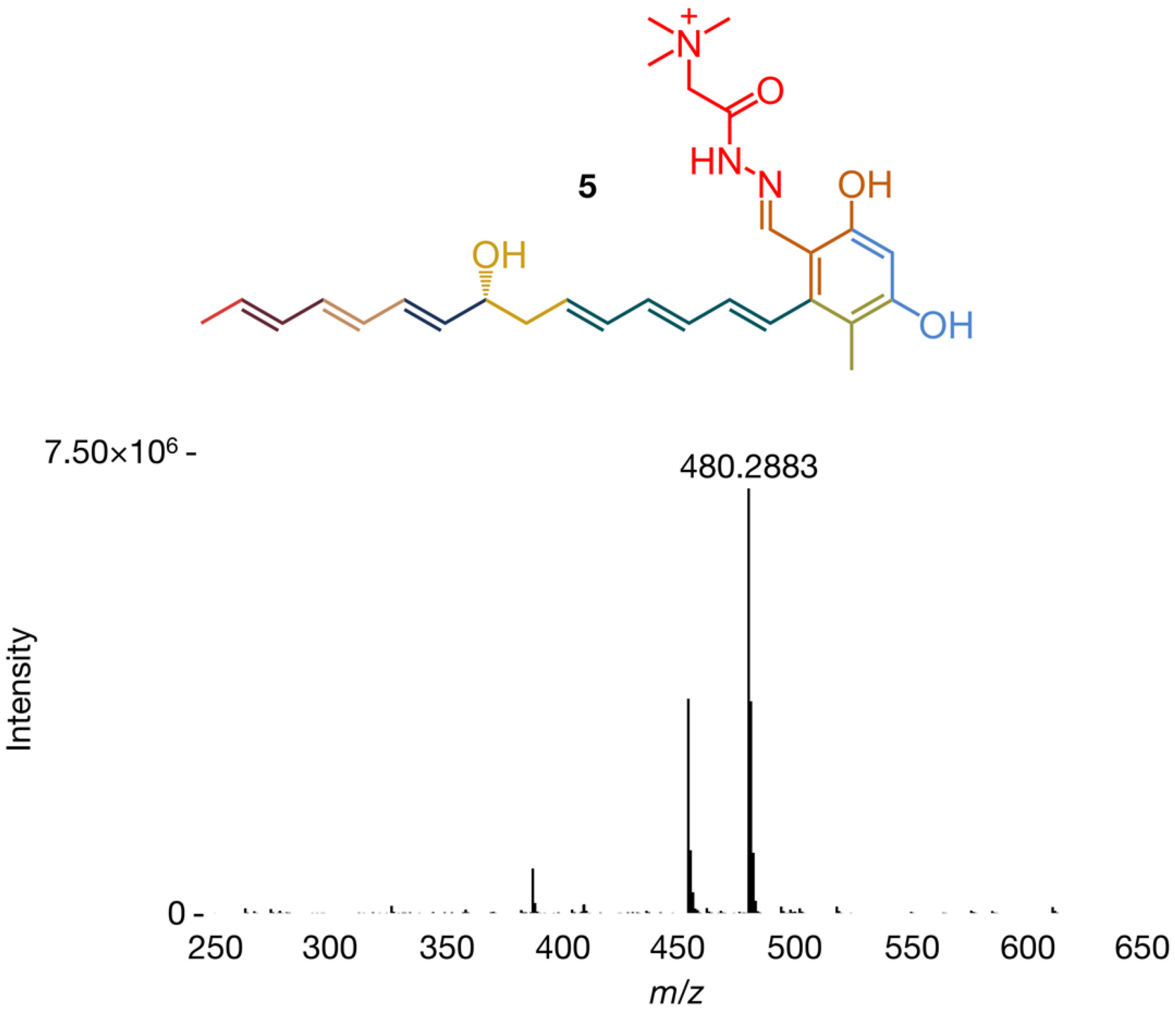

**Figure S25**. Mass spectrum (ESI+) of **5** in *E. coli* pellet extracts treated with Girard's reagent T. **5** has a molecular formula of $C_{28}H_{38}N_3O_4^+$ (observed *m/z* 480.2870, theoretical *m/z* 480.2862, 1.7 ppm). Spectrum acquired on an Agilent 6545 Q-TOF LC-MS system and is representative of at least three independent experimental replicates.

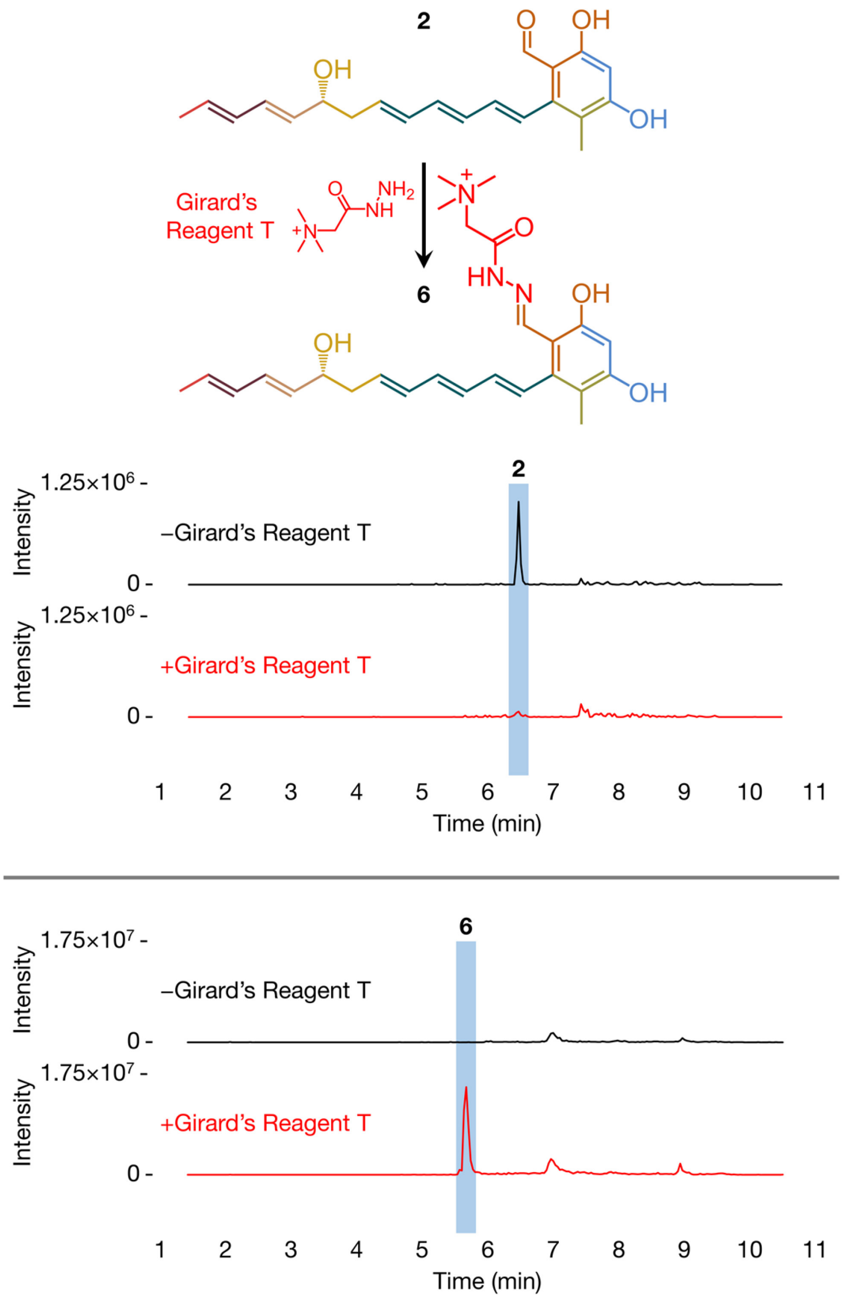

**Figure S26**. EICs (ESI-) of **2** and (ESI+) **6** in *E. coli* pellet extracts. The aldehyde-containing **2** is converted to the hydrazone-containing **6** in the presence of Girard's reagent T. Data acquired on a Waters SQ Detector 2 LC-MS system and are representative of at least three independent experimental replicates.

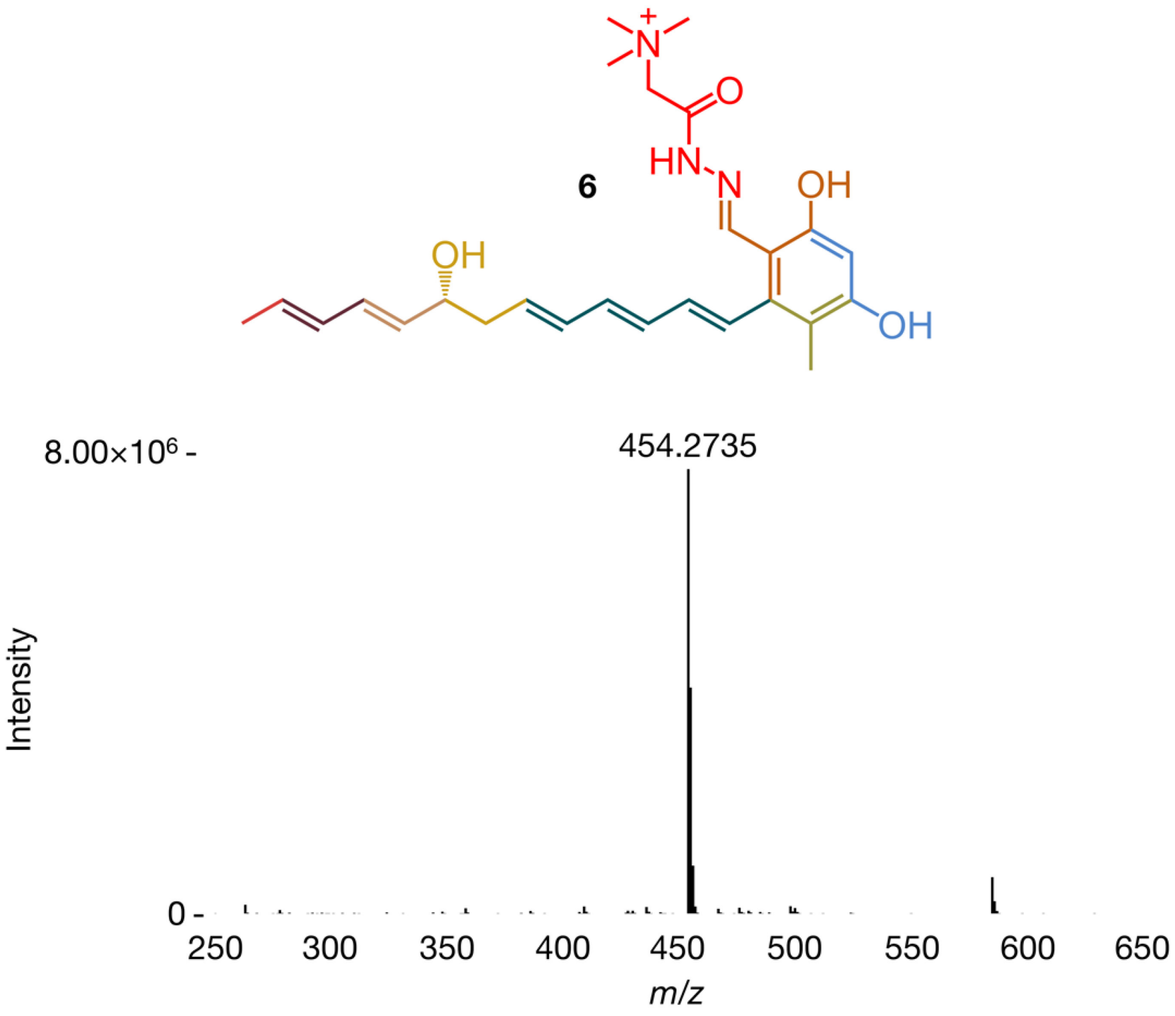

**Figure S27**. Mass spectrum (ESI+) of **6** in *E. coli* pellet extracts treated with Girard's reagent T. **6** has a molecular formula of $C_{26}H_{36}N_3O_4^+$ (observed *m/z* 454.2716, theoretical *m/z* 454.2706, 2.2 ppm). Spectrum acquired on an Agilent 6545 Q-TOF LC-MS system and is representative of at least three independent experimental replicates.

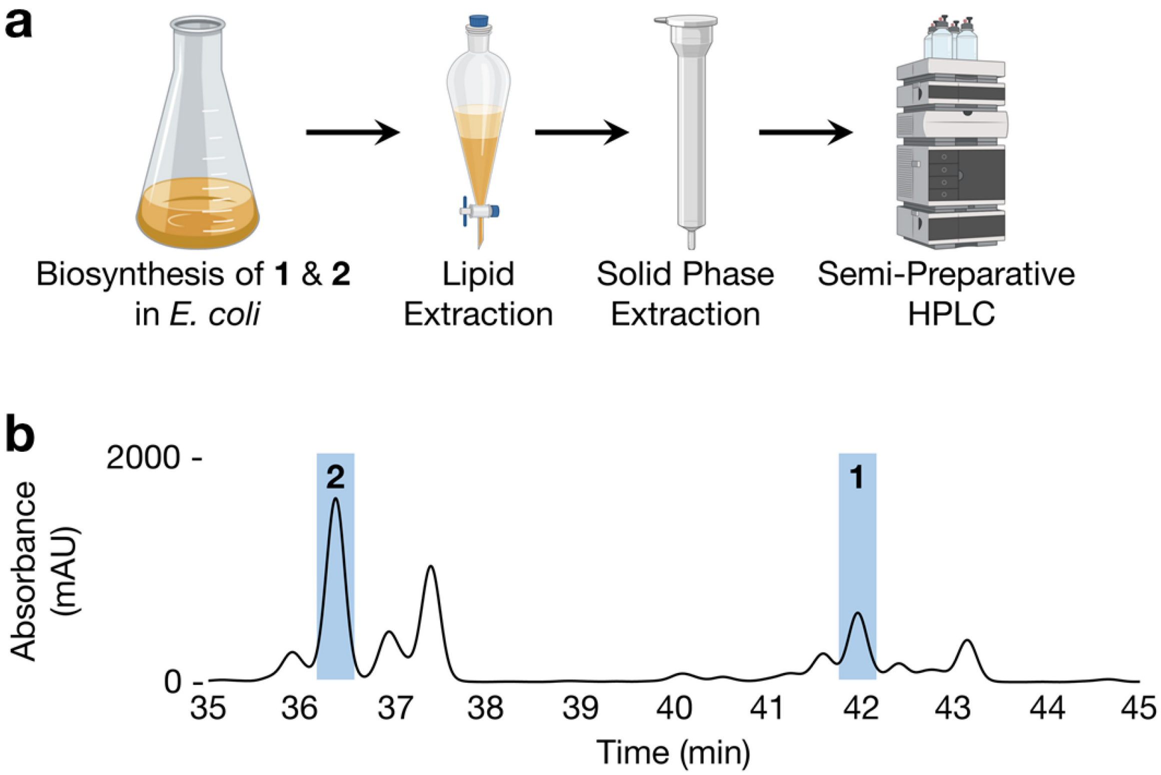

**Figure S28**. (a) Following their biosynthesis in *E. coli*, purifying **1** and **2** requires three steps: lipid extraction, solid phase extraction and semi-preparative HPLC with typical yields on the order of 1-10 mg/L culture. (b) Extracted UV chromatogram (360 nm) acquired during semi-preparative HPLC purification of **1** and **2** and is representative of at least three independent experimental replicates. **1** eluted off an Agilent Eclipse XDB-C8 column at 42.0 min (67/33 HPLC B/HPLC A); **2** eluted off this column at 36.4 min (62/38 HPLC B/HPLC A).

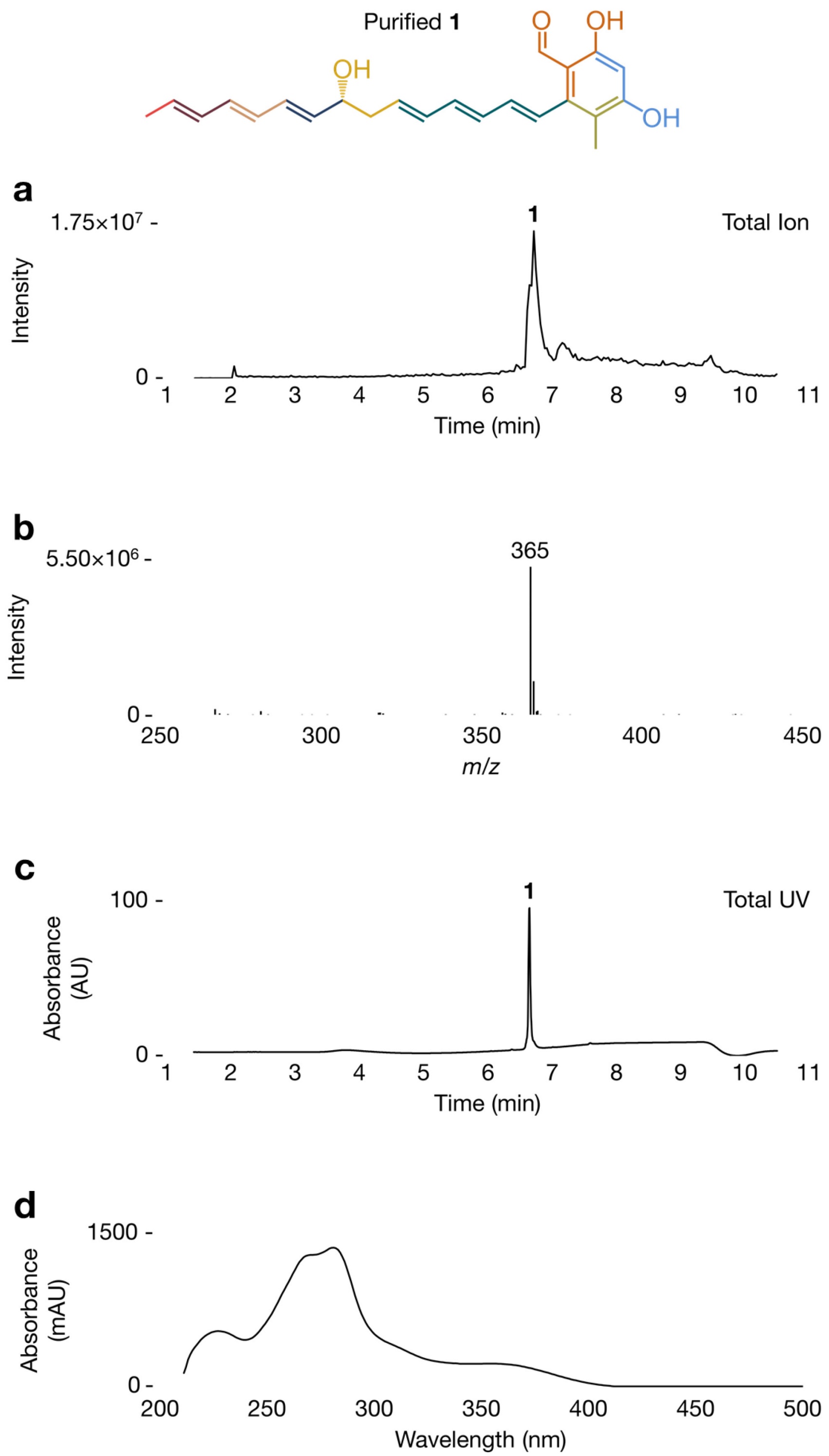

**Figure S29**. (a) Total ion chromatogram (ESI-), (b) MS spectrum (ESI-), (c) total UV chromatogram and (d) UV spectrum of purified **1**. Data acquired on a Waters SQ Detector 2 LC-MS system and are representative of at least three independent experimental replicates.

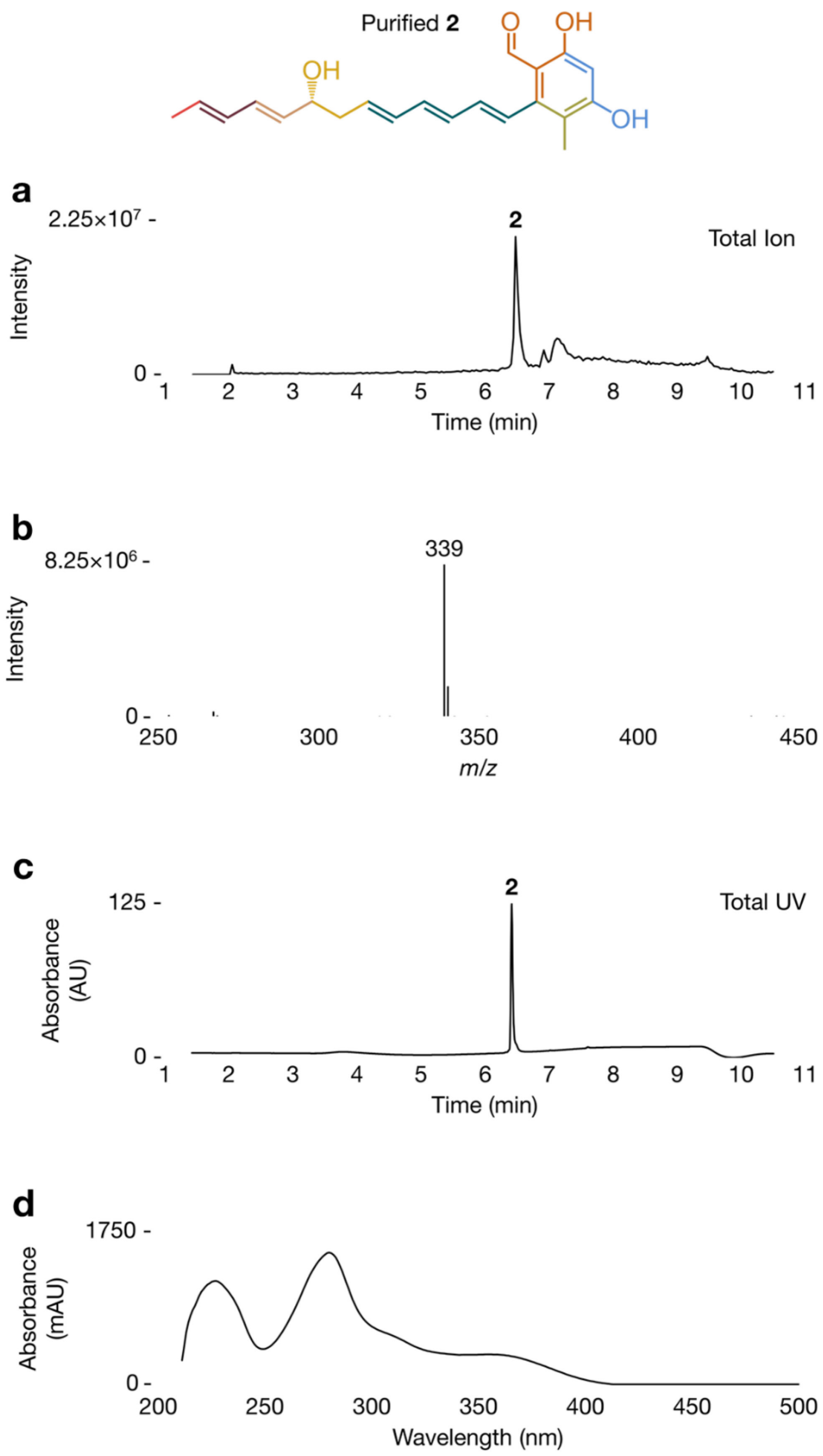

**Figure S30**. (a) Total ion chromatogram (ESI-), (b) MS spectrum (ESI-), (c) total UV chromatogram and (d) UV spectrum of purified **2**. Data acquired on a Waters SQ Detector 2 LC-MS system and are representative of at least three independent experimental replicates.

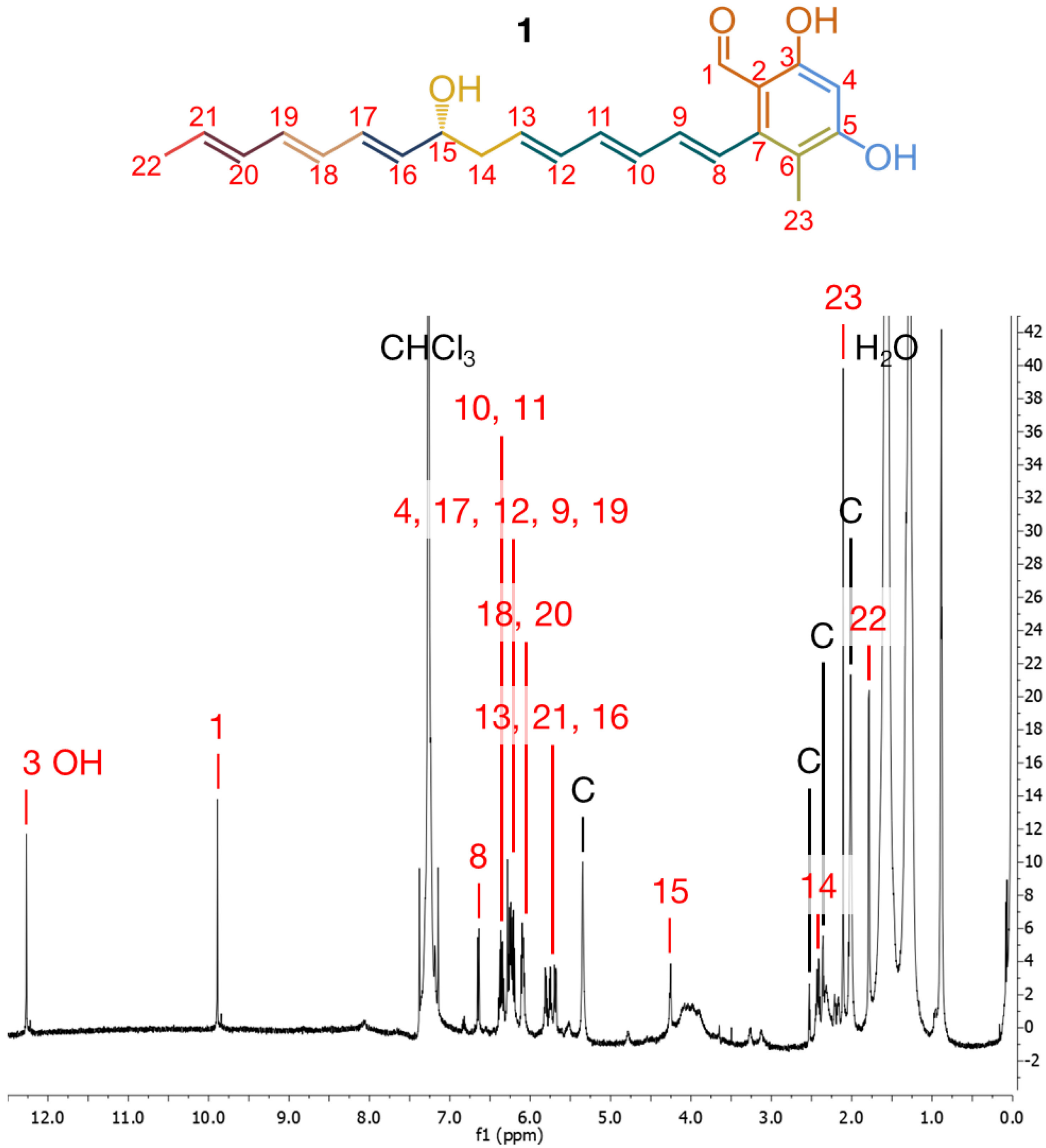

**Figure S31**. [1]H NMR spectrum (x-axis: δH, 12.5-0.0 ppm) of **1** in $CDCl_3$. Key: C, contaminant. Spectrum acquired on a 900 MHz Bruker AVANCE II spectrometer.

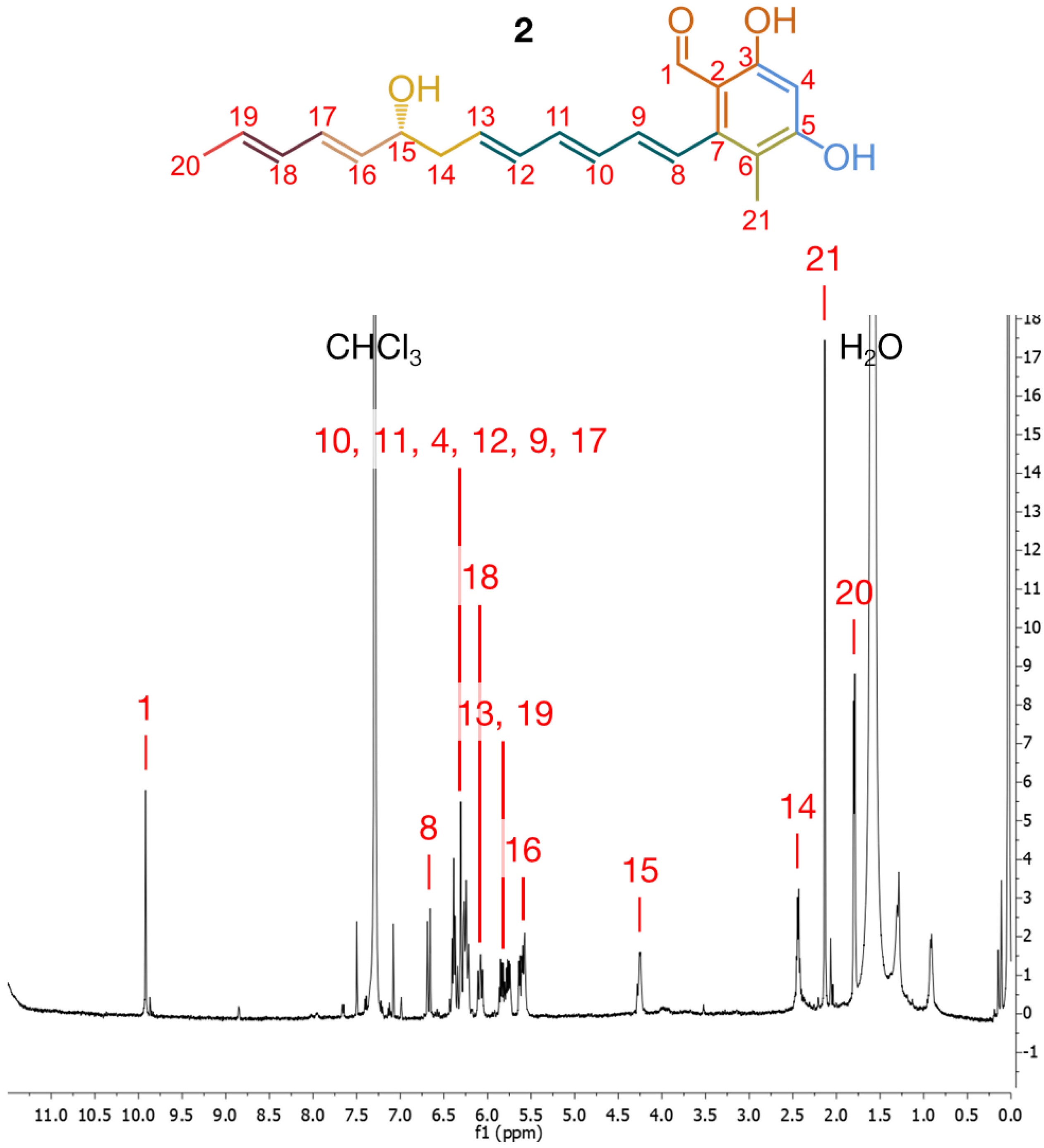

**Figure S32**. $^{1}H$ NMR spectrum (x-axis: δH, 11.5-0.0 ppm) of **2** in $CDCl_3$. Spectrum acquired on a 500 MHz Bruker AVANCE spectrometer.

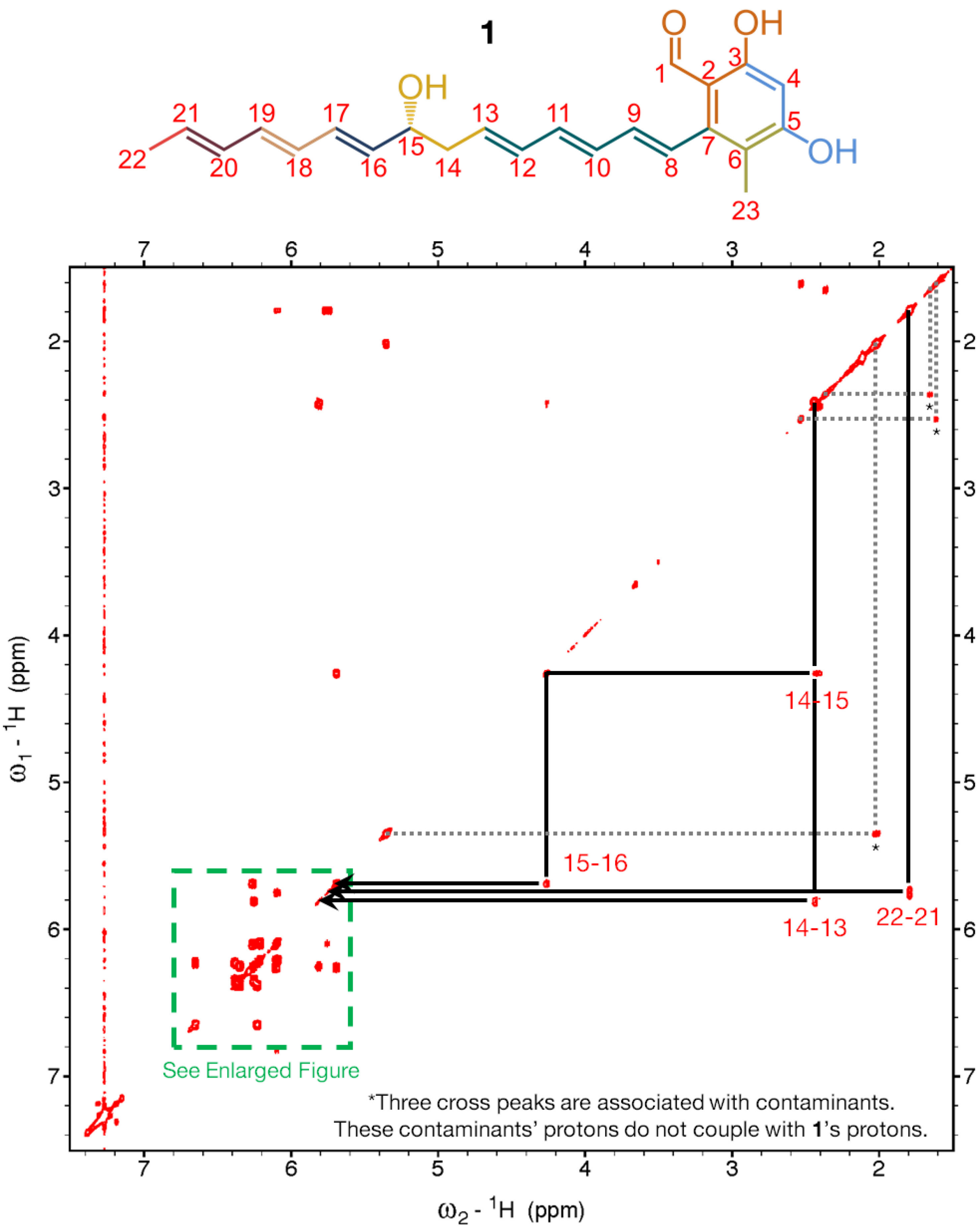

**Figure S33**. $^1$H-$^1$H COSY spectrum (positive signals in red; negative signals in blue; x-axis: δH, 7.5-1.5 ppm; y-axis: δH, 7.5-1.5 ppm) of **1** in $CDCl_3$. Spectrum acquired on a 900 MHz Bruker AVANCE II spectrometer.

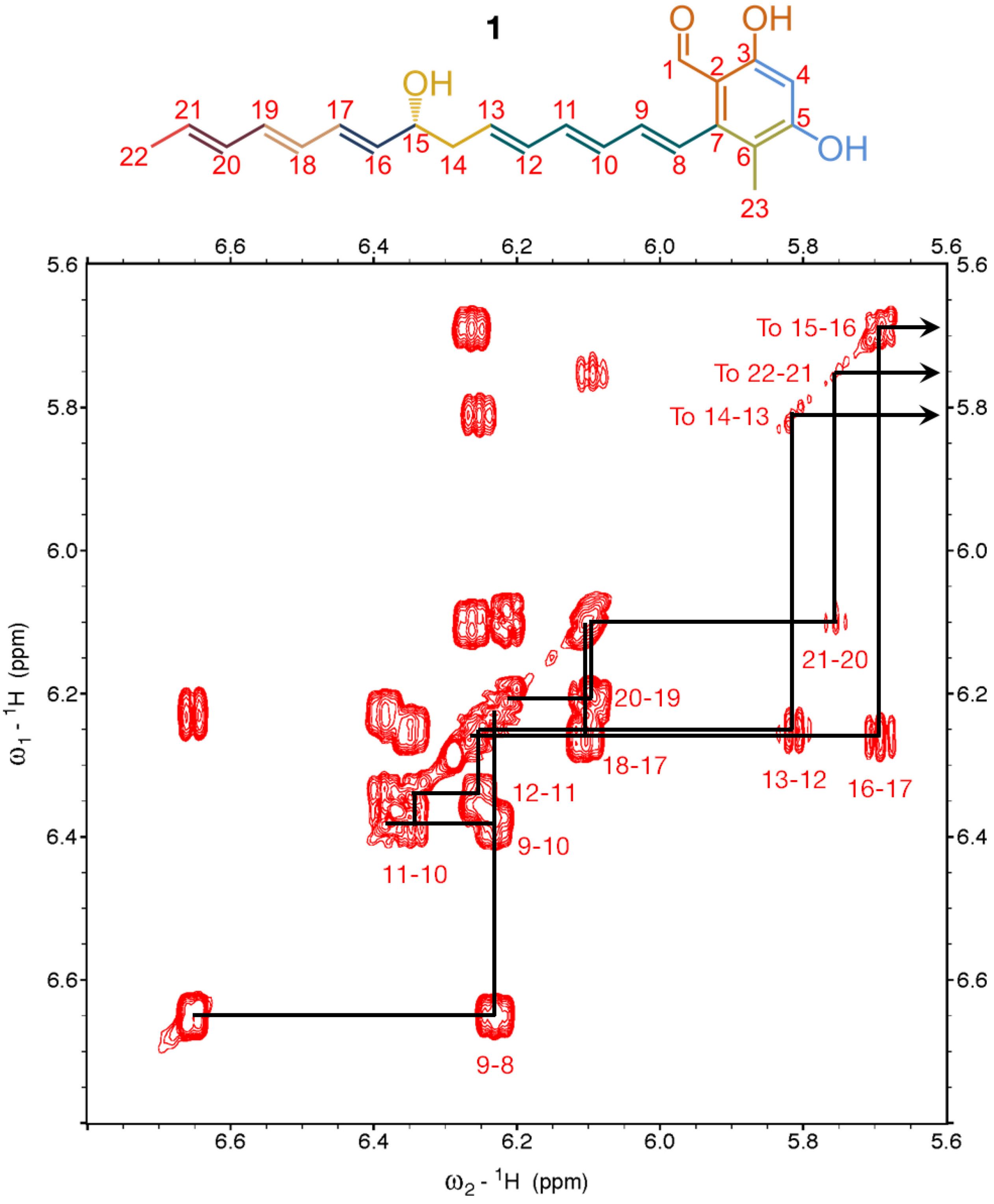

**Figure S34**. $^1H$-$^1H$ COSY spectrum (positive signals in red; negative signals in blue; x-axis: δH, 6.8-5.6 ppm; y-axis: δH, 6.8-5.6 ppm) of **1** in $CDCl_3$. Spectrum acquired on a 900 MHz Bruker AVANCE II spectrometer.

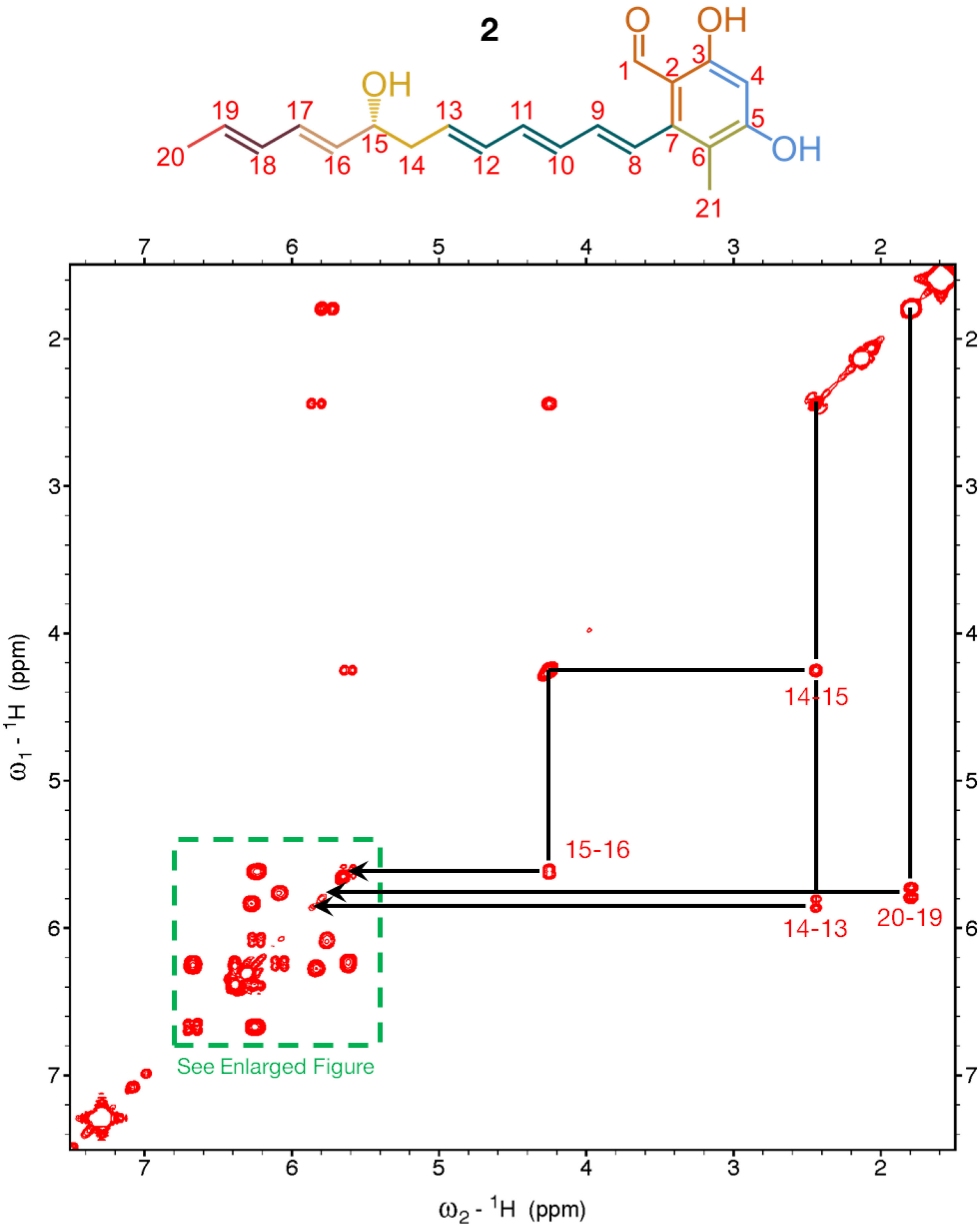

**Figure S35**. $^1H$-$^1H$ COSY spectrum (positive signals in red; negative signals in blue; x-axis: δH, 7.5-1.5 ppm; y-axis: δH, 7.5-1.5 ppm) of **2** in $CDCl_3$. Spectrum acquired on a 500 MHz Bruker AVANCE spectrometer.

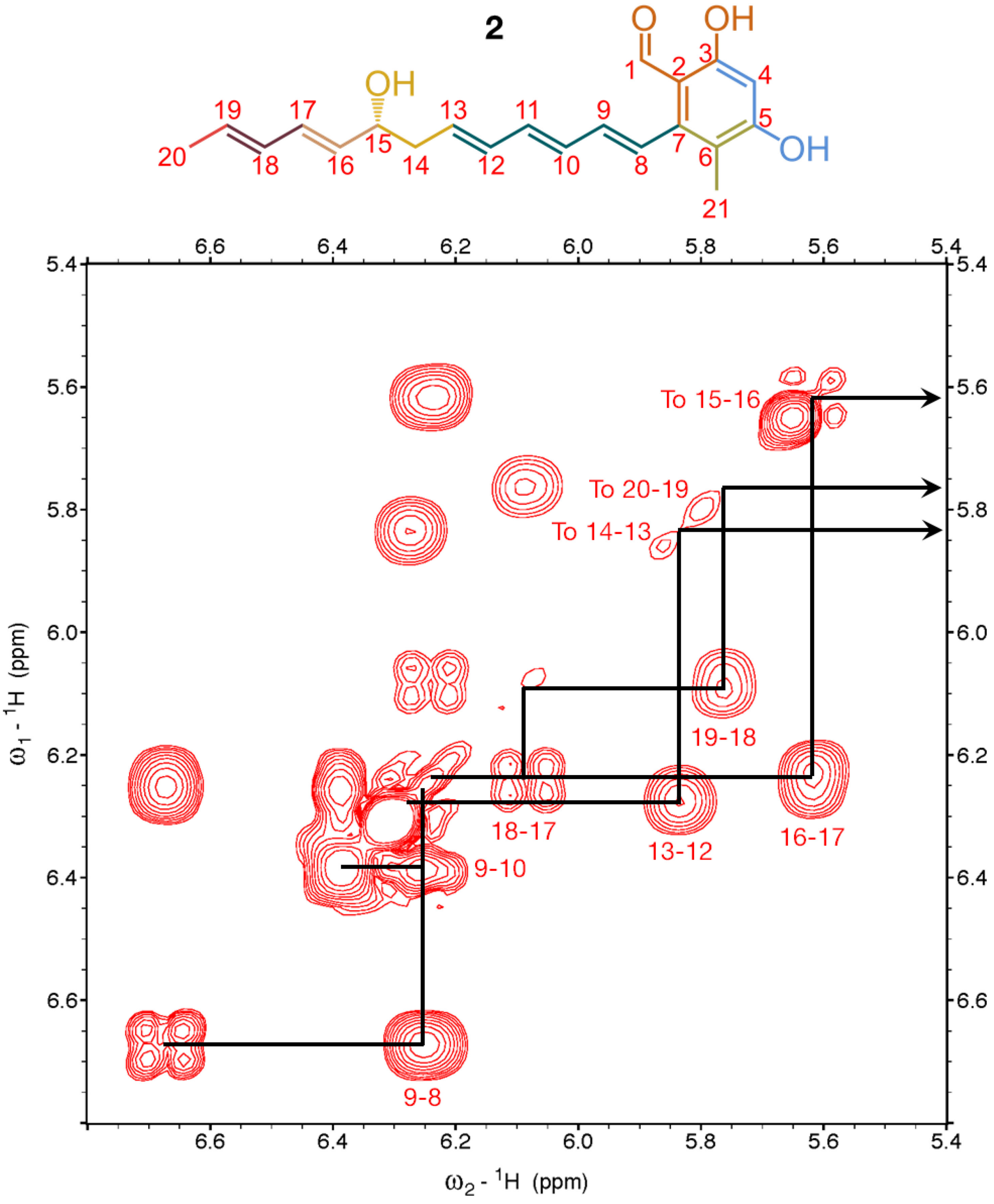

**Figure S36**. ¹H-¹H COSY spectrum (positive signals in red; negative signals in blue; x-axis: δH, 6.8-5.4 ppm; y-axis: δH, 6.8-5.4 ppm) of **2** in $CDCl_3$. Spectrum acquired on a 500 MHz Bruker AVANCE spectrometer.

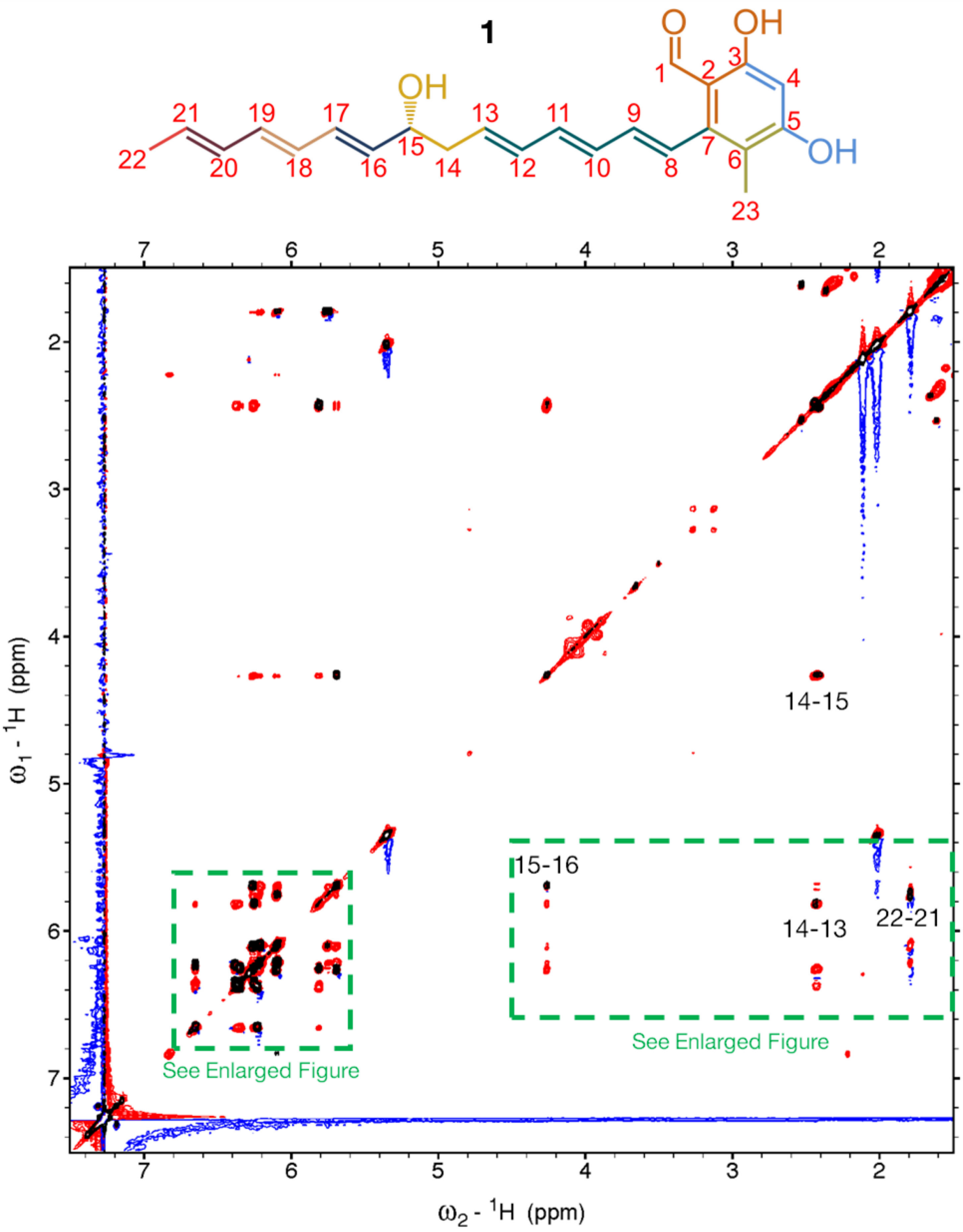

**Figure S37**. $^1H$-$^1H$ TOCSY (COSY overlaid in black for positive signals and orange for negative signals) spectrum (positive signals in red; negative signals in blue; x-axis: δH, 7.5-1.5 ppm; y-axis: δH, 7.5-1.5 ppm) of **1** in $CDCl_3$. Spectrum acquired on a 900 MHz Bruker AVANCE II spectrometer.

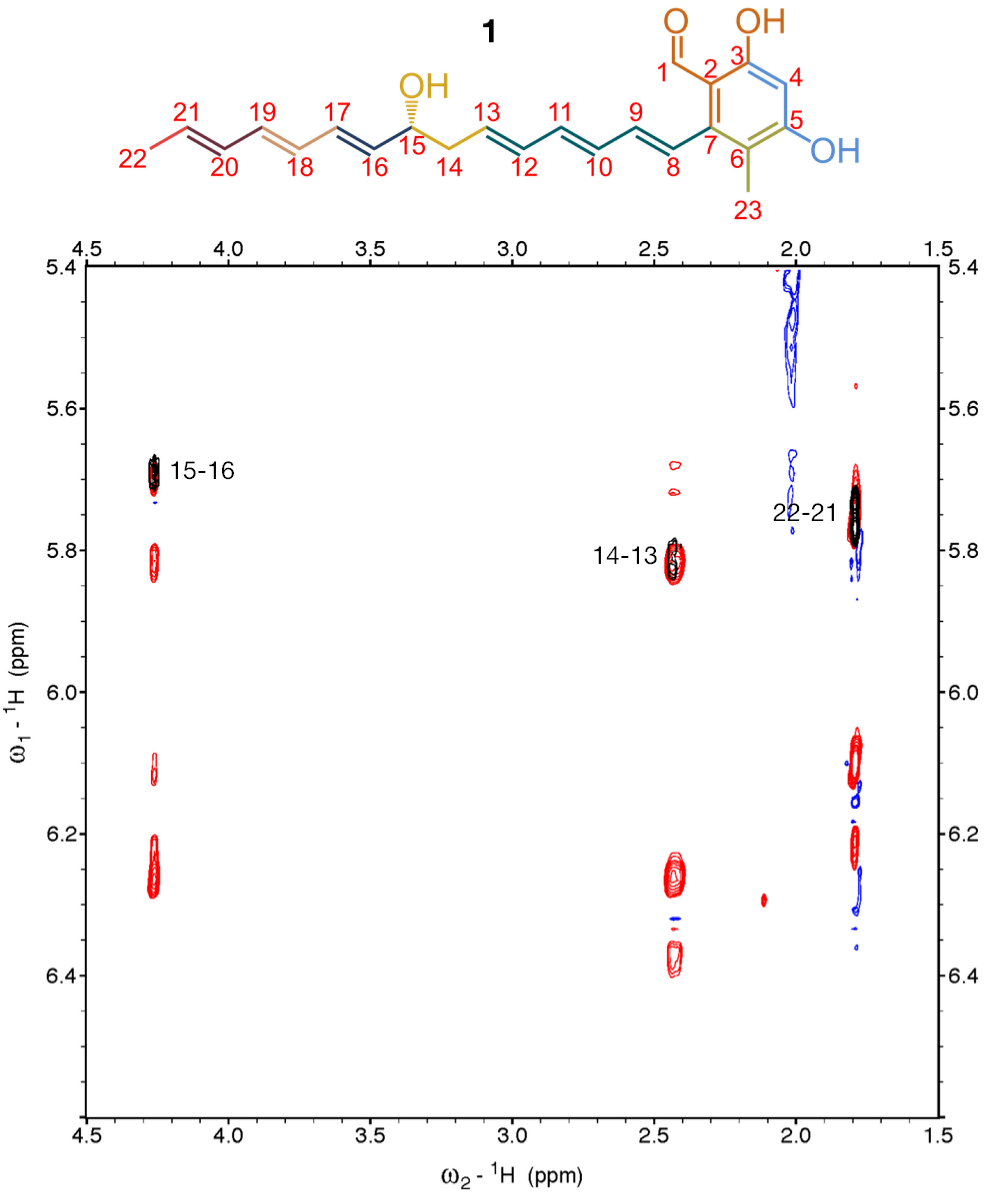

**Figure S38**. $^1$H-$^1$H TOCSY (COSY overlaid in black for positive signals and orange for negative signals) spectrum (positive signals in red; negative signals in blue; x-axis: δH, 4.5-1.5 ppm; y-axis: δH, 6.6-5.4 ppm) of **1** in $CDCl_3$. Spectrum acquired on a 900 MHz Bruker AVANCE II spectrometer.

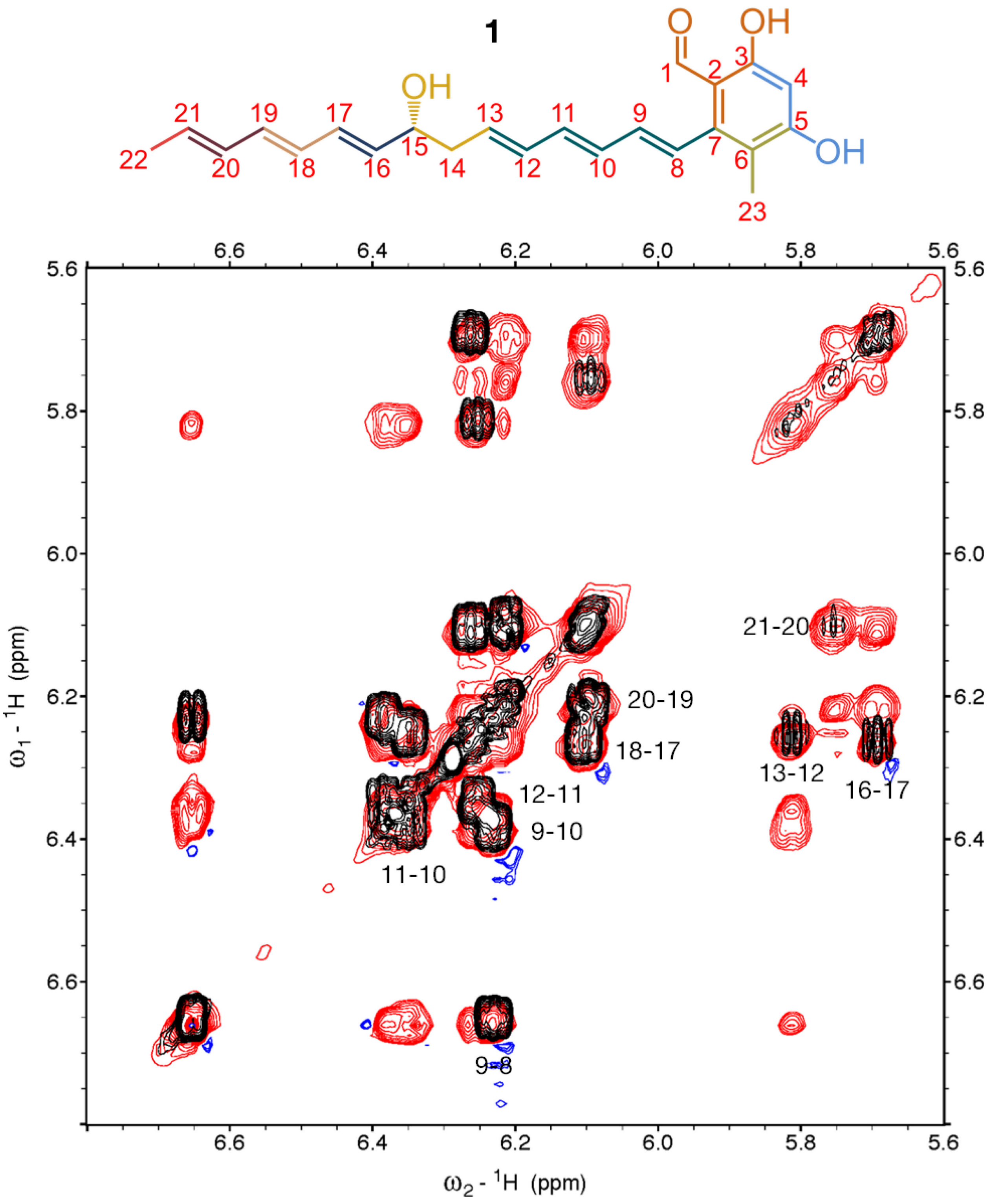

**Figure S39**. $^1H$-$^1H$ TOCSY (COSY overlaid in black for positive signals and orange for negative signals) spectrum (positive signals in red; negative signals in blue; x-axis: δH, 6.8-5.6 ppm; y-axis: δH, 6.8-5.6 ppm) of **1** in $CDCl_3$. Spectrum acquired on a 900 MHz Bruker AVANCE II spectrometer.

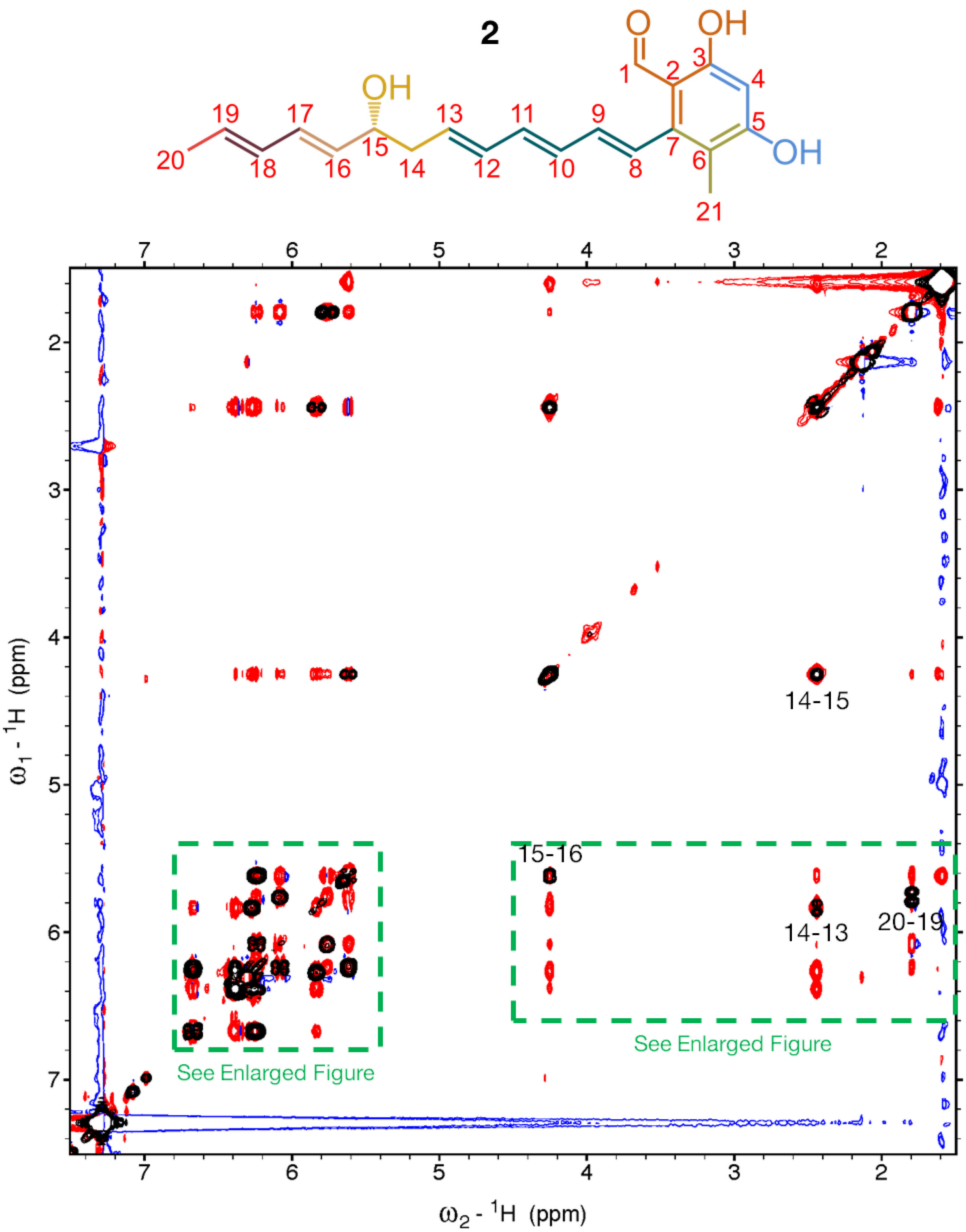

**Figure S40**. $^1$H-$^1$H TOCSY (COSY overlaid in black for positive signals and orange for negative signals) spectrum (positive signals in red; negative signals in blue; x-axis: δH, 7.5-1.5 ppm; y-axis: δH, 7.5-1.5 ppm) of **2** in $CDCl_3$. Spectrum acquired on a 500 MHz Bruker AVANCE spectrometer.

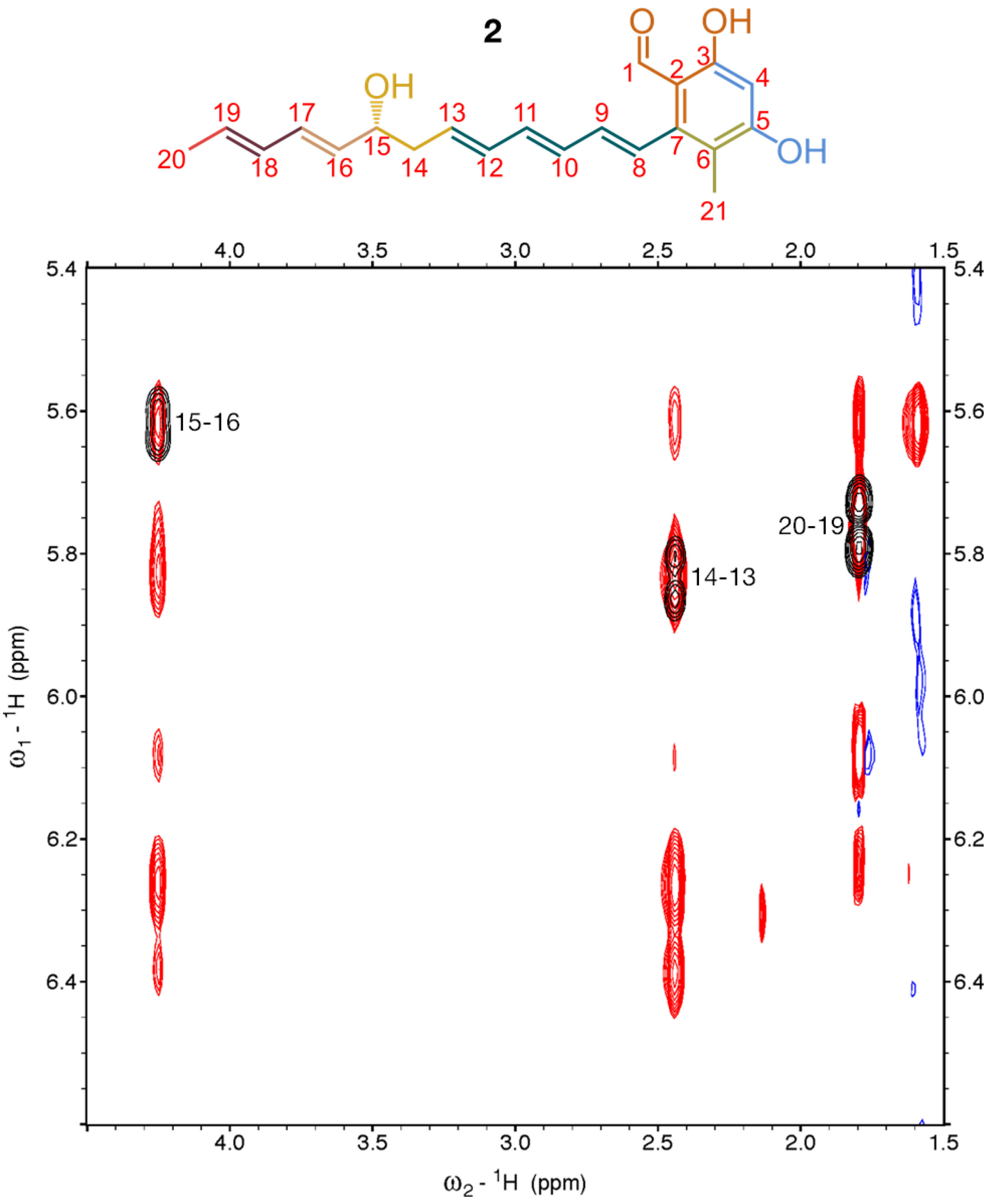

**Figure S41**. $^1$H-$^1$H TOCSY (COSY overlaid in black for positive signals and orange for negative signals) spectrum (positive signals in red; negative signals in blue; x-axis: δH, 4.5-1.5 ppm; y-axis: δH, 6.6-5.4 ppm) of **2** in $CDCl_3$. Spectrum acquired on a 500 MHz Bruker AVANCE spectrometer.

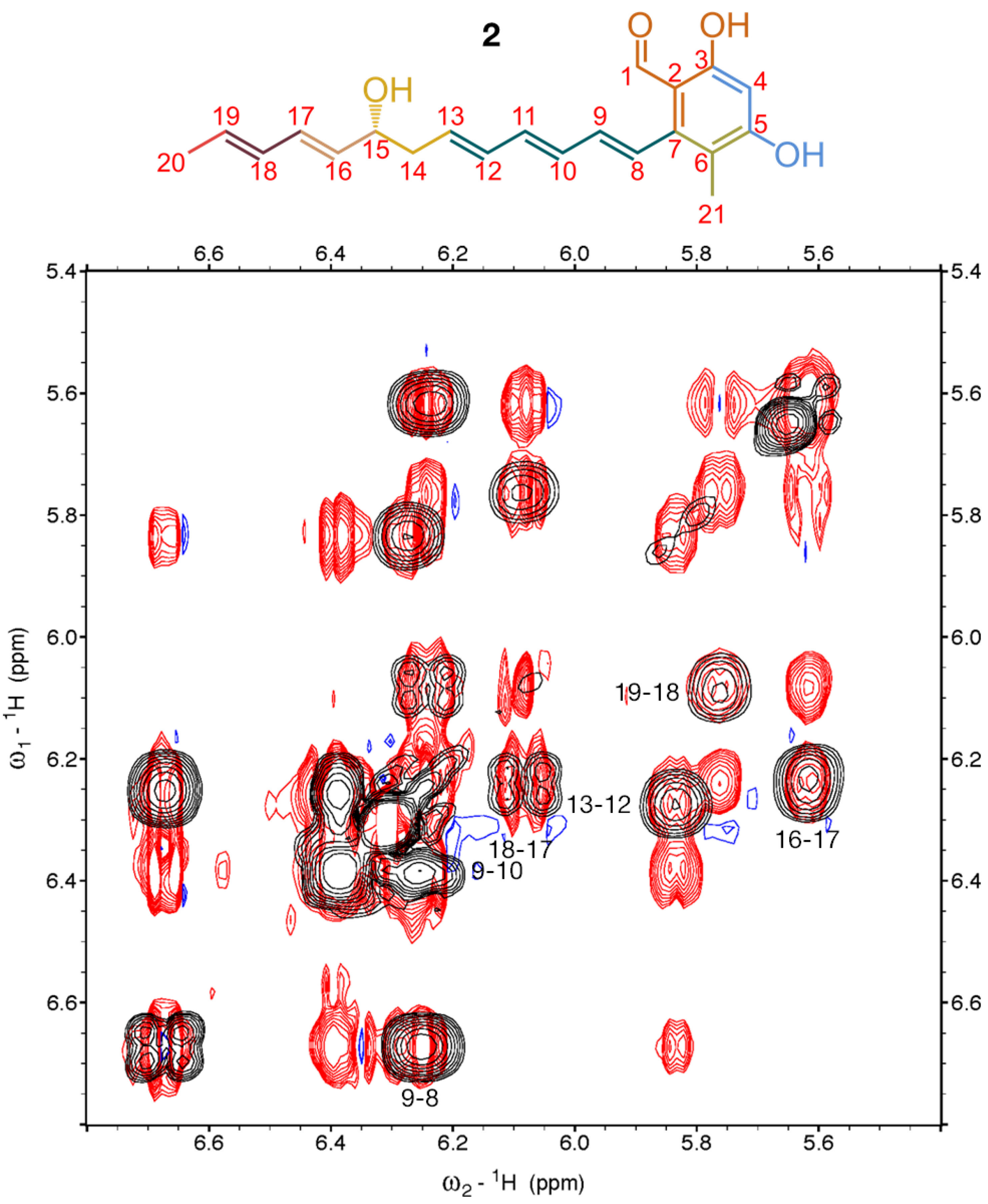

**Figure S42**. $^1H$-$^1H$ TOCSY (COSY overlaid in black for positive signals and orange for negative signals) spectrum (positive signals in red; negative signals in blue; x-axis: δH, 6.8-5.4 ppm; y-axis: δH, 6.8-5.4 ppm) of **2** in $CDCl_3$. Spectrum acquired on a 500 MHz Bruker AVANCE spectrometer.

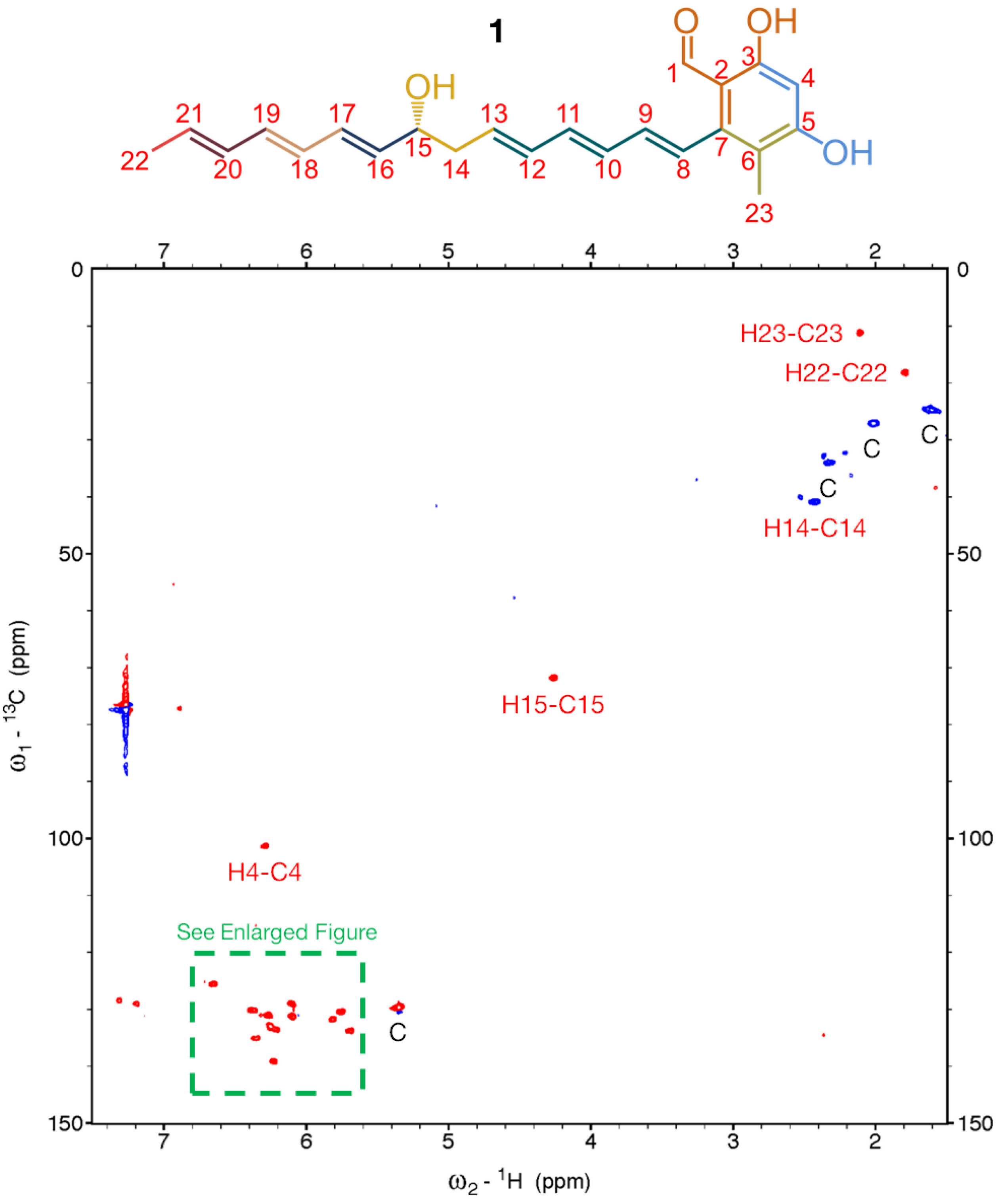

**Figure S43**. $^{1}H$-$^{13}C$ HSQC spectrum (positive signals in red; negative signals in blue; x-axis: δH, 7.5-1.5 ppm; y-axis: δC, 150.0-0.0 ppm) of **1** in $CDCl_3$. Key: C, contaminant. Spectrum acquired on a 900 MHz Bruker AVANCE II spectrometer.

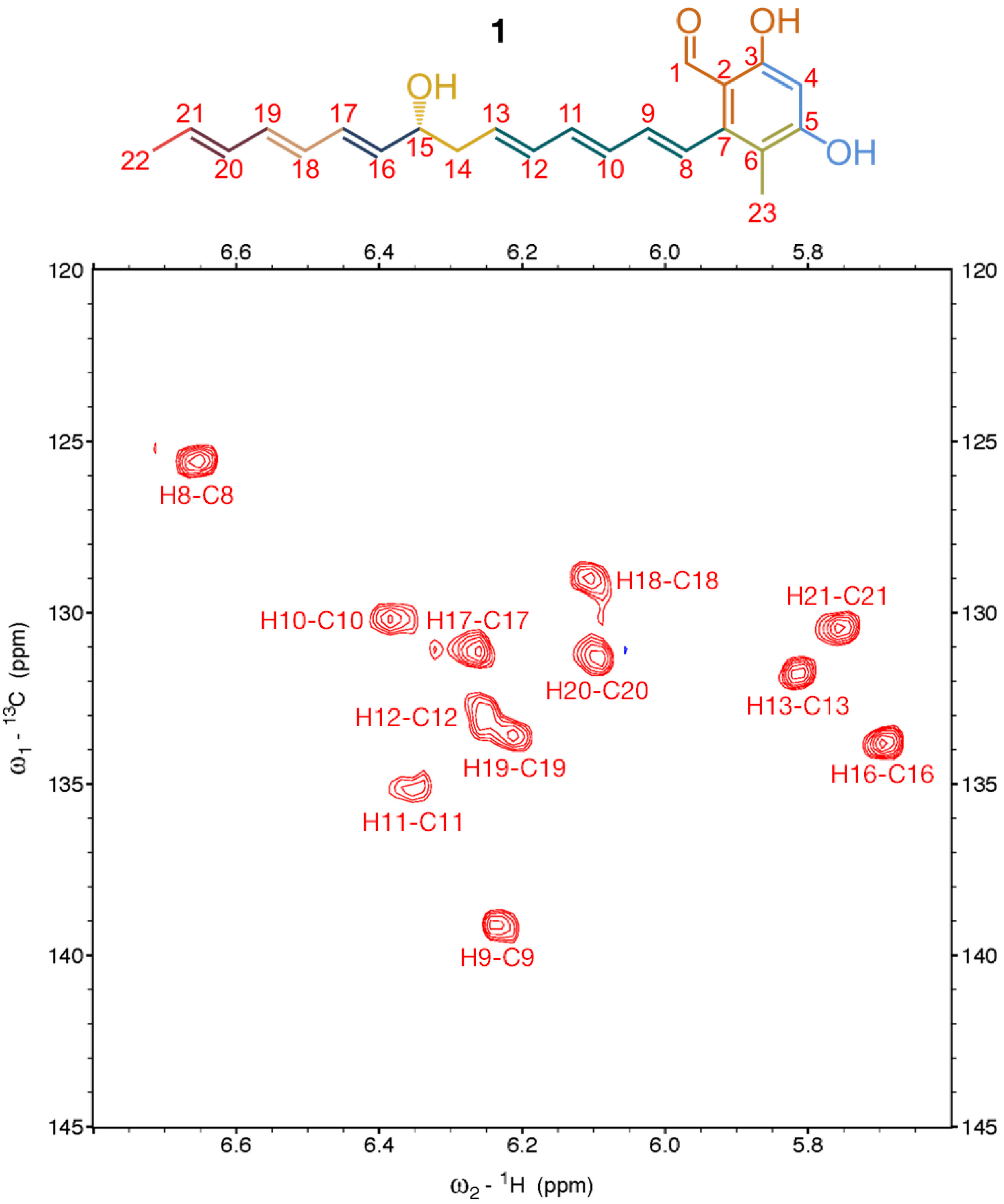

**Figure S44**. $^{1}$H-$^{13}$C HSQC spectrum (positive signals in red; negative signals in blue; x-axis: δH, 6.8-5.6 ppm; y-axis: δC, 145.0-120.0 ppm) of **1** in $CDCl_3$. Spectrum acquired on a 900 MHz Bruker AVANCE II spectrometer.

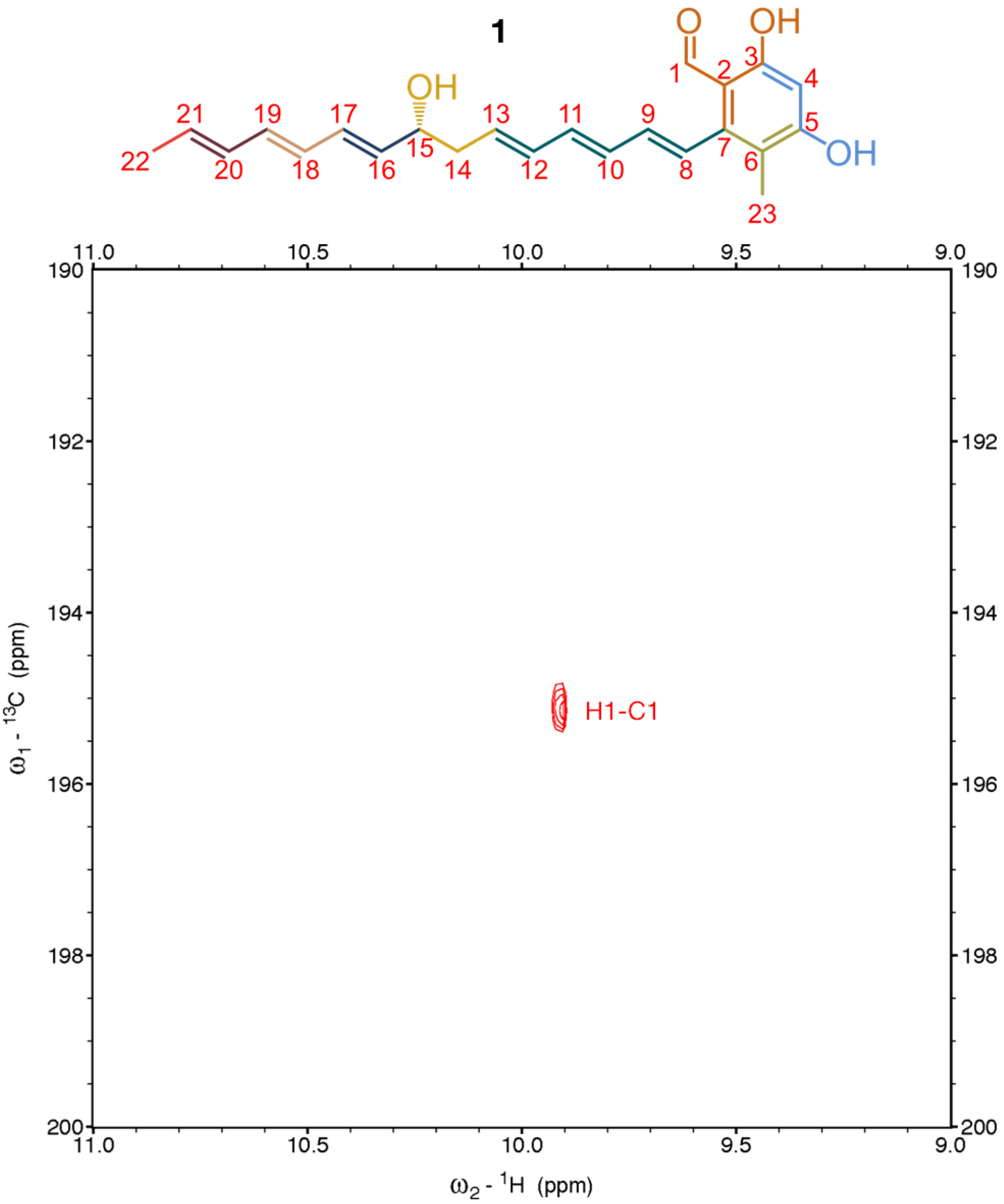

**Figure S45**. $^{1}H$-$^{13}C$ HSQC spectrum (positive signals in red; negative signals in blue; x-axis: δH, 11.0-9.0 ppm; y-axis: δC, 200.0-190.0 ppm) of **1** in $CDCl_3$. Spectrum acquired on a 900 MHz Bruker AVANCE II spectrometer.

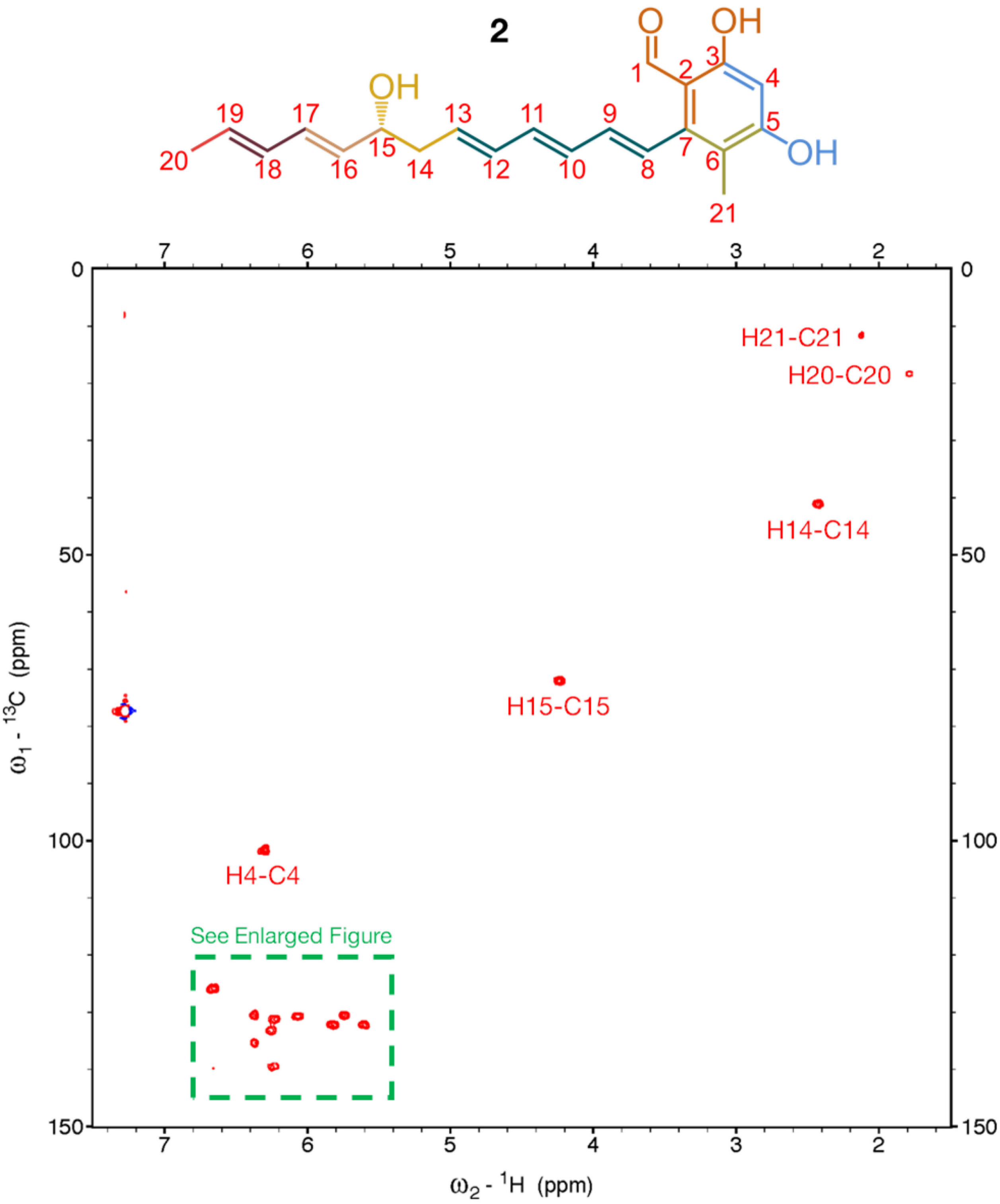

**Figure S46**. $^1H$-$^{13}C$ HSQC spectrum (positive signals in red; negative signals in blue; x-axis: δH, 7.5-1.5 ppm; y-axis: δC, 150.0-0.0 ppm) of **2** in $CDCl_3$. Spectrum acquired on a 500 MHz Bruker AVANCE spectrometer.

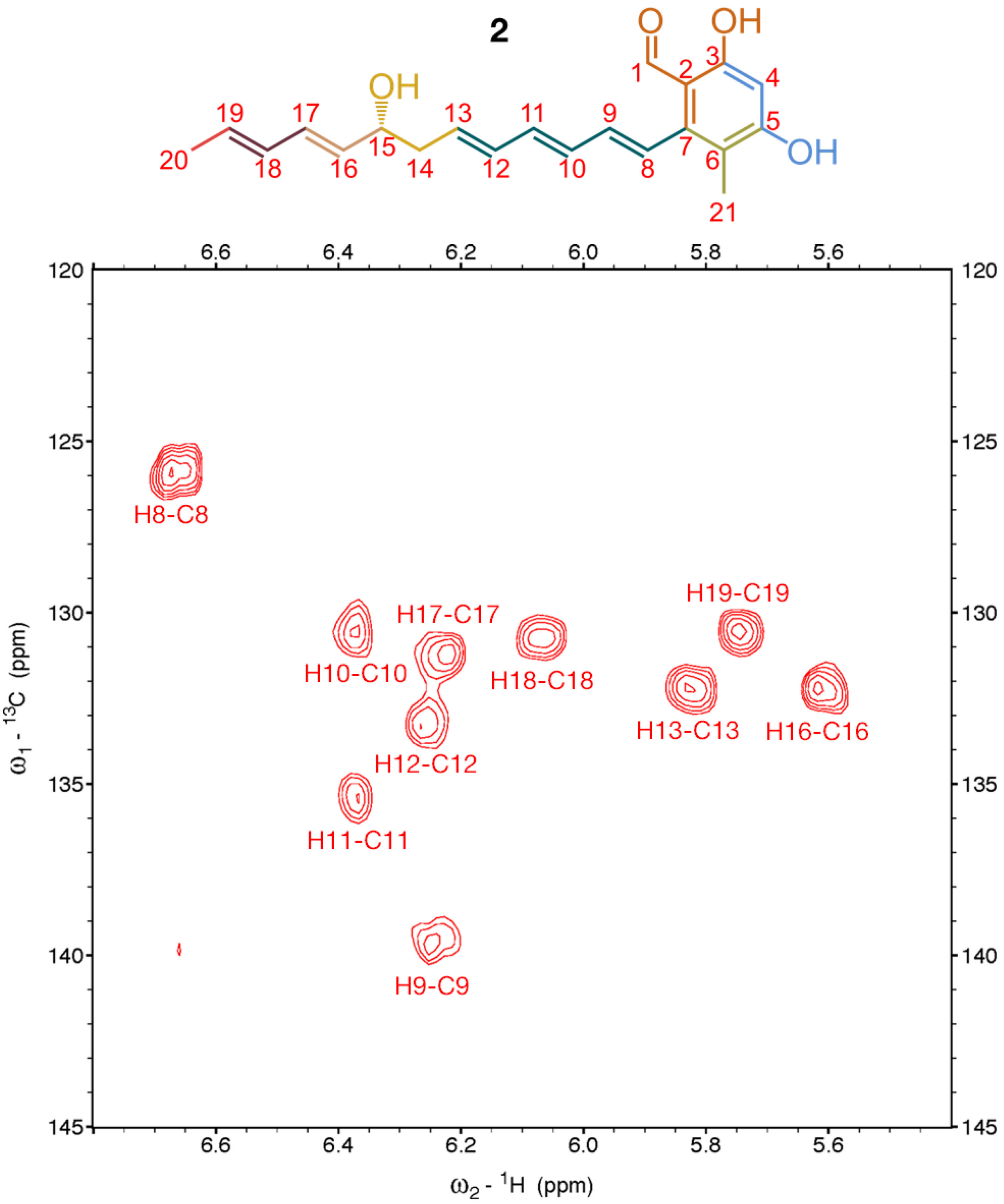

**Figure S47**. $^{1}H$-$^{13}C$ HSQC spectrum (positive signals in red; negative signals in blue; x-axis: δH, 6.8-5.4 ppm; y-axis: δC, 145.0-120.0 ppm) of **2** in $CDCl_3$. Spectrum acquired on a 500 MHz Bruker AVANCE spectrometer.

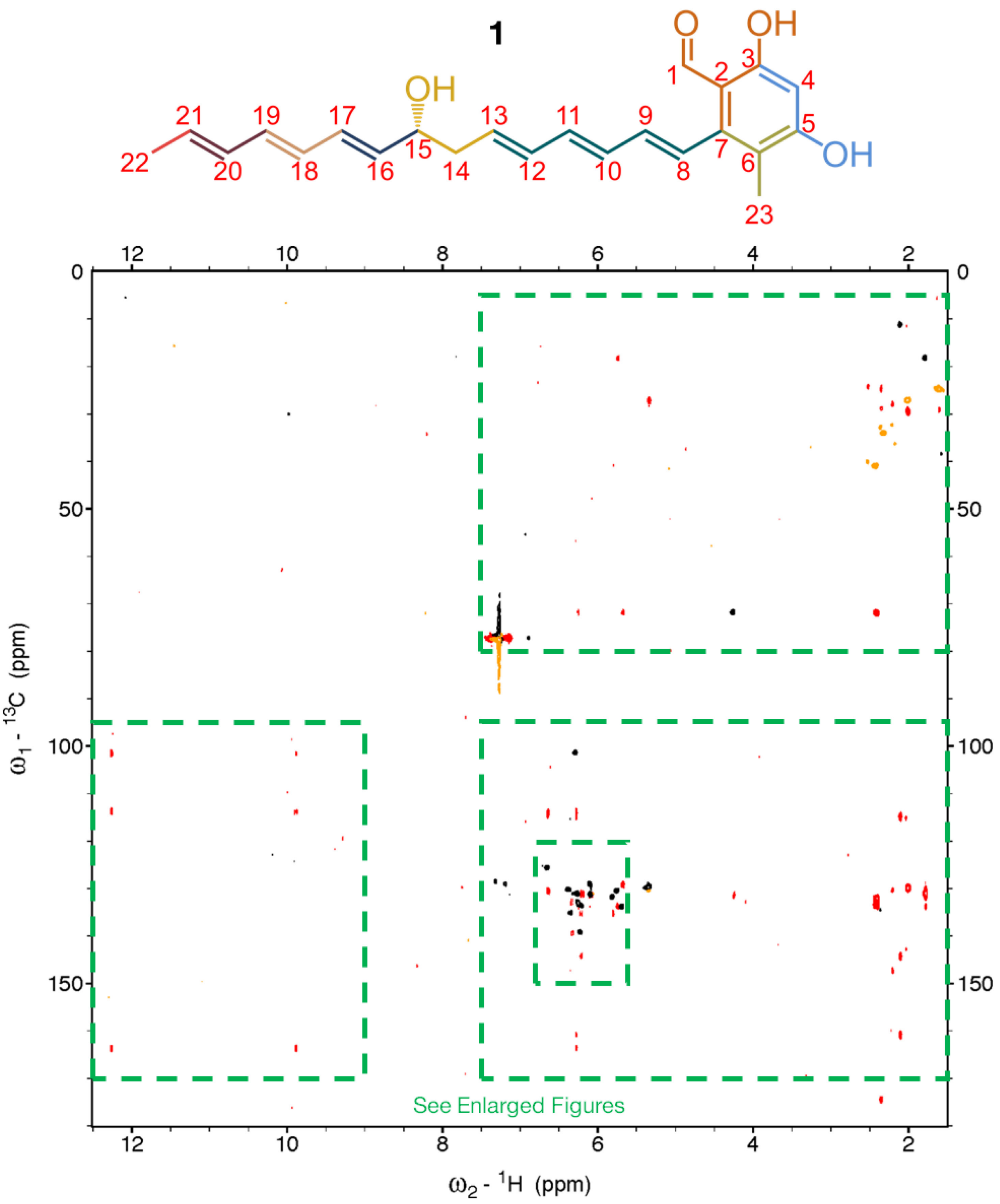

**Figure S48**. $^{1}H$-$^{13}C$ HMBC (HSQC overlaid in black for positive signals and orange for negative signals) spectrum (positive signals in red; negative signals in blue; x-axis: δH, 12.5-1.5 ppm; y-axis: δC, 180.0-0.0 ppm) of **1** in $CDCl_3$. Spectrum acquired on a 900 MHz Bruker AVANCE II spectrometer.

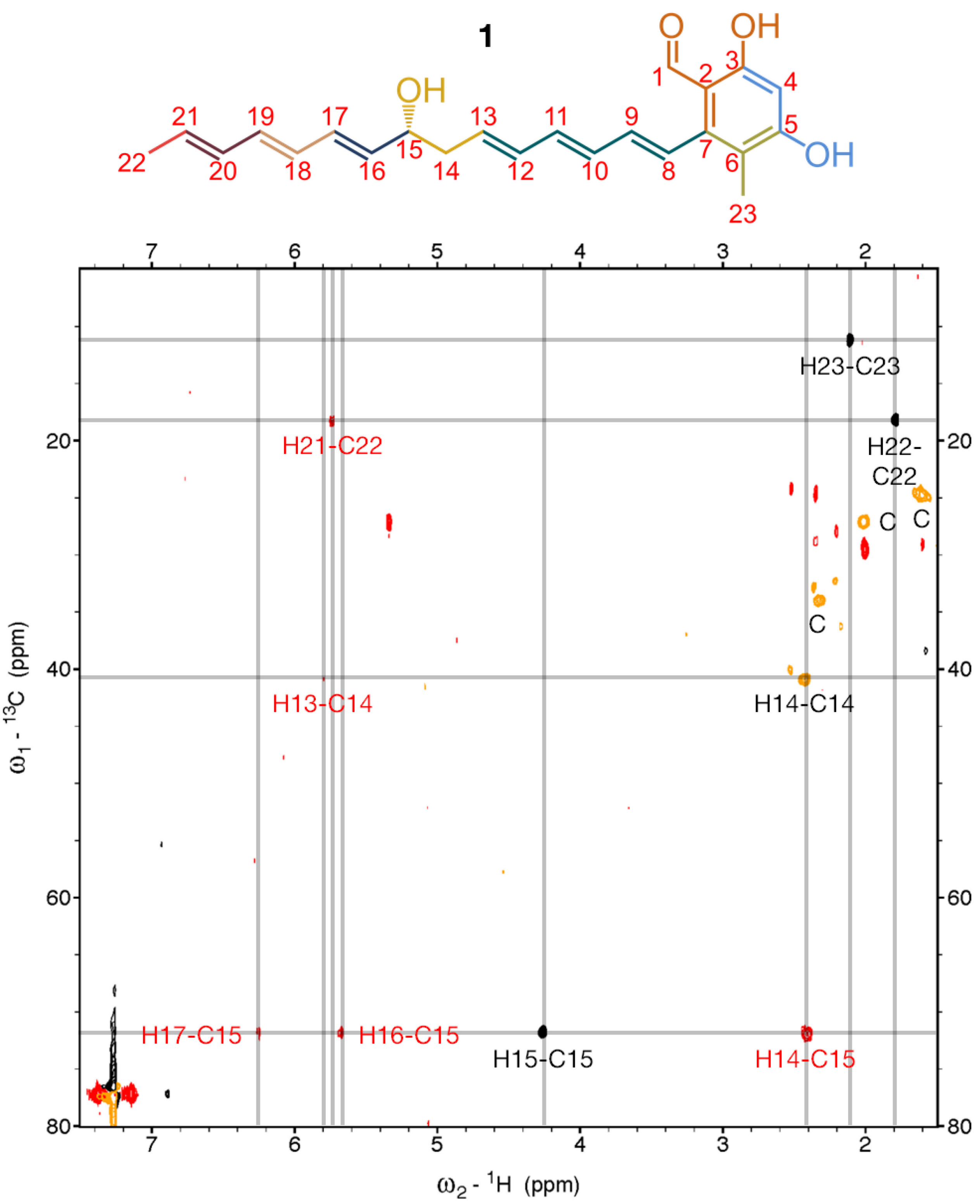

**Figure S49**. $^{1}$H-$^{13}$C HMBC (HSQC overlaid in black for positive signals and orange for negative signals) spectrum (positive signals in red; negative signals in blue; x-axis: δH, 7.5-1.5 ppm; y-axis: δC, 80.0-5.0 ppm) of **1** in $CDCl_3$. Key: C, contaminant. Spectrum acquired on a 900 MHz Bruker AVANCE II spectrometer.

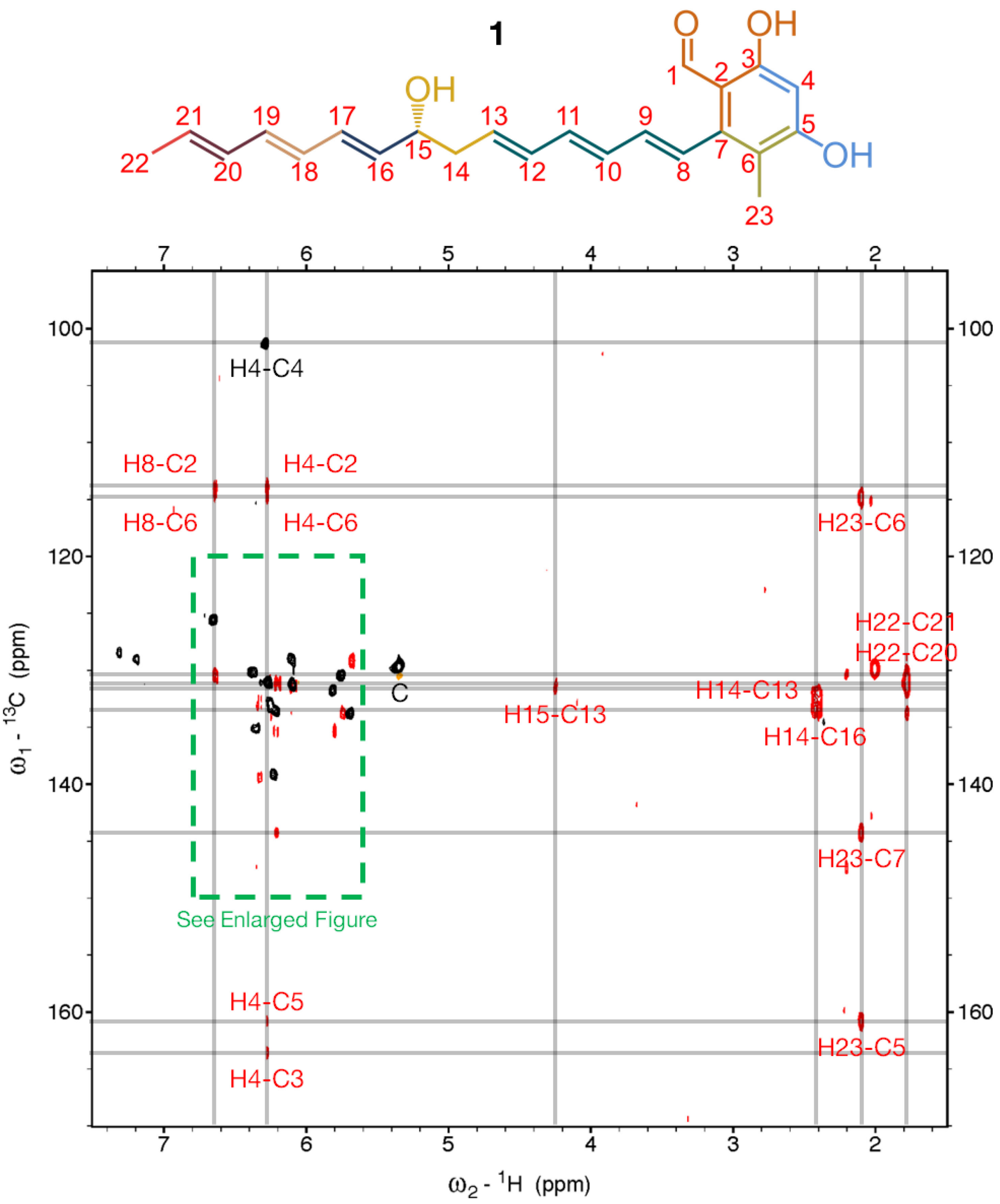

**Figure S50**. $^{1}H$-$^{13}C$ HMBC (HSQC overlaid in black for positive signals and orange for negative signals) spectrum (positive signals in red; negative signals in blue; x-axis: δH, 7.5-1.5 ppm; y-axis: δC, 170.0-95.0 ppm) of **1** in $CDCl_3$. Key: C, contaminant. Spectrum acquired on a 900 MHz Bruker AVANCE II spectrometer.

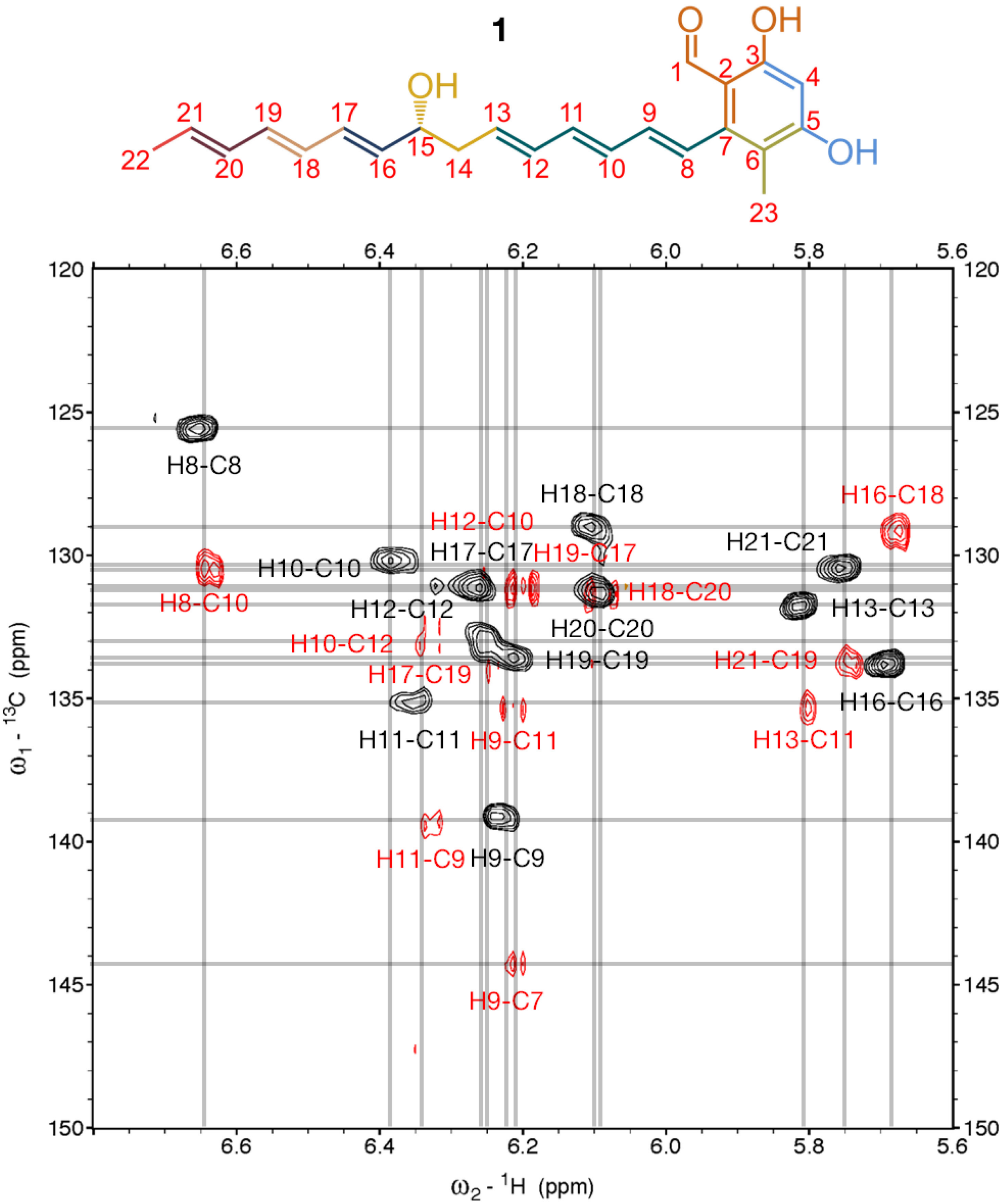

**Figure S51**. $^1$H-$^{13}$C HMBC (HSQC overlaid in black for positive signals and orange for negative signals) spectrum (positive signals in red; negative signals in blue; x-axis: δH, 6.8-5.6 ppm; y-axis: δC, 150.0-120.0 ppm) of **1** in $CDCl_3$. Spectrum acquired on a 900 MHz Bruker AVANCE II spectrometer.

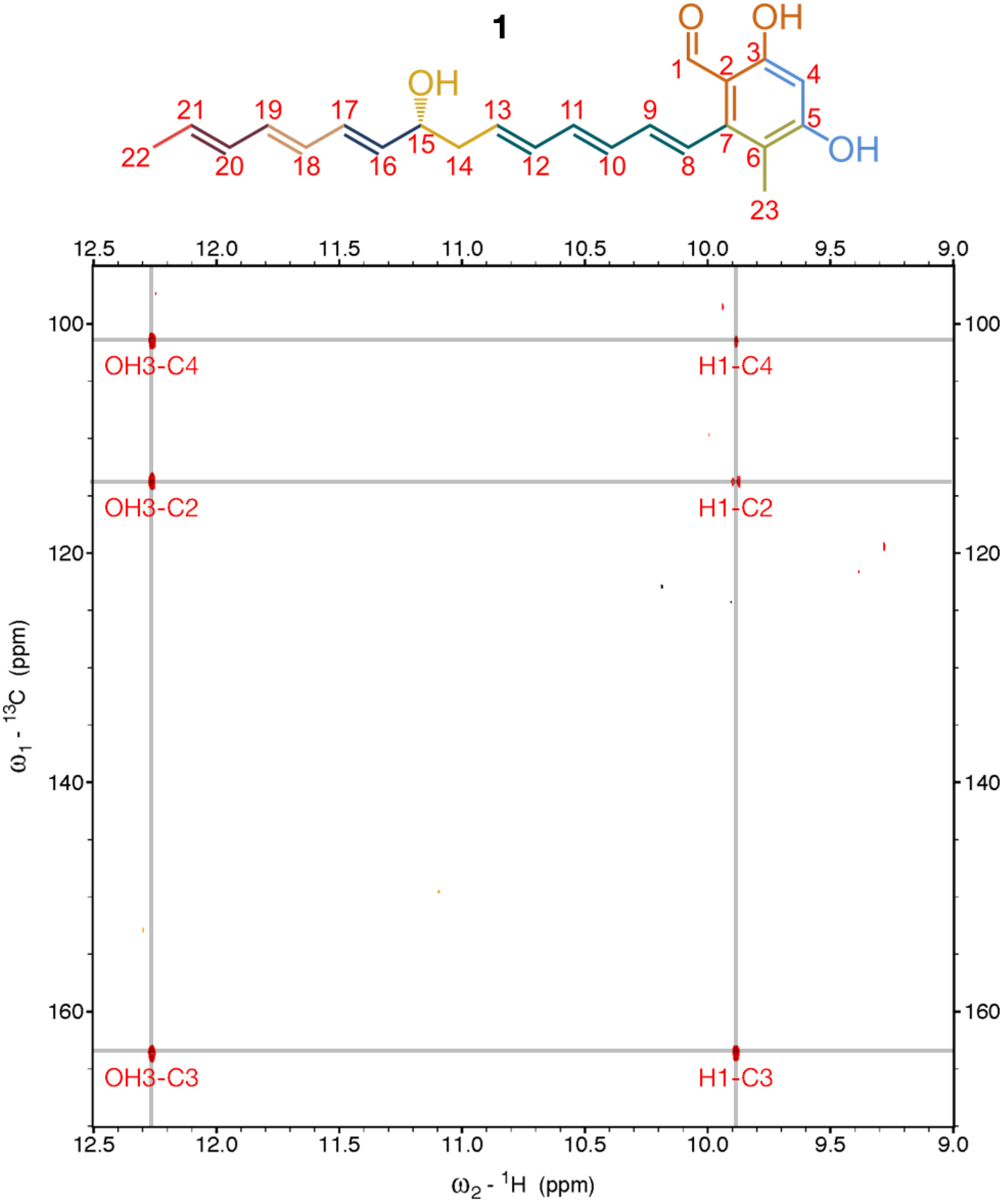

**Figure S52**. $^{1}H$-$^{13}C$ HMBC (HSQC overlaid in black for positive signals and orange for negative signals) spectrum (positive signals in red; negative signals in blue; x-axis: δH, 12.5-9.0 ppm; y-axis: δC, 170.0-95.0 ppm) of **1** in $CDCl_3$. Spectrum acquired on a 900 MHz Bruker AVANCE II spectrometer.

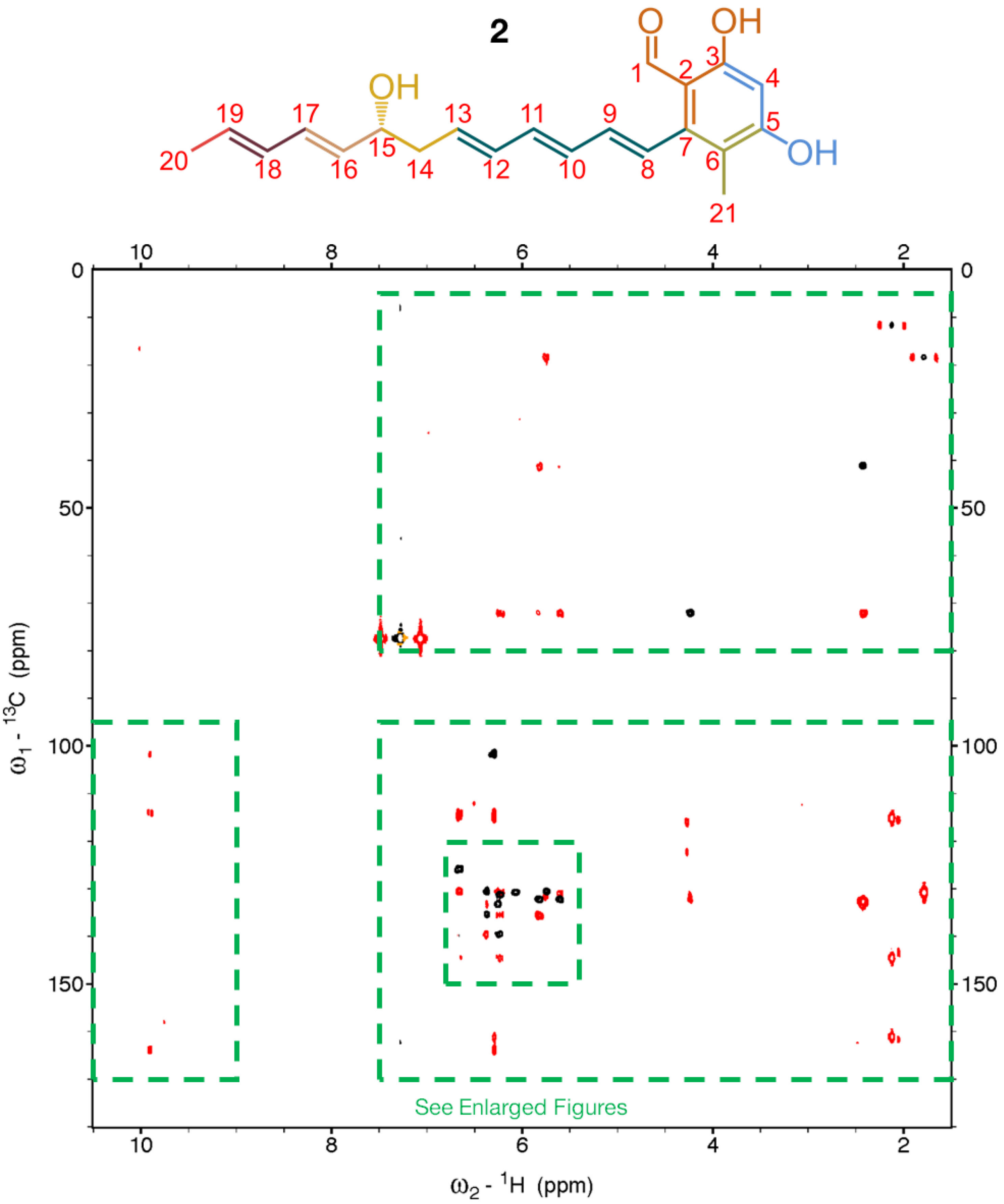

**Figure S53**. $^1$H-$^{13}$C HMBC (HSQC overlaid in black for positive signals and orange for negative signals) spectrum (positive signals in red; negative signals in blue; x-axis: δH, 10.5-1.5 ppm; y-axis: δC, 180.0-0.0 ppm) of **2** in $CDCl_3$. Spectrum acquired on a 500 MHz Bruker AVANCE spectrometer.

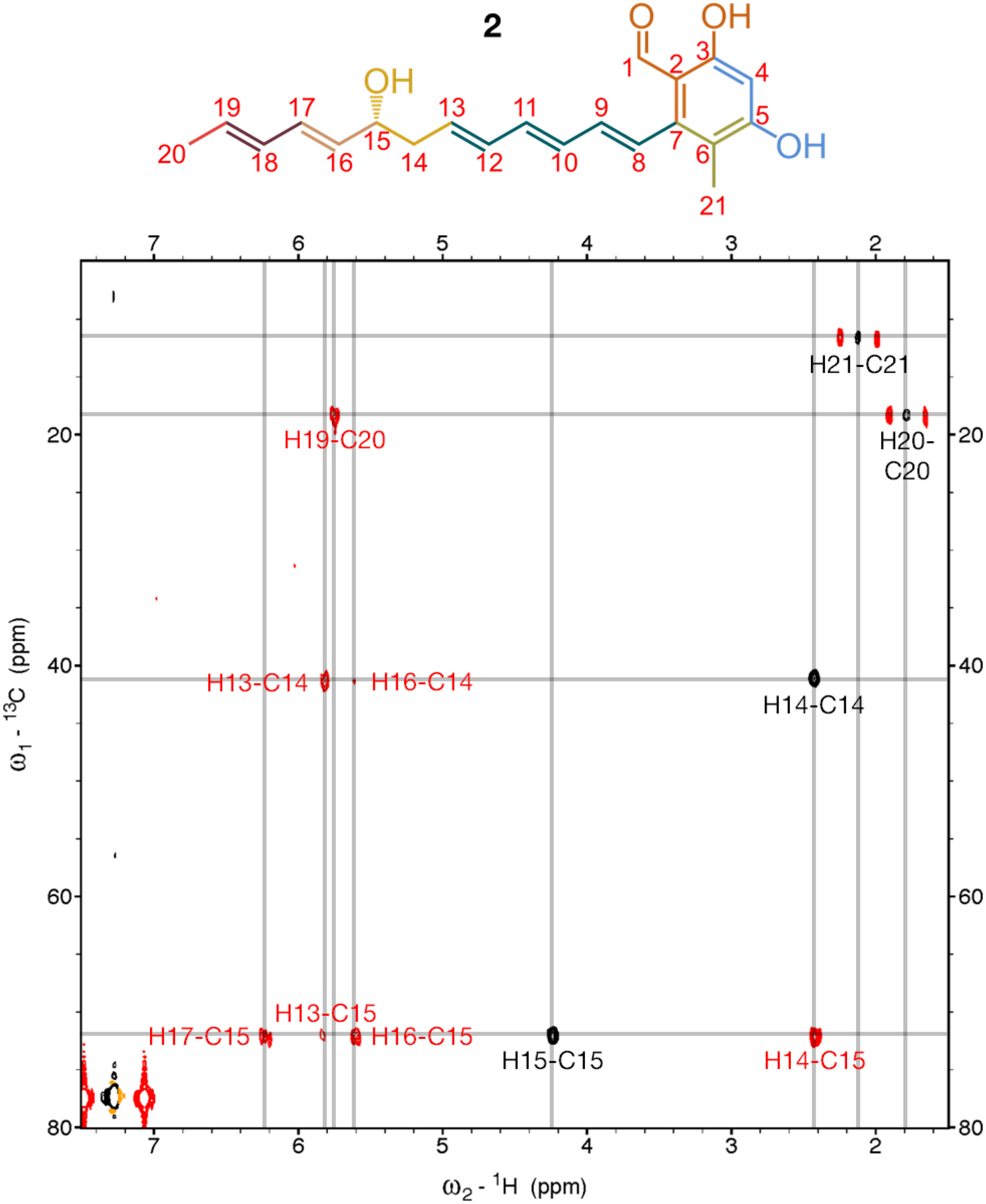

**Figure S54**. ¹H-¹³C HMBC (HSQC overlaid in black for positive signals and orange for negative signals) spectrum (positive signals in red; negative signals in blue; x-axis: δH, 7.5-1.5 ppm; y-axis: δC, 80.0-5.0 ppm) of **2** in $CDCl_3$. Spectrum acquired on a 500 MHz Bruker AVANCE spectrometer.

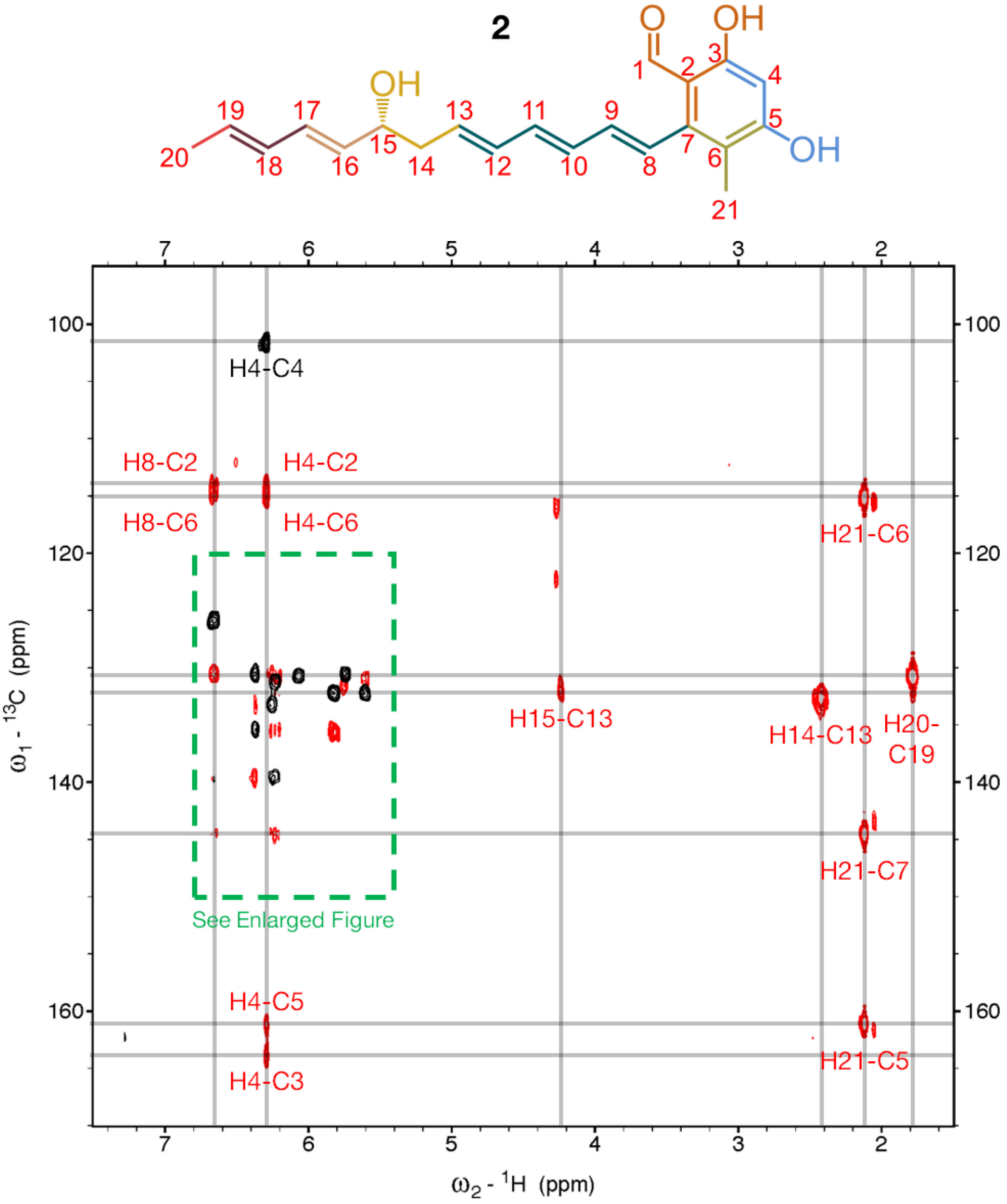

**Figure S55**. $^{1}$H-$^{13}$C HMBC (HSQC overlaid in black for positive signals and orange for negative signals) spectrum (positive signals in red; negative signals in blue; x-axis: δH, 7.5-1.5 ppm; y-axis: δC, 170.0-95.0 ppm) of **2** in $CDCl_3$. Spectrum acquired on a 500 MHz Bruker AVANCE spectrometer.

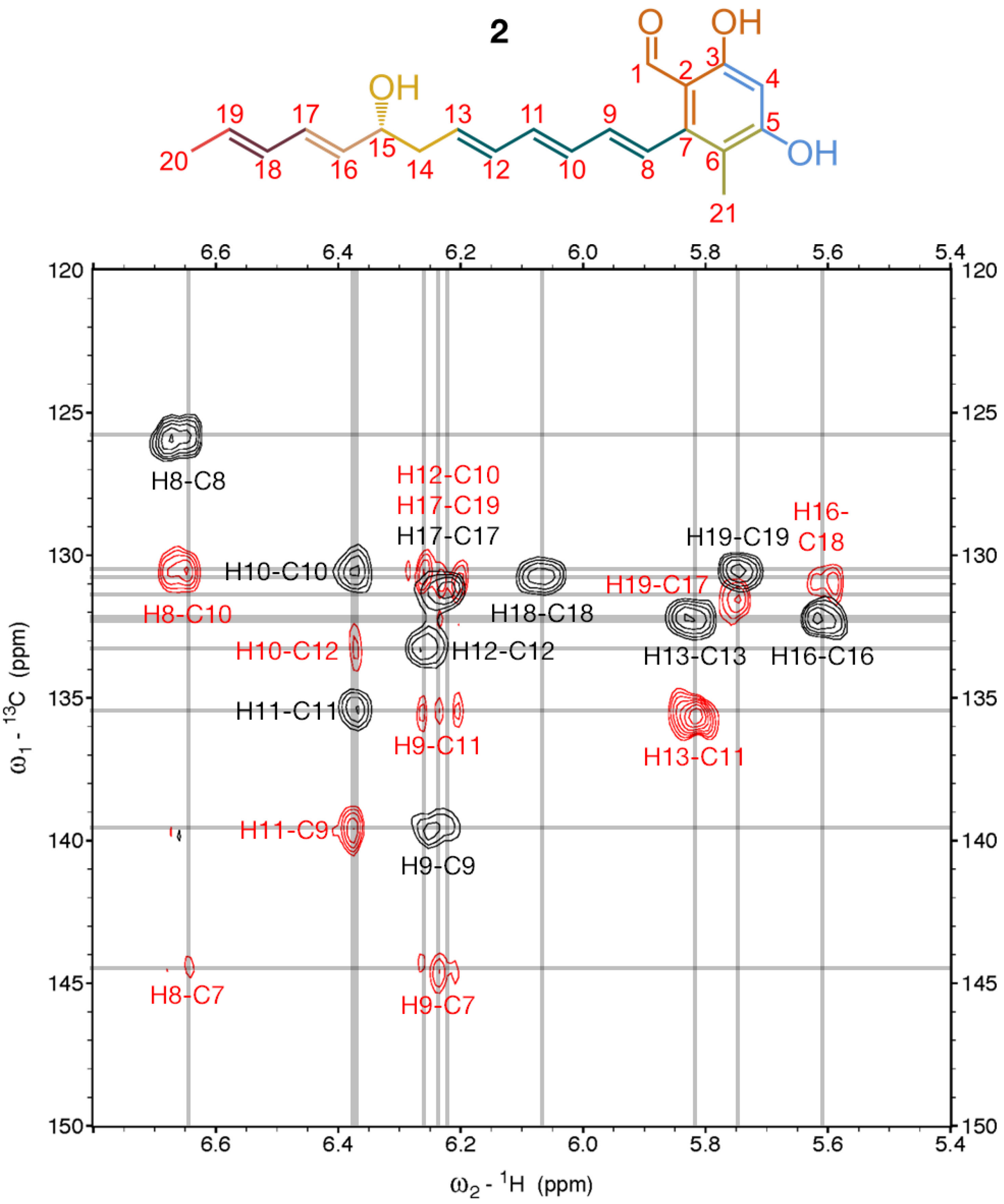

**Figure S56**. $^1H$-$^{13}C$ HMBC (HSQC overlaid in black for positive signals and orange for negative signals) spectrum (positive signals in red; negative signals in blue; x-axis: δH, 6.8-5.4 ppm; y-axis: δC, 150.0-120.0 ppm) of **2** in $CDCl_3$. Spectrum acquired on a 500 MHz Bruker AVANCE spectrometer.

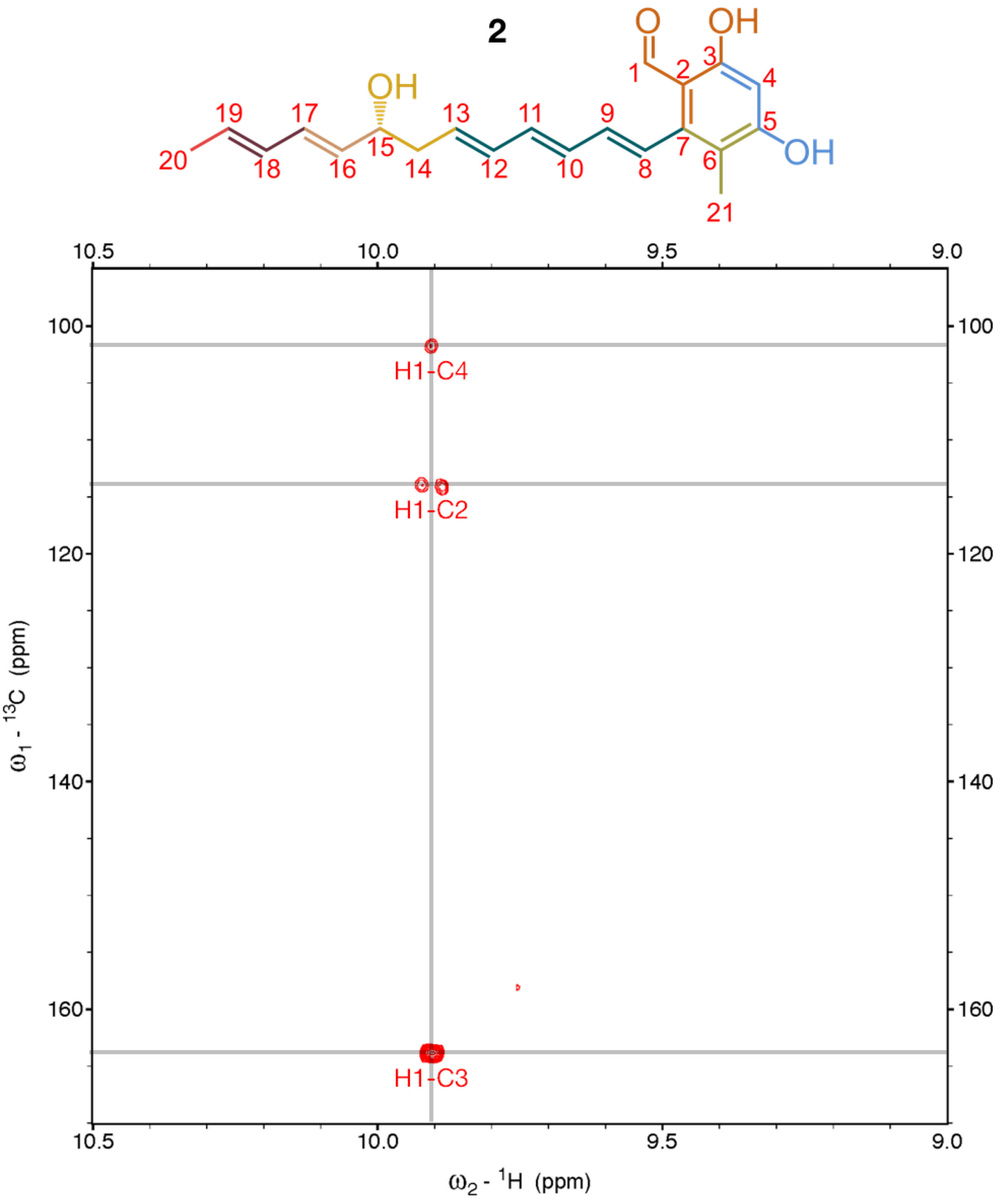

**Figure S57**. $^{1}H$-$^{13}C$ HMBC (HSQC overlaid in black for positive signals and orange for negative signals) spectrum (positive signals in red; negative signals in blue; x-axis: δH, 10.5-9.0 ppm; y-axis: δC, 170.0-95.0 ppm) of **2** in $CDCl_3$. Spectrum acquired on a 500 MHz Bruker AVANCE spectrometer.

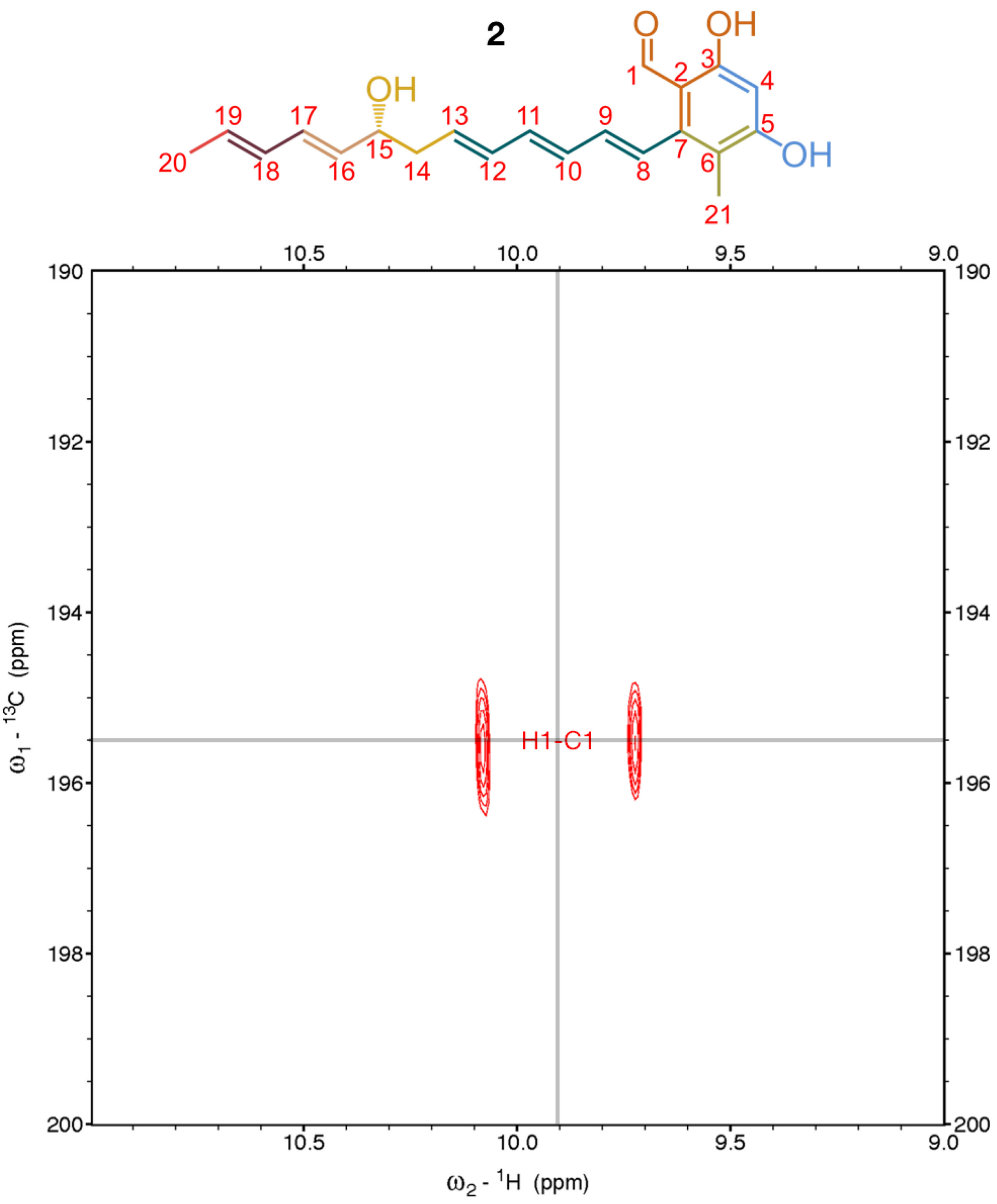

**Figure S58**. [1]H-[13]C HMBC spectrum (positive signals in red; negative signals in blue; x-axis: δH, 11.0-9.0 ppm; y-axis: δC, 200.0-190.0 ppm) of **2** in $CDCl_3$. Spectrum acquired on a 500 MHz Bruker AVANCE spectrometer.

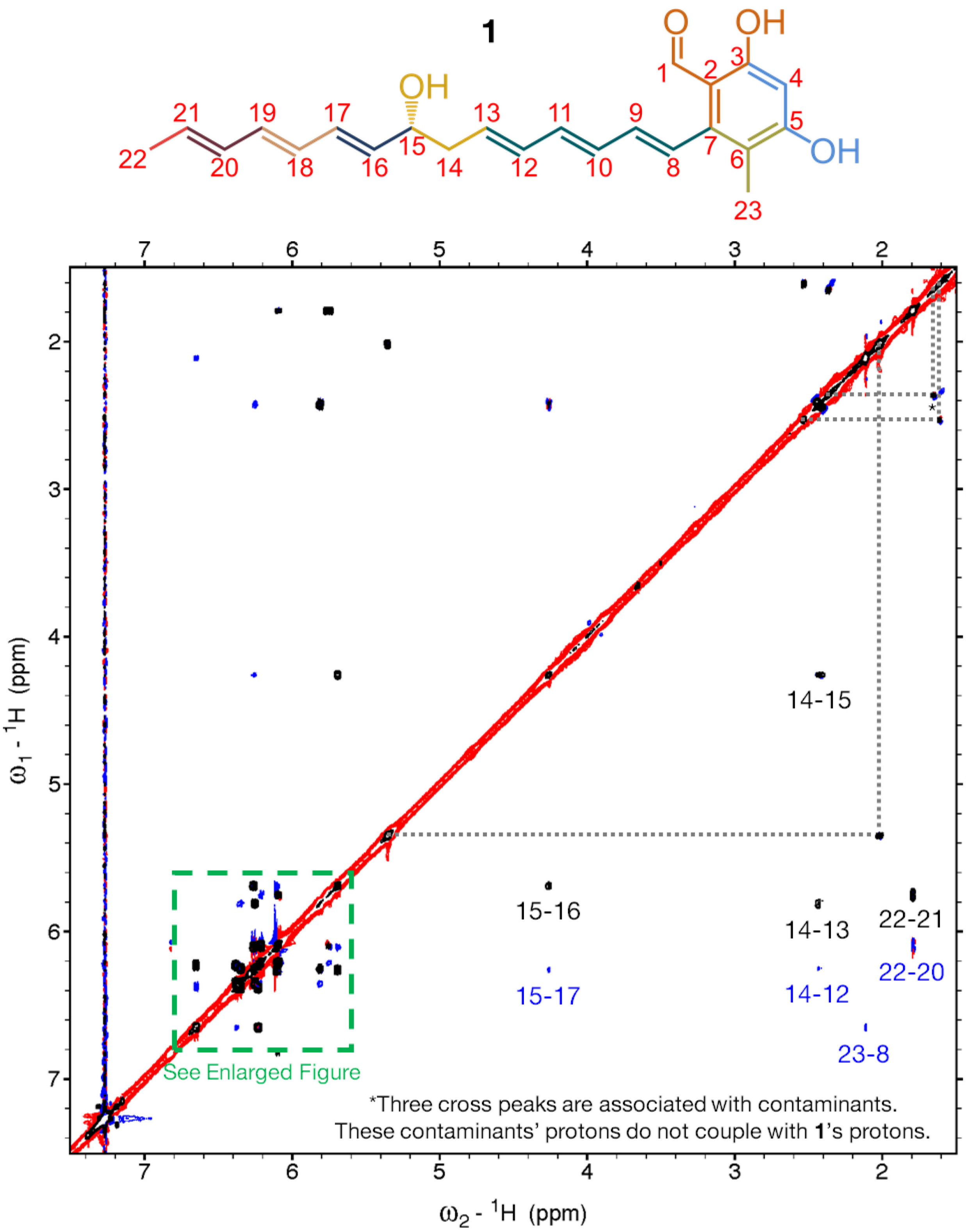

**Figure S59**. [1]H-[1]H ROESY (COSY overlaid in black for positive signals and orange for negative signals) spectrum (positive signals in red; negative signals in blue; x-axis: δH, 7.5-1.5 ppm; y-axis: δH, 7.5-1.5 ppm) of **1** in $CDCl_3$. Spectrum acquired on a 900 MHz Bruker AVANCE II spectrometer.

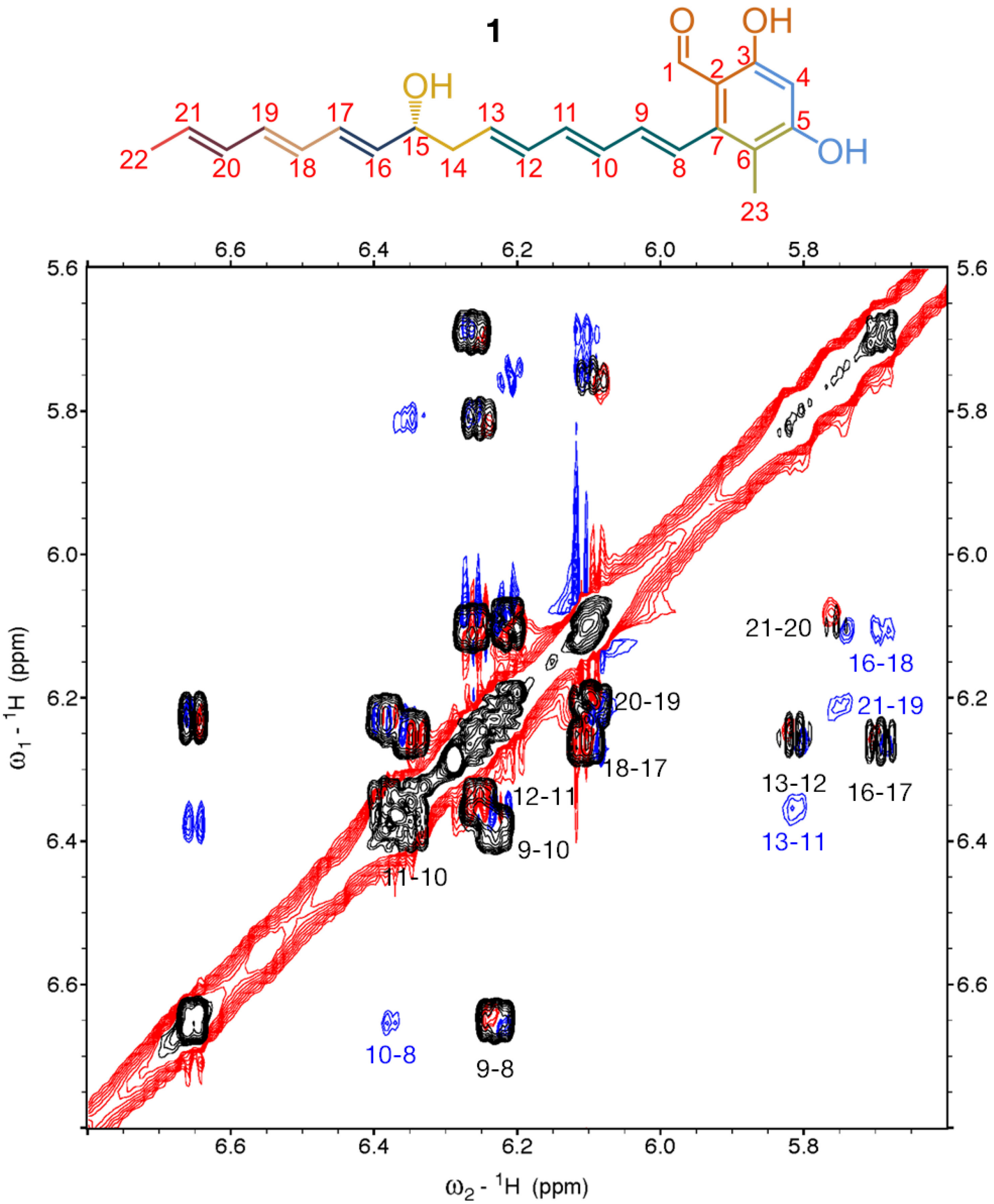

**Figure S60**. $^1$H-$^1$H ROESY (COSY overlaid in black for positive signals and orange for negative signals) spectrum (positive signals in red; negative signals in blue; x-axis: δH, 6.8-5.6 ppm; y-axis: δH, 6.8-5.6 ppm) of **1** in $CDCl_3$. Spectrum acquired on a 900 MHz Bruker AVANCE II spectrometer.

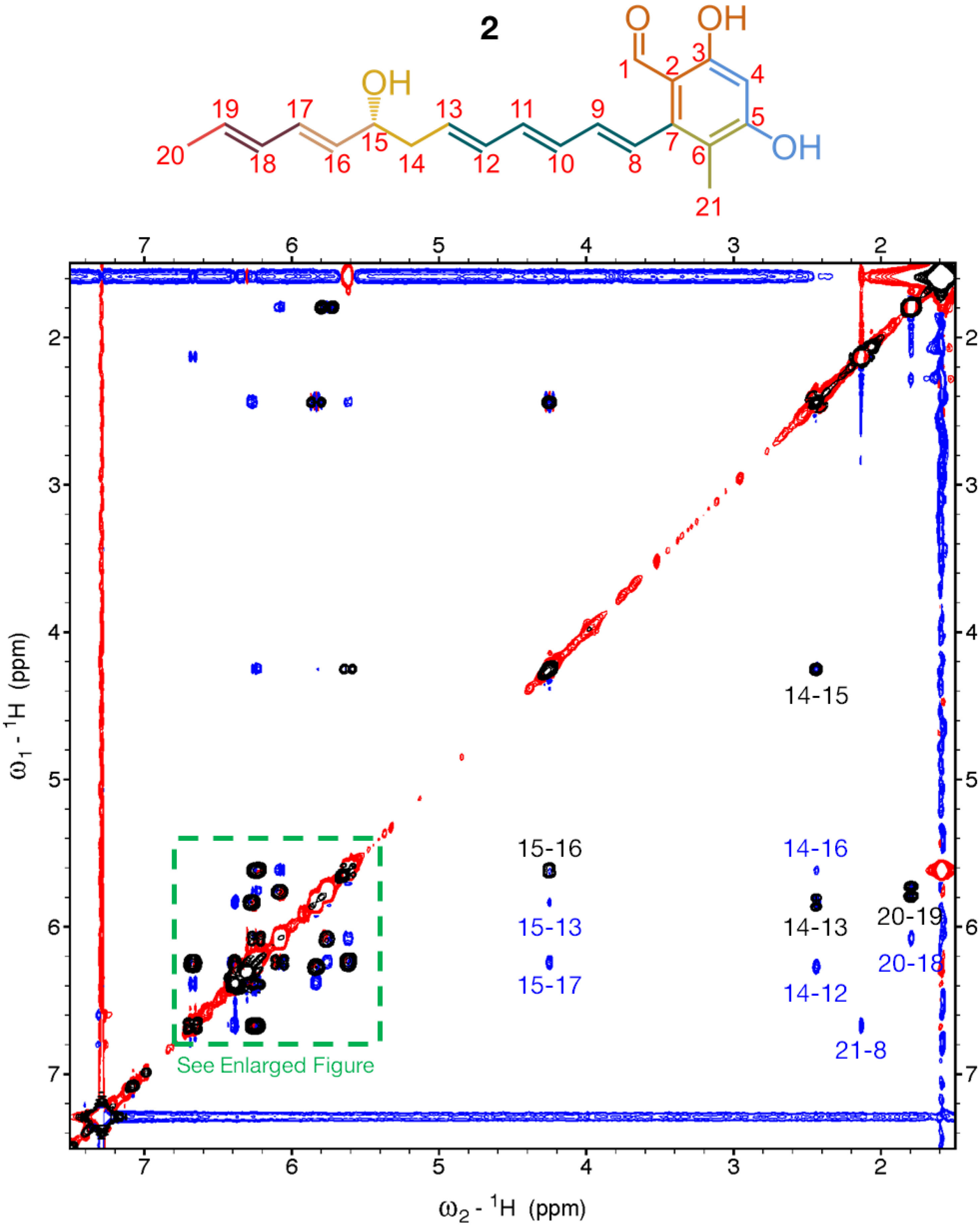

**Figure S61**. $^1$H-$^1$H NOESY (COSY overlaid in black for positive signals and orange for negative signals) spectrum (positive signals in red; negative signals in blue; x-axis: δH, 7.5-1.5 ppm; y-axis: δH, 7.5-1.5 ppm) of **2** in $CDCl_3$. Spectrum acquired on a 500 MHz Bruker AVANCE spectrometer.

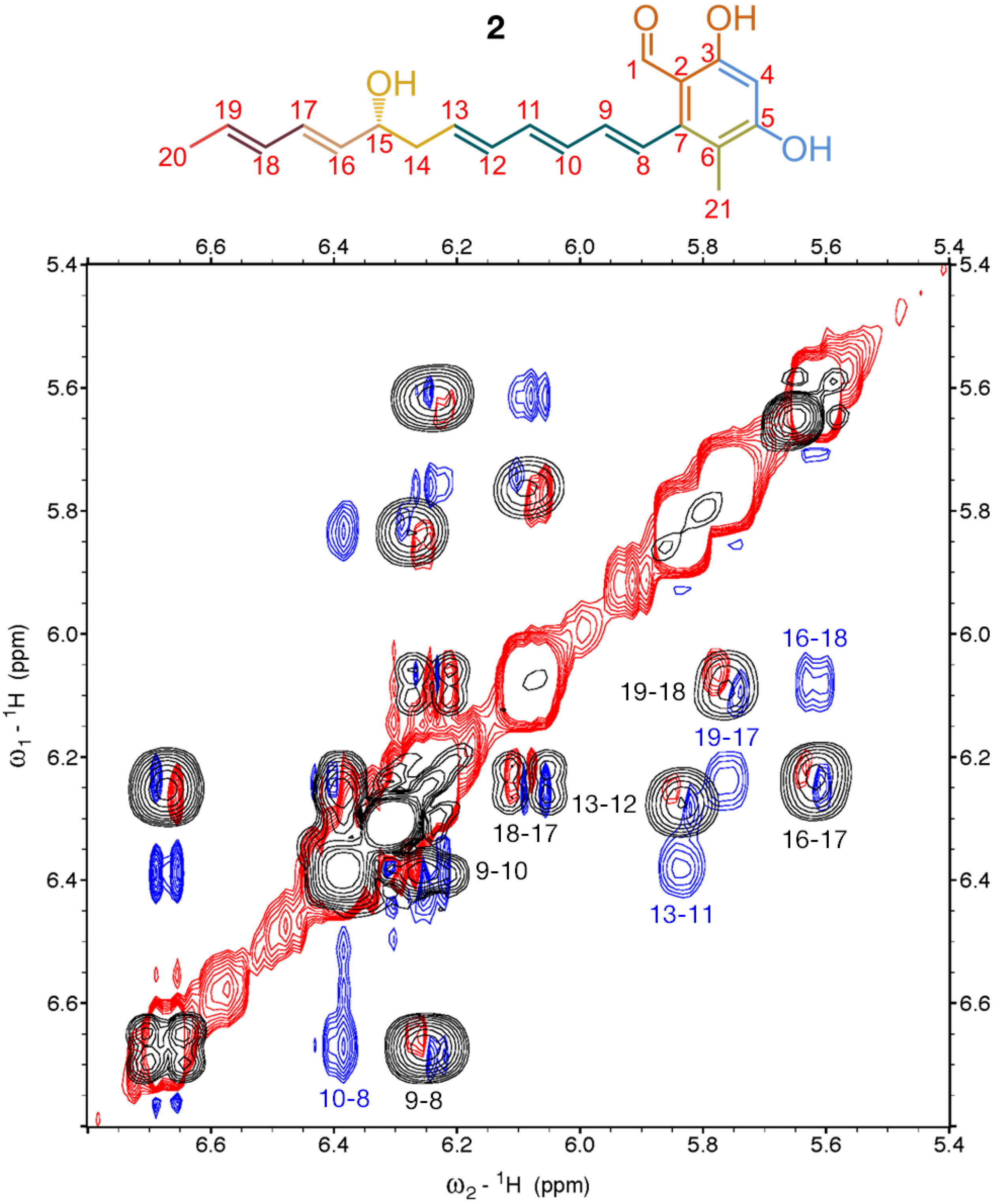

**Figure S62**. $^1H$-$^1H$ NOESY (COSY overlaid in black for positive signals and orange for negative signals) spectrum (positive signals in red; negative signals in blue; x-axis: δH, 6.8-5.4 ppm; y-axis: δH, 6.8-5.4 ppm) of **2** in $CDCl_3$. Spectrum acquired on a 500 MHz Bruker AVANCE spectrometer.

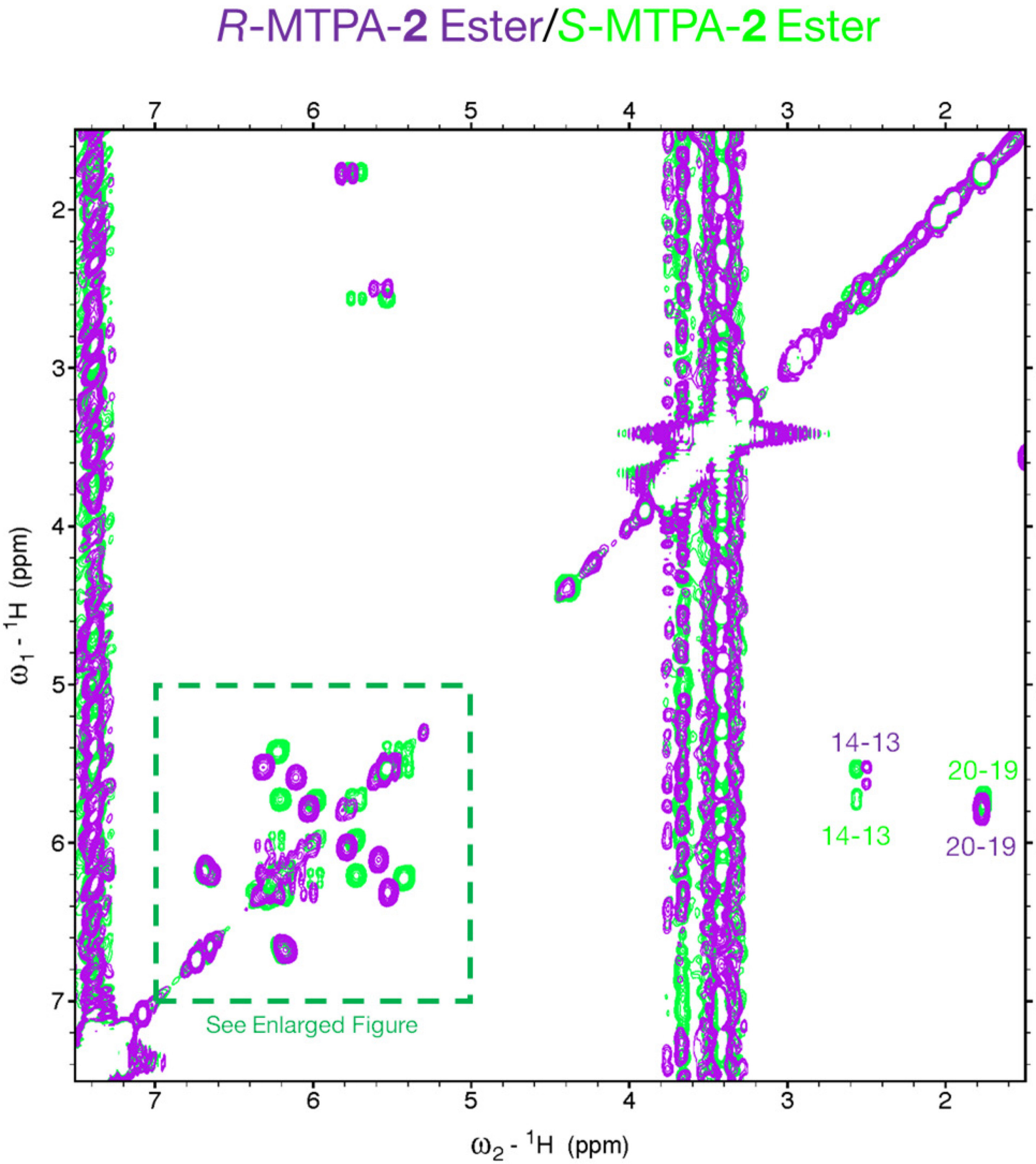

**Figure S63**. Overlaid [1]H-[1]H COSY spectra (x-axis: δH, 7.5-1.5 ppm; y-axis: δH, 7.5-1.5 ppm) of *R*-MTPA-**2** ester (positive signals in purple; negative signals in blue) and *S*-MTPA-**2** ester (positive signals in green; negative signals in orange) in $CDCl_3$. Spectrum acquired on a 500 MHz Bruker AVANCE spectrometer.

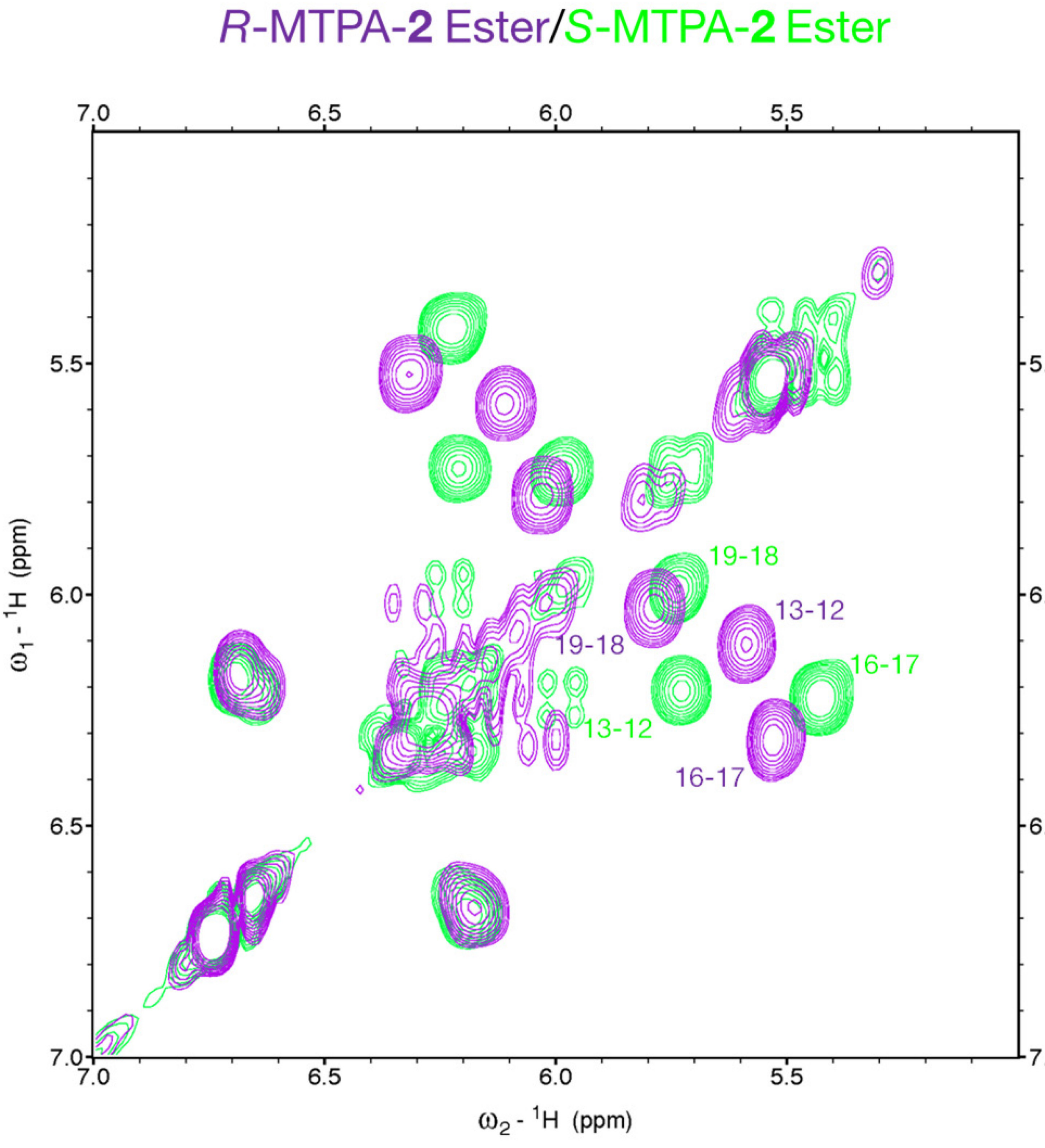

**Figure S64**. Overlaid [1]H-[1]H COSY spectra (x-axis: δH, 7.0-5.0 ppm; y-axis: δH, 7.0-5.0 ppm) of *R*-MTPA-**2** ester (positive signals in purple; negative signals in blue) and *S*-MTPA-**2** ester (positive signals in green; negative signals in orange) in $CDCl_3$. Spectrum acquired on a 500 MHz Bruker AVANCE spectrometer.

**2**

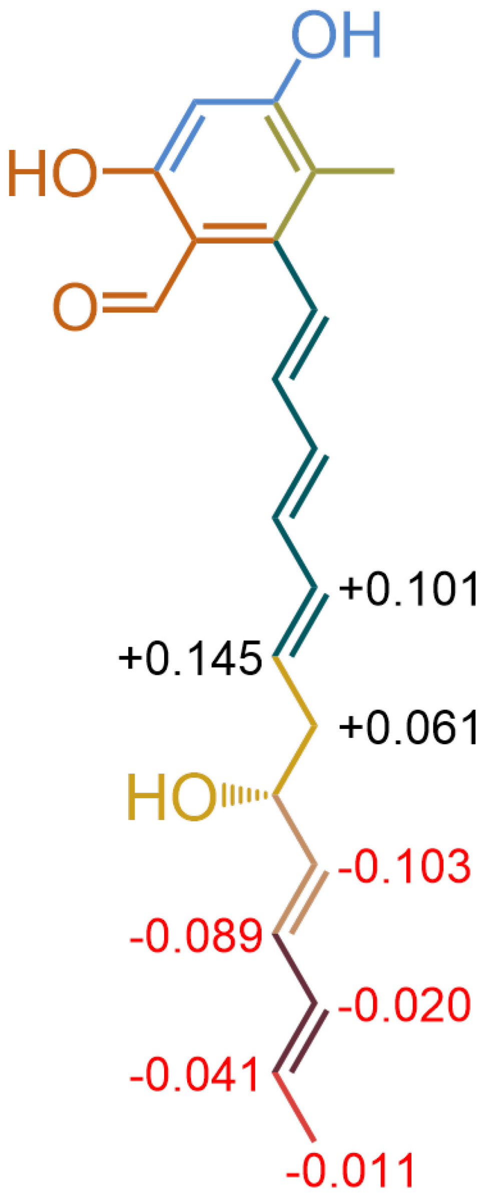

**Figure S65**. ΔδSR values (in ppm) for the protons that flank the C-15 hydroxyl substituent in **2**.

| Domain(s) | KR Motif | Predicted | Found |
|---|---|---|---|
| KR1/DH1 | GVVHCAGV**LDD** | D(OH) → *E*-DB | *E*-DB |
| KR2/DH2 | GVVHCAGV**LDD** | D(OH) → *E*-DB | *E*-DB |
| KR4 | GVLHCAGS**VST** | L(OH) | L(OH) |
| KR5/DH5 | VLYHGAGQ**LRD** | D(OH) → *E*-DB | *E*-DB |

**Figure S66**. Bioinformatic inspection of the KR domains[11] of the NOCAP synthase predicts that 1) all of **1** and **2**'s double bonds have *trans* stereoconfigurations and 2) the absolute configuration at C-15 is *R*.

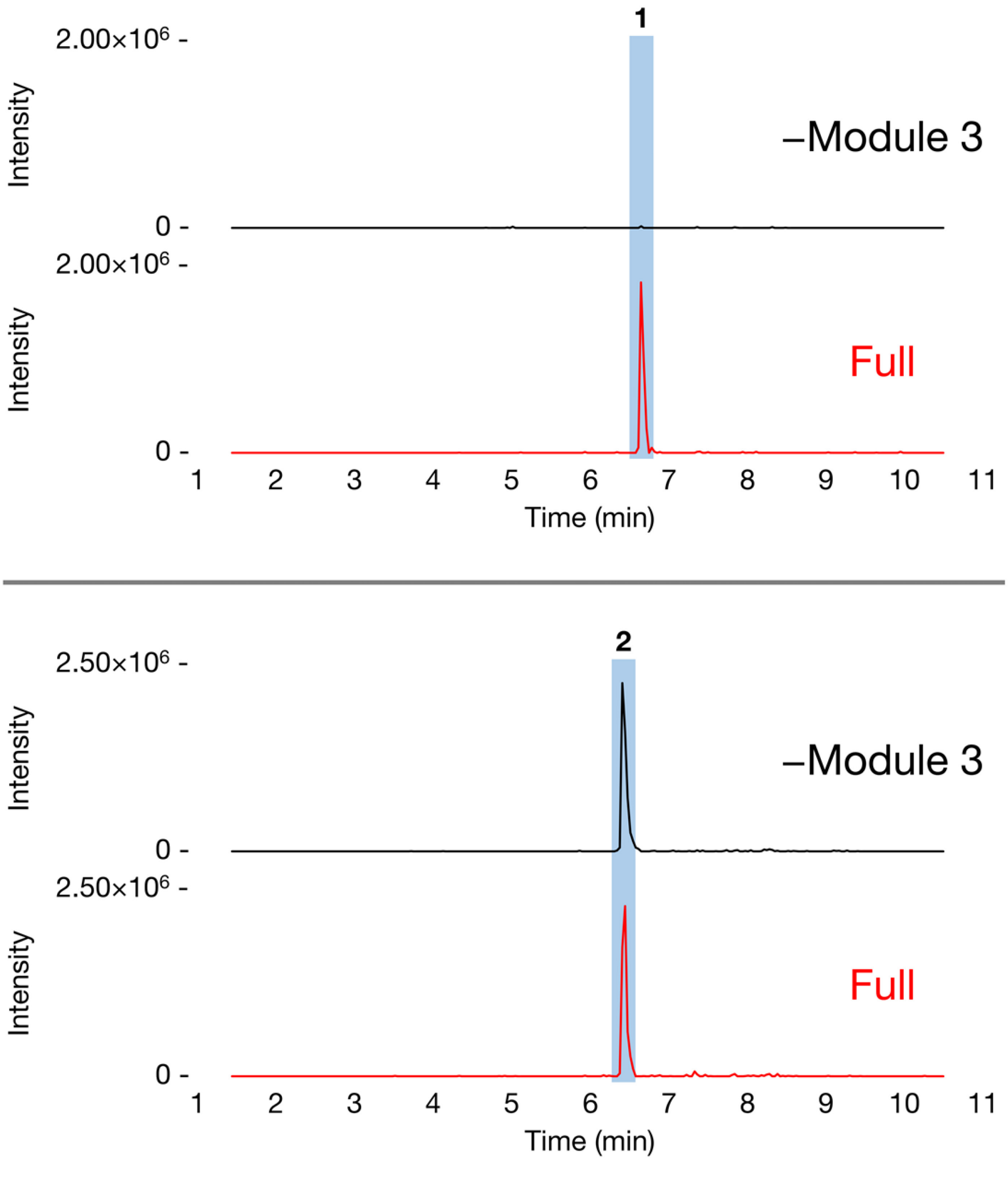

**Figure S67**. EICs (ESI-) of **1** and **2** produced in either *E. coli* BAP1[pCK-KPY222/pCK-KPY259/pCK-KPY178] (full system) or *E. coli* BAP1[pCK-KPY102/pCK-KPY259/pCK-KPY178] (module 3 is omitted). Data acquired on a Waters SQ Detector 2 LC-MS system and are representative of at least three independent experimental replicates.

**Table S1**. *Nocardia* strains that contain the NOCAP synthase.

| *Nocardia* Species | DSM Strain No. | Genbank Accession No. (Main PKS Contig) | Reference |
|---|---|---|---|
| *N. abscessus* | 44432, 44557 | BAFP01000080.1 | 12 |
| *N. amamiensis* | 45066 | BDBA01000032.1 | 13 |
| *N. araoensis* | 44729 | BAFR01000107.1 | 14 |
| *N. arthritidis* | 44731 | BDBB01000017.1 | 15 |
| *N. asiatica* | 44668, 44700 | BAFS01000454.1 | 16 |
| *N. beijingensis* | 43474, 45494 | BDBC01000032.1 | 17 |
| *N. bhagyanarayanae* | 103495 | VFPG01000001.1 | 18 |
| *N. exalbida* | 44883, 44884 | BAFZ01000063.1 | 19 |
| *N. gamkensis* | 44956 | BDBM01000012.1 | 20 |
| *N. niwae* | 45340 | BDCK01000022.1 | 21 |
| *N. paucivorans* | 44386, 44543, 44563 | BAGE01000052.1 | 22 |
| *N. pneumoniae* | 44730 | BAGF01000151.1 | 14 |
| *N. puris* | 44599, 44699 | BDBW01000050.1 | 23 |

**Table S2**. NMR assignments for **1** in $CDCl_3$ (900 MHz) and **2** in $CDCl_3$ (500 MHz).

| **1** | | | **2** | | |
|---|---|---|---|---|---|
| Carbon | δC | δH | Carbon | δC | δH |
| **1** | 195.1 | 9.890 | **1** | 195.5 | 9.917 |
| **2** | 113.8 | - | **2** | 113.9 | - |
| **3** | 163.6 | 12.269 OH | **3** | 163.7 | - |
| **4** | 101.5 | 6.278 | **4** | 101.6 | 6.309 |
| **5** | 160.8 | - | **5** | 161.1 | - |
| **6** | 114.7 | - | **6** | 114.9 | - |
| **7** | 144.3 | - | **7** | 144.4 | - |
| **8** | 125.6 | 6.645 | **8** | 125.8 | 6.661 |
| **9** | 139.3 | 6.220 | **9** | 139.4 | 6.243 |
| **10** | 130.4 | 6.384 | **10** | 130.5 | 6.381 |
| **11** | 135.3 | 6.341 | **11** | 135.5 | 6.376 |
| **12** | 133.1 | 6.252 | **12** | 133.1 | 6.255 |
| **13** | 131.8 | 5.808 | **13** | 132.3 | 5.823 |
| **14** | 40.91 | 2.418 | **14** | 41.21 | 2.429 |
| **15** | 71.87 | 4.253 | **15** | 71.99 | 4.245 |
| **16** | 133.9 | 5.685 | **16** | 132.1 | 5.608 |
| **17** | 131.1 | 6.256 | **17** | 131.3 | 6.229 |
| **18** | 129.1 | 6.098 | **18** | 130.7 | 6.067 |
| **19** | 133.6 | 6.207 | **19** | 130.5 | 5.750 |
| **20** | 131.3 | 6.092 | **20** | 18.22 | 1.787 |
| **21** | 130.5 | 5.749 | **21** | 11.45 | 2.127 |
| **22** | 18.28 | 1.783 | | | |
| **23** | 11.17 | 2.103 | | | |

#### Supporting Information References
